# Social Experience and Motivated Cognition Shape the Representation of Concepts: A Behavioural and Functional Neuroimaging Study

**DOI:** 10.64898/2026.09.03.749162

**Authors:** Doina-Irina Giurgea, Veronica Diveica, Penny M. Pexman, Richard J. Binney

## Abstract

The ventrolateral anterior temporal lobe (vlATL), including the anterior inferior temporal and fusiform gyri, is the centre of a supramodal and category-general hub for semantic representation. Dorsolateral ATL regions, such as the temporal pole (TP) and anterior middle temporal gyrus (aMTG), have been associated with more specialised semantic function, including social concept processing, although the precise location and laterality of these effects vary across studies, prohibiting firm conclusions. The ‘socialness’ of concepts has been defined in numerous different ways which, along with uncontrolled effects of other semantic dimensions, could explain inconsistencies in previous behavioural and neuroimaging findings. Therefore, we used a new set of validated socialness norms to compare social and non-social concept processing, while controlling for other important semantic variables, including concreteness and valence. Moreover, we report the first investigation of a potential link between socialness and motivation. This was aimed at unpacking the informational content or processes that make social concepts unique, and particularly a putative role of reward processing. We report two pre-registered experiments, one behavioural (N = 89) and one using fMRI (N = 30), examining the orthogonal effects of socialness and motivation on semantic decisions. Consistent with other recent findings, social concepts were processed more accurately than non-social concepts. Left TP activation was driven by social concepts but not non-social concepts, and some evidence suggests there is also right TP selectivity for social concepts. All concept types engaged the left vlATL and bilateral aMTG suggestive of a category-general semantic function. Multivariate pattern analyses found distinguishable local activation patterns associated with social vs non-social concepts in the left TP, aMTG, and vlATL, and for high vs low motivation concepts in the left TP and aMTG. The activation patterns encoding socialness in the aMTG generalized across levels of motivation, suggesting that socialness is encoded independently of motivation-related information. These findings align with multiple representation theories that claim social and reward-related experiences contribute to semantic representation. They also particularly pinpoint the left TP as having a preference for social concepts.

## 1. Introduction

Embodied cognition models of semantic representation are highly influential accounts that have emphasised the role of sensory and motor experience in shaping conceptual knowledge. There is growing recognition, however, that other forms of experience, such as language, emotion, cognitive states and social interactions, make a key contribution to the learning and informational content of concepts (e.g., Barsalou, 2020). In the present study, we explored the roles of information derived from social experience and motivated cognition in semantic processing.

In recent years there has been a surge of interest in how the brain represents socially relevant concepts like TRUST and DEMOCRACY (Borghi et al., 2022; Pexman et al., 2023). The neuroimaging literature frequently treats social concepts as a discrete class, with a special, or even privileged status, over other types of semantic knowledge (Conca et al., 2021; Ross & Olson, 2010; Zahn et al., 2007). Such perspectives stem from evidence for selective engagement of lateral anterior temporal lobe (ATL) regions for social relative to non-social concepts (Lin et al., 2015, 2018; Ross & Olson, 2010; Zahn et al., 2007).

However, the strength of these claims is greatly limited by inconsistencies in the way ‘socialness’ has been defined (Pexman et al., 2023). Indeed, existing neural data could have been biased by confounds and other potential explanations which require further exploration.

When it comes to motivation-related information and its influence on semantic processing, there has been little discussion in the contemporary literature (until very recently; see Giurgea et al., 2025). This is surprising given that motivation and the pursuit of reward greatly influence the types and frequency of experiences we engage in and, therefore, will shape the ways in which we learn the meaning of things (Hughes & Zaki, 2015; Madan, 2017). Motivation may also be important for understanding the informational content or processes that make social concepts unique; social motivation is important for driving interpersonal interactions and maintaining relationships (McClelland et al., 1953; Quirin et al., 2013) and could be a mechanism for ‘grounding’ certain social concepts (i.e., linking meaning to bodily experiences, including internal sensations; Reilly et al., 2025). Moreover, reward-related processing has been closely linked to social semantic processing in neurocognitive accounts of acquired social impairment (Belder et al., 2023; Rankin, 2020).

Therefore, the present study had the following primary aims: 1) assess how the socialness of concepts shapes behavioural and neural responses during semantic decisions; 2) examine, for the first time, how motivation-related information might also influence these responses; and 3) evaluate whether social semantic effects can be accounted for, at least in part, by motivation-related information. We expand on these aims below.

### 1.2. Social semantic effects on behaviour and neural activity

In recent work (Diveica et al., 2023, 2024b; Pexman et al., 2023) we have sought to better understand the degree to which, and the contexts in which, social experience makes a unique contribution to concept processing. This involved addressing important limitations of the prior literature. First, there is the fundamental question of what is meant by socialness.

Indeed, socialness can be defined in numerous ways, and social concepts have been distinguished from non-social concepts on divergent sets of criteria (Pexman et al., 2023). On one hand, socialness has been operationalised as the degree to which a stimulus is suggestive of social contexts, as opposed to scenarios involving individuals acting alone. On the other hand, socialness has been an index of the scale of interaction/number of agents or the degree to which a referent has human-like intentions (Pexman et al., 2023; see Table 1). Adding further complexity is the fact that, in many cases, these definitions have been implemented without formal and at-scale attempts to establish their validity and reliability. This raises the following question: to what degree has prior research into social concepts been studying the same cognitive processes and phenomena?

**Table 1.**
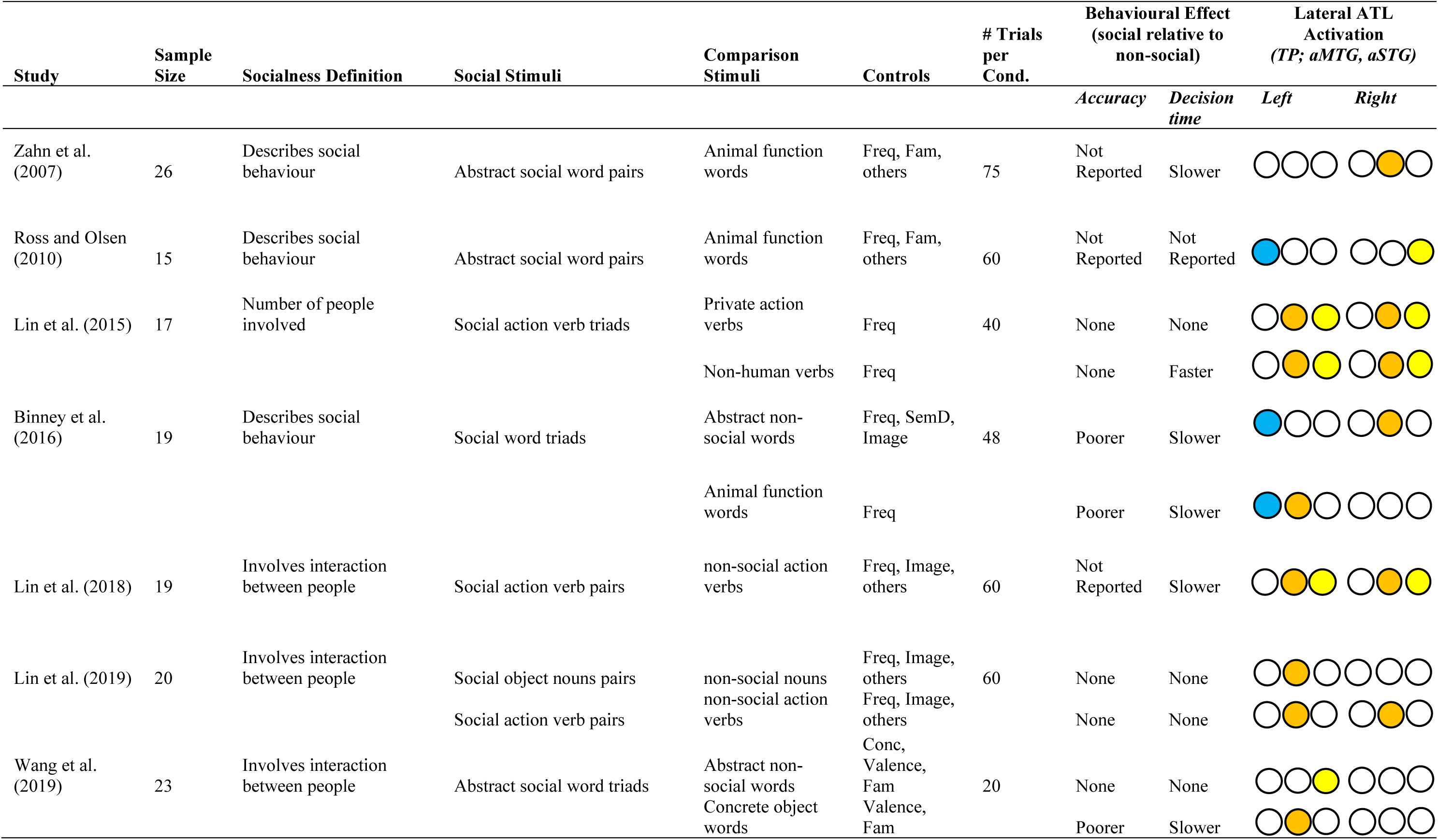
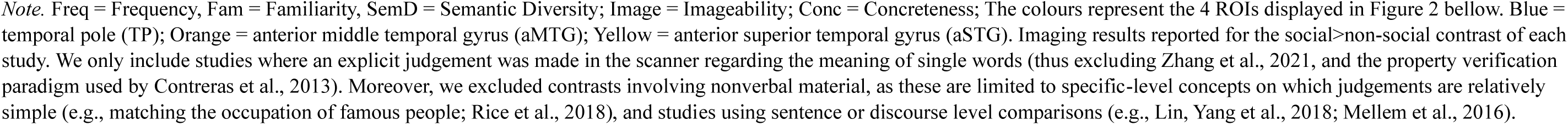
Details of representative prior fMRI studies examining social concept processing.

A second limitation is the fact there has been limited research investigating the behavioural consequences of concepts’ socialness, despite behavioural relevance being a gold standard for psychological theory. Prior to our recent work (Diveica et al., 2023, 2024b), the only behavioural data to be published regarding processing of social versus non-social concepts was that acquired in the context of neuroimaging experiments. These experiments tend to have relatively small sample sizes (see Table 1), and sometimes small numbers of trials, and thus are likely to have very limited power to detect the effects of semantic variables on decision times and task accuracy. It is unsurprising, therefore, that these studies collectively delivered a mixed set of results. In Table 1, we summarize the behavioural findings from representative studies identified in recent meta-analyses and reviews of neuroimaging studies (e.g., Conca et al., 2021; Pexman et al., 2023). Half of these studies reported that social concepts are processed more slowly than non-social concepts, even after controlling for other semantic variables known to influence decision times, like familiarity, imageability and semantic diversity (Binney et al., 2016; Lin et al., 2018; Zahn et al., 2007); the other half found there was no effect on decision times (Lin et al., 2015; Lin et al., 2019; Wang et al., 2019). Only one study reported accuracy scores, with inhibitory effects for social concepts when abstractness was controlled for (Binney et al., 2016). It is difficult to draw a clear conclusion from this set of results, particularly because of the issue of power and false negatives, but also because the definition of socialness, the stimuli, and the contrasts (e.g., the nature of the non-social baseline condition) differ in nontrivial ways.

The neuroimaging data are also inconsistent, further limiting the conclusions that can be drawn about the ‘special status’ of social concepts. Specifically, while the studies to date all converge on the ATL as a broad region of interest, the specific subregion and hemisphere implicated varies considerably (see Figure 1 and Table 1 for a summary). Early studies implicated the right anterior superior temporal gyrus (aSTG; Zahn et al., 2007), the left temporal pole (TP), and the right middle temporal gyrus (aMTG; Ross & Olson, 2010). Activation of the left TP/aMTG and right aSTG were later replicated by Binney et al. (2016) using tighter controls. More recent fMRI work further provides evidence for a role of more posterior left aMTG (Lin et al., 2015; Lin et al., 2019; Wang et al., 2019), the bilateral anterior superior temporal sulcus (STS; Lin et al., 2015; Lin et al., 2018; Lin et al., 2019), and the bilateral aSTG (Lin et al., 2019). Meta-analytic approaches aggregating reported peak activation foci across studies converge on the aMTG and aSTS bilaterally, but more so in the left hemisphere (Arioli et al., 2021; Zhang et al, 2021; also see Hung et al., 2020). It is unclear what is driving this variability and, as such, it is difficult to pinpoint which ATL subregions are critical for social concept processing. At the very least, some of this variance may be accounted for by the use of different socialness definitions and by differences among other stimulus properties like word class, concreteness and emotional valence (see Table 1).

**Figure 1.**
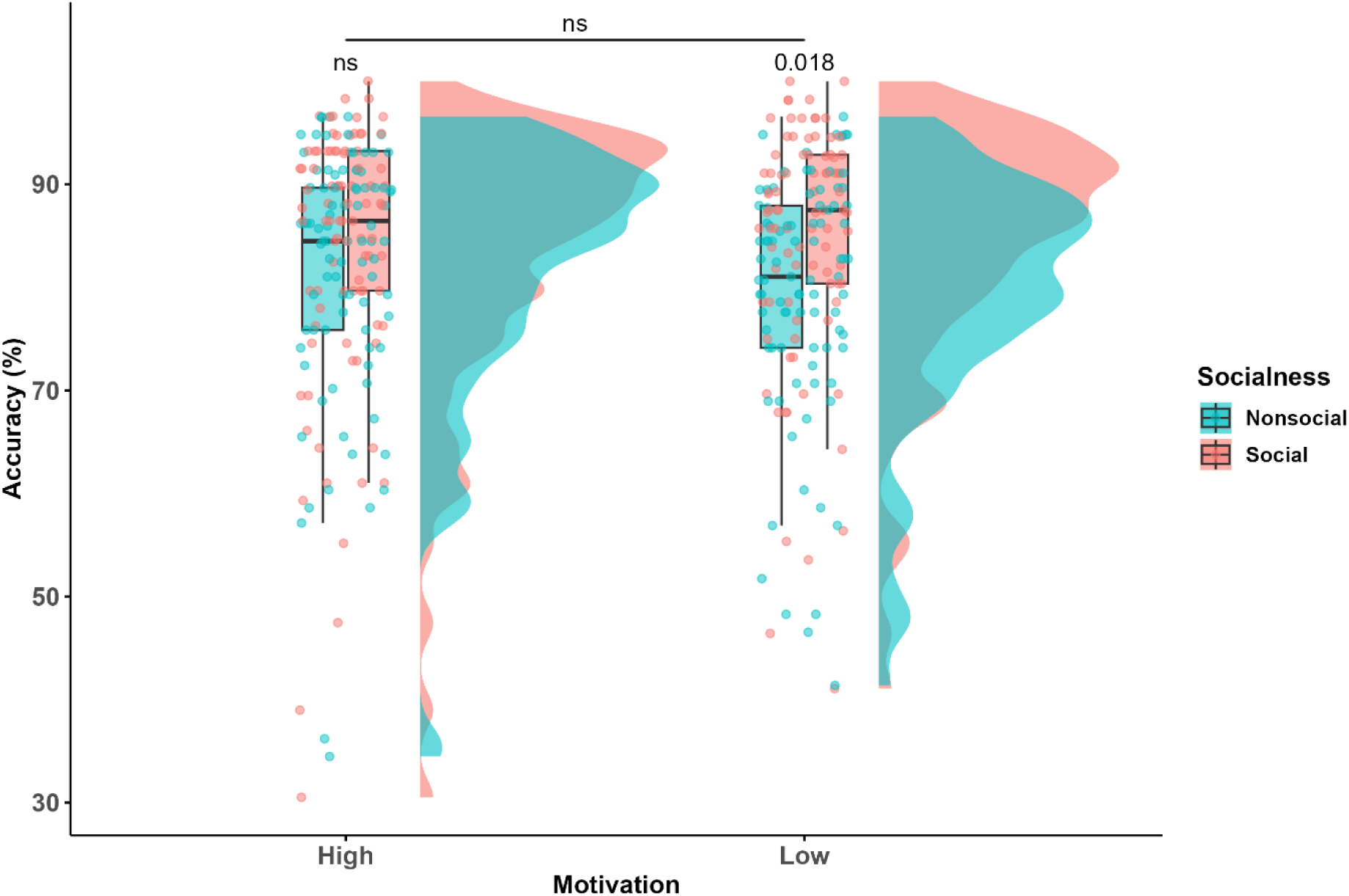
Raincloud plot illustrating the effect of socialness on response accuracy as a function of motivation in the behavioural online study.

The matter of definition and the paucity of behavioural data were directly addressed by Diveica et al. (2023) who collected and validated new socialness norms for over 8000 English words using a Likert scale and a broad definition of socialness that encompassed various kinds of social experience, including roles, actions and attributes. They demonstrated that socialness ratings capture aspects of word meaning that are distinct from other established semantic variables, including concreteness and emotional valence. Further, they capitalised on openly available data from so called behavioural ‘mega studies’ and demonstrated, for the first time, that socialness ratings can explain unique variance in behavioural responses in lexical tasks, in line with the proposal that social experience enriches conceptual knowledge. Diveica et al. (2024b) replicated these findings in further well-powered behavioural investigations, confirming a facilitatory effect of socialness in single-word tasks: auditory lexical decision, recognition memory, concrete semantic judgements, and syntactic classification (also see Goriachun et al., 2025). They also revealed these effects are task-dependent, raising the question of whether they would be reliably observed in the kinds of tasks used in neuroimaging studies (see Section 1.4 for further discussion and related aims of the present study). These newly validated norms now offer the opportunity to address the limitations of prior neuroimaging data, and to reevaluate and refine neurobiological accounts of semantic representation, and particularly the claim that social concepts recruit additional, possibly specialised brain regions. This was the primary aim of the present study, which we shall elaborate in the following section.

### 1.3. Functional specialisation across the ATL: general and social semantics, and the role of reward processing

The anterior temporal lobe (ATL) plays an important role in social cognition (Binney & Ramsey, 2020; Frith & Frith, 2010, 2003; Olson et al., 2013). Damage to this region impairs a wide range of social abilities and behaviours (Binney, Henry, et al., 2016; Irish et al., 2014; Klüver & Bucy, 1938; Rouse et al., 2024) and functional neuroimaging studies of neurotypicals suggest a ubiquitous involvement in high level social processing including moral cognition and mental state attribution (Balgova et al., 2022, 2024; Diveica et al., 2021; Molenberghs et al., 2016; Schurz et al., 2020). The ATL also plays a critical role supporting a long-term store of general conceptual-level knowledge that is involved in transforming sensory inputs into meaningful experiences, and guiding meaning-driven interactions with our environment (Lambon Ralph et al., 2017). This claim is supported by neuropsychological data (Mion et al., 2010; Patterson et al., 2007), as well as brain stimulation and electrophysiological studies, and functional neuroimaging (Binney et al., 2010; Lambon Ralph et al., 2017). To reconcile these parallel observations, it is argued that ATL involvement in social tasks reflects access to conceptual knowledge which provides a fuller understanding of social information and interactions (Binney & Ramsey, 2020; but also Frith & Frith, 2003; Gallagher & Frith, 2003). In support of this claim, there is evidence of parts of the ATL being commonly activated by both social and semantic tasks (Balgova et al., 2022; Balgova et al., 2024).

Nonetheless, there appears to be some functional specialisation in the ATL, in that social semantic knowledge appears to activate lateral ATL regions, including the STG, STS and MTG, more than non-social concepts (e.g., Binney et al., 2016; Zahn et al., 2007; also see the previous section). This contrasts with ventrolateral ATL regions (vlATL; the fusiform and inferior temporal gyri) which are equally engaged by social and non-social concepts, and abstract or concrete concepts of any kind (Binney et al., 2016; Hoffman et al., 2015; Muraki et al., 2025; Rice et al., 2015, 2018; Visser et al., 2012). Functional distinctions such as these are not surprising, as the ATL (broadly defined as the anterior half of the temporal lobe) is comprised of a substantial volume of cortex, amongst which there are numerous subdivisions identifiable based on morphology, cytoarchitecture and/or connectivity (Binney et al., 2012; Ding et al., 2009; Pascual et al., 2015). The *graded semantic hub* hypothesis accounts for these functional distinctions by describing the ATL as a continuous representational space, all of which is engaged by the representation and retrieval of semantic information, but in ways that vary according to proximity and connectivity to sensorimotor systems, as well as affective, language and other cognitive systems (Bajada et al., 2019; Binney et al., 2012, 2016; Jung et al., 2017; Visser & Lambon Ralph, 2011; for a computational exploration of this general hypothesis, see Plaut, 2002).

At the centre of this semantic space, the vlATL represents the point of maximal integration of inputs and its responses are invariant to the types of semantic features that are accessed (Lambon Ralph et al., 2017; Patterson et al., 2007). Towards the edges of this space, however, there may be gradual shifts in semantic function such that regions on this periphery become relatively more specialised. For example, it has been proposed that lateral ATL regions respond more to social concepts because they receive projections from limbic regions (Bajada et al., 2017; Binney et al., 2012; Papinutto et al., 2016) which carry emotion-related information that may be particularly relevant for these types of concepts (Binney et al., 2016). The ATL is implicated in processing valence information and, interestingly, controlling for valence in fMRI studies has impacted on the locus of socialness effects in the ATL (Wang et al., 2019). However, ratings along affective dimensions (e.g., valence, arousal, dominance; Warriner et al., 2013) are only moderately correlate with socialness ratings (Diveica et al., 2023; but see Wang et al., 2019 and Yao et al., 2025 for interactions in behavioural data) and do not appear to fully explain the preferential response of lateral ATL regions to social concepts during semantic relatedness judgements (Wang et al., 2019; cf. Mellem et al., 2016). Therefore, whether socialness is a distinct driver of ATL activation remains an open question which we sought to address in the present study via the following specific questions: (i) does socialness per se drive preferential activation of lateral ATL sub-regions, especially when valence is controlled for?; and (ii) might other types of semantic information account for the socialness effect?

To address the first question, we used the new set of socialness ratings while leveraging the availability of other norms for the same large set of words to tightly control for various lexical and semantic variables (lexical frequency, concreteness, age of acquisition, valence, and semantic diversity). This tight matching has not been possible in prior research because bespoke ratings of socialness were acquired on an ad hoc basis for each study and only for small set of stimuli.

To address the second question, we focused on reward-related information for two reasons. First, it has been proposed that social concepts are more likely than non-social concepts to have hedonic evaluations contribute to their meaning (Arioli et al., 2021; Rankin, 2020). Hedonic evaluations are described as motivation-related processes which are distinct from hedonic affective experience, and Giurgea et al. (2025) recently demonstrated that it is possible to reliably measure a concept’s association with ‘motivation’, and that this is only moderately correlated with affective dimensions of word meaning. Giurgea et al. (2025) went on to demonstrate that ‘motivation’ ratings explain unique variance in participant performance on lexical, semantic, and recognition memory tasks. This suggests that motivation-related information contributes to conceptual processing, but the relationship between motivation and socialness (Arioli et al, 2021; Rankin, 2020), however, has yet to be formally tested.

Second, the lateral and dorsal ATL are strongly connected to orbitofrontal cortex (OFC) and ventromedial frontal regions that are involved in reward processing (via the uncinate fasciculus; Bajada et al., 2017; Binney et al., 2012; Papinutto et al., 2016). Thus, from a *graded hub* perspective, the apparent selectivity of lateral and dorsal ATL sub-regions for processing social concepts could reflect a role in integrating motivation computed by frontal regions into semantic representations (Rankin, 2020). The OFC and subgenual cingulate cortex (sgACC), in particular, are thought to support reward-based decision making and value estimation of actions or events (even those which have not been directly experienced; Klein-Flügge et al., 2022; Rogers et al., 2004; Rudebeck & Murray, 2014). As such, they may be a source of motivation-related information for semantic processing.

Intriguingly, there is some evidence that atrophy to the OFC and sgACC correlates with impairment in comprehension of social concepts (Rijpma et al., 2023). There has yet to be a direct test of the association between lateral ATL activation and motivation-related semantic information using fMRI, and of whether motivation-related information can account for the socialness effect in lexical-semantic processing.

### 1.4. The Present Study

The present study had two parts. First, we performed a behavioural study to examine the effects of socialness while tightly controlling for several lexical and semantic control variables that have well-established effects on lexical-semantic performance, namely number of letters, frequency (Brysbaert & New, 2009), age of acquisition (Kuperman et al., 2012), concreteness (Brysbaert et al., 2014), valence (Warriner et al., 2013), and semantic diversity (Hoffman et al., 2013). A key methodological feature was the use of an alternative forced-choice semantic relatedness judgement task, which has been used in many of the prior neuroimaging studies. This is important because our recent studies that report facilitatory socialness effects on behavioural performance (i.e., faster reaction times) examined tasks involving rapid judgements about lexical status, word class or a general semantic quality (e.g., abstract/concrete decisions) of single words. However, semantic effects, including socialness effects, are task-dependent (Diveica et al., 2024b; Goriachun et al., 2025) and do not always generalize to more complex tasks where relationships between different concepts need to be evaluated. Thus, we sought to close this gap and glean further insight by assessing the socialness effects, and effects associated with motivation ratings (Giurgea et al., 2025), in a semantic relatedness task.

The second part was an fMRI study performed on an independent sample using the same task, stimuli and controls. The purpose was to reevaluate socialness effects at the neural level (see the previous section). It was also the first attempt to investigate whether motivation ratings modulate brain activity during semantic processing. Finally, in light of proposals that social concepts are more likely than non-social concepts to have motivation contribute to their meaning (Arioli et al., 2021; Rankin, 2020) we asked specifically whether socialness is distinct from motivation-related information in both experiments.

If there were to be behavioural effects, we expected these to be facilitatory effects of both socialness and motivation, in line with ‘semantic richness’ (Pexman, 2012) effects observed in single-word tasks (Diveica et al., 2024b; Giurgea et al., 2025). At the neural level, we predicted all concept types would engage the vlATL (the fusiform/inferior temporal gyri), and equally so, given its role as an omni-category supramodal semantic hub (Binney et al., 2016; Muraki et al., 2025). To assess this, we used ATL-optimised imaging sequences capable of acquiring signal from across the whole region (Balgova et al., 2022; Muraki et al., 2025). Moreover, we anticipated social and motivation-related concepts might preferentially activate lateral ATL sub-regions (the STG, STS and MTG) given their connectivity to limbic and ventromedial prefrontal region involved in affective and reward-related processing (see above; Bajada et al., 2017; Binney et al., 2012).

## 2. Experiment 1: Online behavioural study of socialness and motivation effects

First, we report a large (N=90) behavioural study that examined effects associated with socialness and motivation ratings while tightly controlling for several lexical and semantic control variables that have well-established effects on behaviour. Our hypothesis was that socialness and motivation will each facilitate semantic processing, such that there will be a main effect for either dimension pertaining to faster decision times and fewer errors for social words and words with meaning strongly linked to motivation, respectively (Hypotheses 1 & 2). Moreover, if socialness and motivation capture distinct aspects of word meaning, then we hypothesised that there will be an additive facilitatory effect of these variables on semantic processing, such that responses in a two-alternative forced-choice semantic judgment task will be fastest and most accurate in trials involving social words that are also strongly linked to motivation. These will be followed by non-social words that are strongly associated with motivation, and social words that are weakly associated with motivation. Decisions will be slowest and least accurate for non-social words that are weakly associated with motivation (Hypothesis 3). Alternatively, if ratings of socialness and the association with motivation are capturing the same dimension of word meaning, then we hypothesised that there will be an interaction between these variables, such that there will only be an effect of socialness on decision times and accuracy for words that are weakly associated with motivation, but not those which are strongly associated. Similarly, there will be an effect of the association with motivation on decision times and accuracy for words that are weakly associated with socialness, but not those which are strongly associated. Put another way, words that are both social and strongly linked to motivation will be processed as fast and as accurately as words with only one of these characteristics (Hypothesis 4).

### 2.1. Methods

#### 2.1.1. Participants

Participants were recruited online using Prolific (2022; https://www.prolific.com). All participants were first-language English speakers, with no self-reported history of neurological or psychiatric conditions, or language impairments. Participants reported that they had completed, on average, 16 years of formal education (SD = 4.3). All had a 100% approval rate on the Prolific online platform and they were compensated £4 for their participation. Ninety participants took part in this experiment (43 female, M_age_ = 37.83; SD_age_ = 12.19; ranging 20-66 years).

We determined our pre-registered (https://osf.io/3chvw/overview) sample size by conducting a simulation-based power-analysis using the methodology and script from Diveica et al. (2024b). As recommended by DeBruine and Barr (2021), we ran simulations based on effect size estimates from pilot data (n = 10; data collection followed the same procedures described in the present study) to make sure that our study has more than .8 power with a .05 alpha error probability to detect an effect of socialness and motivation on RTs (ms) using linear mixed effect modelling. We ran 1000 iterations and calculated power by computing the percentage of iterations in which we observed a significant effect of socialness on RTs. Our power analysis suggested that 80 participants would provide 0.87 power to detect a main effect of socialness with an effect size of beta = 27.15. However, this sample provides only 2% power to detect an effect of motivation (with a beta of -2.79) and 28% power to detect an interaction (with a beta of 26.13). We also performed a simulation-based sensitivity analysis to estimate the power (α = .05) provided by a sample of 80 participants to detect a range of combinations of socialness, motivation and interaction effect sizes, by calculating the percentage of iterations in which we observed a significant effect. This set of analyses indicated that 80 participants would afford 91% power to detect a 30ms difference between concepts strongly associated with motivation and those weakly associated with motivation. This effect size is realistic as it is similar to those observed in previous investigations of semantic richness effects (Muraki et al., 2022). This sample would also afford 82% power to detect an interaction with a beta of 50ms. To account for any data exclusions, we decided to collect 90 datasets. The pilot data and script used are available at https://osf.io/r5dj6/.

#### 2.1.2. Experimental and Baseline Tasks

Participants completed a two-alternative forced-choice semantic relatedness judgement task (SJT) in which they were required to indicate which of two word choices (a target and a distractor) is more related in meaning to a probe word. On each trial, the probe was presented in the middle of the screen, while the target and the distractor were presented slightly below and either on the left or the right (in a triangular array). The baseline task was a 2-alternative forced-choice numerical judgement task (NJT) in which the participants were asked to indicate which of two 2-digit numbers is closer in magnitude to a probe number (also 2-digit). The NJT was selected as a baseline task to provide comparable visual stimuli and decision type to that of the SJT, based on the assumption that it would require similar levels of attention and general cognitive effort, but minimal semantic processing (Binney et al., 2016; Hoffman et al., 2015).

#### 2.1.3. Stimuli

All stimuli were checked in a pilot study (stimuli and data available on the OSF project page) for any outliers in RT or accuracy that could have indicated general differences in difficulty between stimuli, and also to estimate the required sample size for the behavioural study.

##### SJT

There were 240 SJT trials, each comprising one probe, one target, and one distractor. Each trial belonged to one of four different conditions (60 trials per condition). These were: (a) Non-Social Low Motivation; (b) Non-Social High Motivation; (c) Social Low Motivation; (d) Social High Motivation. Of the 240 probe items, 151 were nouns, 24 were verbs, and 65 were adjectives (see Section S1 of Supplementary Materials (SM) for breakdown by condition). Trials were categorised by the semantic properties of the probe item, and specifically the socialness ratings reported by Diveica et al. (2023; range: 1 (low) to 7 (high)), and motivation ratings reported by Giurgea et al. (2025; range: 1 (low) to 7 (high)). The non-social probe words have mean socialness ratings that range between 1.2 and 3.7, while social probe words have mean socialness ratings ranging between 4.5 and 6.7. The low motivation probe words have mean motivation ratings that range between 1.6 and 3.5, while the high motivation probe words have mean ratings ranging between 4.5 and 6.7. Wilcoxon t-tests confirmed there were significant differences between the social and non-social conditions on mean socialness ratings, and between high motivation and low motivation conditions on mean motivation ratings (all p<0.05; see Table 2 for mean psycholinguistic properties).

**Table 2.** Mean psycholinguistic properties

| Property | Probes |  |  |  | Targets |  |  |  | Distractors |  |  |  |
| --- | --- | --- | --- | --- | --- | --- | --- | --- | --- | --- | --- | --- |
|  | Non-social |  | Social |  | Non-social |  | Social |  | Non-social |  | Social |  |
|  | LowMot | HighMot | LowMot | HighMot | LowMot | HighMot | LowMot | HighMot | LowMot | HighMot | LowMot | HighMot |
| Length | 7.35 | 7.70 | 7.65 | 7.87 | 7.12 | 7.32 | 7.72 | 8.25 | 6.95 | 7.13 | 7.27 | 8.02 |
| Frequency | 2.12 | 2.09 | 2.06 | 2.13 | 2.37 | 2.43 | 2.20 | 2.35 | 2.23 | 2.27 | 2.15 | 2.34 |
| Age of Acquisition | 9.40 | 9.68 | 9.82 | 10.01 | 8.93 | 8.54 | 9.36 | 9.60 | 9.36 | 9.28 | 9.48 | 9.22 |
| Concreteness | 3.10 | 2.94 | 3.08 | 2.79 | 3.08 | 2.90 | 3.02 | 2.76 | 2.92 | 2.79 | 3.08 | 2.81 |
| Valence | 5.16 | 5.53 | 4.88 | 5.16 | 5.22 | 5.40 | 5.13 | 5.31 | 5.45 | 5.65 | 5.55 | 5.61 |
| Socialness | 2.91 | 3.02 | 5.24 | 5.29 | 2.91 | 3.19 | 5.16 | 5.36 | 3.00 | 3.15 | 5.17 | 5.20 |
| Motivation | 2.79 | 4.97 | 2.93 | 5.00 | 3.09 | 4.87 | 3.26 | 4.97 | 2.97 | 4.90 | 3.16 | 5.03 |
| Semantic Diversity | 1.62 | 1.67 | 1.60 | 1.61 | 1.73 | 1.84 | 1.62 | 1.69 | 1.66 | 1.64 | 1.56 | 1.62 |

*Note.* Mot = Motivation

When selecting individual probe stimuli, we also matched the conditions on 6 different psycholinguistic control variables known to influence lexical and semantic task performance: number of letters, log-frequency (Brysbaert & New, 2009), rating-based age of acquisition (AoA, Kuperman et al., 2012), concreteness (Brysbaert et al., 2014), valence (Warriner et al., 2013), and semantic diversity (Hoffman et al., 2013). This was achieved using the R shiny app LexOPS (Taylor et al., 2020) and manual refinement, and pairwise Wilcoxon tests confirmed there were no significant differences between any of the four conditions on these variables (all p>0.05, see SM Section 1). We also controlled as much as possible for part of speech as defined by Brysbaert et al. (2012).

Target words were similar in meaning or associated with the probe. Where possible (e.g., if there was more than one alternative), we selected target words that were similar to the probe in terms of socialness and motivation ratings, and also the number of letters, log-frequency, concreteness, AoA, valence and semantic diversity ratings/scores. Finally, for each trial, distractor items (i.e., the incorrect answer) were selected to be semantically unrelated/dissimilar to the probe, but to have a comparable number of letters, log-frequency, concreteness, AoA, valence, semantic diversity scores, socialness, and motivation scores/ratings to the probe (see Table 3 for example stimuli). Pairwise Wilcoxon t-tests revealed that, at the set level, there were significant differences between the social and non-social conditions on mean socialness ratings in the case of the probes, targets and distractors, as intended. This was also the case for the high versus low motivation conditions in terms of mean motivation ratings. Regarding matching for all the control variables, further pairwise Wilcoxon t-tests confirmed there were no significant differences between conditions for either probes, targets or distractors, with the following exceptions: AoA and semantic diversity for targets (see SM Section 1). One of the distractors was repeated within the same condition, and 7 targets and 15 probes were also used as distractors.

**Table 3.** Example stimuli

| Non-social |  |  |  |  |  | Social |  |  |  |  |  |
| --- | --- | --- | --- | --- | --- | --- | --- | --- | --- | --- | --- |
| Low Motivation |  |  | High Motivation |  |  | Low Motivation |  |  | High Motivation |  |  |
| Probe | Target | Distractor | Probe | Target | Distractor | Probe | Target | Distractor | Probe | Target | Distractor |
| eyesight | glimpse | slumber | doomsday | catastrophe | ache | custodian | keeper | conductor | rebellion | revolution | tournament |
| prop | rotate | juggle | brighten | lighten | recharge | console | solace | confer | flirt | embrace | connect |
| cottage | farmhouse | register | avalanche | snowstorm | dynamite | toddler | youngster | historian | pregnancy | birth | bargaining |
| cancel | abolish | adjourn | heal | fix | aim | cater | dine | lend | execute | punish | employ |
| den | burrow | guinea | caffeine | stimulant | velocity | columnist | reviewer | valentine | investor | collaborator | storyteller |
| whimsical | unusual | unfit | optimum | favorable | spirited | eloquent | verbal | tribal | lovable | comforting | patriotic |
| illusion | mirage | basis | phenomenon | rarity | mission | prank | joke | lunch | expert | specialist | kidnapper |
| window | door | jingle | budget | cost | diet | secretary | messenger | heir | excursion | venture | fitness |
| incomplete | defective | eastern | alert | quick | hungry | careless | reckless | communal | vengeful | murderous | voluntary |
| semester | term | height | jolt | collision | countdown | government | administration | player | flattery | adoration | evolution |

We also took steps to ensure that the main effects of socialness or motivation were not confounded with the level of difficulty in either selecting the target or rejecting the distractor. This involved comparing the four conditions in terms of mean semantic similarity scores between (i) probes and targets, and (ii) probes and distractors, respectively. Semantic similarity was estimated using a custom measure; we extended the sensorimotor distance measure of Wingfield and Connell (2023), which used 11 dimensions of sensorimotor experience (Lynott et al., 2019), to include ratings on valence, arousal, and socialness, and thereby also capture social and affective experience. We created a vector of the 14 ratings for each word and then computed the reverse cosine values (1 minus the cosine of the angle between the vectors) for each pair of words. There were no significant differences between any of the pairs of conditions in terms of semantic similarity between probes and targets, or probes and distractors (see pre-registration). Moreover, as would be expected, semantic similarity was, on average, smaller (i.e., smaller mean reverse cosine distance) between probes and distractors than between probes and targets in all conditions (see Table 3 for example stimuli).

### NJT

The NJT comprised 128 number trials, each containing one probe, one target and one distractor. The probes were randomly generated 2-digit numbers between 0 and 99, while the targets and distractors were chosen based on how close or far, respectively, they were in magnitude to the probe. On average, trials had large numerical distances between the probe and the target (greatest difference = 24) and probe and distractor (greatest difference = 26), but small differences between the target and distractor (greatest difference = 2), as Banks et al. (1976) and pilot testing have shown this increases the difficulty of the judgements, with smaller gaps between numbers associated with longer RT, and we wanted to ensure a similar overall level of difficulty between the SJT and NJT.

#### 2.1.4. Procedure

The study was conducted online via the Gorilla (Anwyl-Irvine et al., 2020; www.gorilla.sc) testing platform. Each SJT trial began with a black fixation cross presented for 500ms, which was replaced by a word triad. The position of the target (i.e., word pair for SJT or closest number for NJT) was varied so that it was in the left hand position for 50% of trials and in the right hand position for 50% of trials. The word stimuli remained on the screen for 2250ms or until a response was made, after which the next trial began immediately. The participants were instructed to press the “r” or the “g” keys on their keyboard to indicate that the word on the left or the right, respectively, was more similar in meaning to the probe. No indication of which hand should be used to respond was provided. They were also asked to respond as quickly and accurately as possible. The NJT had the same format except it began with a blue fixation cross, and participants were required to pick which of 2 numbers was closest in value to the probe. Before commencing the main experiment, the participants completed a practice session where they were presented with 12 NJT trials and 16 SJT trials. In a deviation from the pre-registration, the participants did not receive any feedback on performance during the practice. The order of presentation of the conditions followed a fixed sequence (which was repeated 4 times). However, the stimuli belonging to the respective conditions were randomised using Gorilla’s "randomise trials" function.

#### 2.1.5. Data Cleaning

We sequentially implemented exclusion criteria at the participant, item, and individual trial level. Data from participants with an overall accuracy score less than 0.571 (equivalent to above chance accuracy as determined by a binomial test using a p-value of 0.05) were excluded. Data associated with words that received correct responses from less than half of the remaining participants were excluded. Finally, trials with responses faster than 250ms or greater than +3 SD from each participant’s mean RT were excluded. Data from participants who did not complete the experiment in full were deleted and the participants were asked to return their submission in Prolific.

We collected a total of 33121 observations. We excluded data from one participant who had below-chance accuracy. Nine trials were excluded because less than 50% of the participants provided correct responses. Of the experimental trials, 1142 (5.29%) were identified as RT outliers and were excluded from the analyses. The final analyses included 20458 observations, out of which 16825 were responded to correctly.

#### 2.1.6. Analyses

The analysis plan was pre-registered at https://osf.io/r5dj6/. All data cleaning, analyses, and visualizations were performed in RStudio (version 4.3.3, 2025). We used the packages “lme4” (Bates et al., 2018) and “afex” (Singmann et al., 2017) to perform our analyses. A linear mixed effects model was used to test the possible effect of socialness and motivation information on RTs. The analysis was restricted to the correct word trials. Word socialness and motivation were effect-coded (social/high motivation: -0.5; non-social/low motivation: +0.5). We started by fitting the maximal model including the random effects structure justified by the design using the following formula: RT ∼ Socialness*Motivation + (1 + Socialness*Motivation| Participant) + (1|Word). Given that we did not have hypotheses about the random effects, we tested whether the maximal model was overfitted by following the procedure outlined by Bates et al. (2018) to identify the optimally parsimonious model for our data. We used likelihood ratio tests for model comparison. To test the possible effect of socialness and motivation on response accuracy, a logistic mixed-effects model was used on all word trials (correct and incorrect), following the same procedure as for the model predicting RTs. The R package “lmerTest” (Kuznetsova et al., 2017) was used to compute p-values for the estimated mixed-effects models via the Satterthwaite’s degrees of freedom method (Giesbrecht & Burns, 1985; Hrong-Tai Fai & Cornelius, 1996). An α level of *p* < .05 was used to make inferences about the statistical significance of the results.

We also conducted a version of the frequentist mixed effects models above using Bayesian mixed-effects regression, which allowed us to quantify evidence in favour of null effects (Diveica et al., 2024b; Vasishth et al., 2018). We conducted a region of practical equivalence analysis (ROPE), which calculates what percentage of the 95% most credible values (i.e., the highest density interval; HDI) for each estimated parameter falls within a range of values that are equivalent to 0 (i.e., representing the null). We followed the recommendation of a range of ± 0.1 SDs in the dependent variable as the ROPE (Kruschke, 2018), which has been termed the equivalent of a negligible effect size (Cohen, 1988; Diveica et al., 2024b). Similarly, we did not aim to make a discrete decision regarding the null or alternate hypothesis, and we report the percentage of the HDI that falls within the ROPE (Kruschke, 2018), which refers to the percentage of the most likely effect sizes that can be considered negligible. We also report the probability of direction (PD), which varies between 50% and 100%, and quantifies the certainty of the direction of an effect (i.e., whether it is positive or negative). Similar to the p-value threshold in frequentist statistics, this is sensitive only to the amount of evidence for the alternative hypothesis (Makowski et al., 2019). These analyses were conducted using the R packages “rstanarm” (Goodrich et al., 2020) and “bayestestR” (Makowski et al., 2019).

### 2.2. Results

The descriptive statistics for both tasks are presented in Table 4. Initial pairwise Wilcoxon tests revealed no significant differences between SJT conditions and the NJT in RT or accuracy. There were also no significant differences between any of the SJT conditions.

**Table 4.**
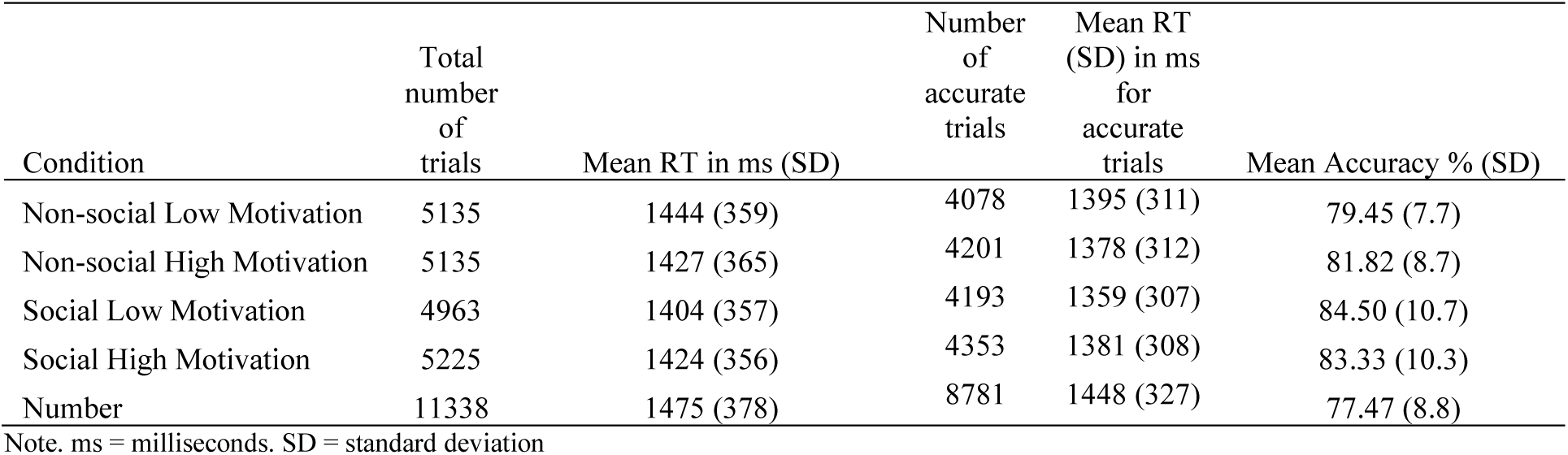
Descriptive statistics by condition after data cleaning

| Condition | Total number of trials | Mean RT in ms (SD) | Number of accurate trials | Mean RT (SD) in ms for accurate trials | Mean Accuracy % (SD) |
| --- | --- | --- | --- | --- | --- |
| Non-social Low Motivation | 5135 | 1444 (359) | 4078 | 1395 (311) | 79.45 (7.7) |
| Non-social High Motivation | 5135 | 1427 (365) | 4201 | 1378 (312) | 81.82 (8.7) |
| Social Low Motivation | 4963 | 1404 (357) | 4193 | 1359 (307) | 84.50 (10.7) |
| Social High Motivation | 5225 | 1424 (356) | 4353 | 1381 (308) | 83.33 (10.3) |
| Number | 11338 | 1475 (378) | 8781 | 1448 (327) | 77.47 (8.8) |

Note. ms = milliseconds. SD = standard deviation

The raw RT data did not meet the assumption of normality and, thus, we conducted all the below analyses on log transformed RTs. The model with the optimal random effects structure for our data included one random intercept per participant and one random intercept per word. This was implemented using the following formula: logRT ∼ Socialness*Motivation + (1|Participant) + (1|Word). The main effects of socialness and motivation, and their interaction were nonsignificant (see Table 5 for summary of the mixed effects model). The Bayesian analyses estimated that 85% of the HDI for socialness (93.59% PD), 100% for motivation (51.53% PD), and 37.38% for the interaction (93.49% PD) fell within the ROPE (-0.01 – 0.01).

**Table 5.** Mixed-effects models predicting RT and accuracy in the online behavioural experiment

| *Predictors* | **Log RT** |  |  | **Accuracy** |  |  |
| --- | --- | --- | --- | --- | --- | --- |
|  | *Estimates* | *CI* | *p* | *Odds Ratios* | *CI* | *p* |
| (Intercept) | 3.13 | 3.12 – 3.14 | **<0.001** | 6.53 | 5.37 – 7.95 | **<0.001** |
| Socialness | 0.01 | -0.00 – 0.01 | 0.132 | 0.77 | 0.62 – 0.96 | **0.018** |
| Motivation | 0.00 | -0.01 – 0.01 | 0.973 | 0.98 | 0.79 – 1.21 | 0.833 |
| Socialness X Motivation | 0.01 | -0.00 – 0.03 | 0.121 | 0.75 | 0.49 – 1.15 | 0.188 |
| **Random Effects** |  |  |  |  |  |  |
| $\sigma^2$ | 0.01 | | | 3.29 | | |
| $\tau_{00}$ Word | 0.00 | | | 0.58 | | |
| $\tau_{00}$ PID | 0.00 | | | 0.62 | | |
| ICC | 0.37 |  |  | 0.27 |  |  |
| N<sub>PID</sub> | 89 |  |  | 89 |  |  |
| N<sub>Word</sub> | 231 |  |  | 231 |  |  |
| Observations | 16825 |  |  | 20458 |  |  |
| Marginal R<sup>2</sup> / Conditional R<sup>2</sup> | 0.002 / 0.375 |  |  | 0.005 / 0.270 |  |  |

Note. CI = confidence interval; Socialness is a factor variable with social words set as the reference group. Motivation is a factor variable with high motivation set as the reference group. Accuracy is a binary dependent variable with inaccurate responses as the reference group (0) and accurate responses as the focus group (1). The marginal R<sup>2</sup> includes only the variance from the fixed effects and the conditional R<sup>2</sup> includes variance from both the fixed and random effects.

The model with the optimal random effects structure for the accuracy data also included one random intercept per participant and one random intercept per word: Accuracy ∼ Socialness*Motivation + (1| Participant) + (1|Word). The main effect of socialness was significant, showing that responses were more accurate for words high in socialness ratings (see Figure 1 and Table 5 for summary of the mixed effect model). We also performed post-hoc tests to explore simple effects using emmeans (2.0.0; Lenth, 2025). These analyses showed that social concepts had higher accuracy scores than non-social concepts, specifically for words associated with low motivation. In contrast, the effect of socialness was not significant for words associated with high motivation (see Section S2 in SM). The effect of motivation and the interaction between socialness and motivation were nonsignificant. The Bayesian analysis estimated that 23.21% of the HDI for socialness (98.96% PD), 93.75% for motivation (58.02% PD), and 30.35% for the interaction (90.57% PD) fell within the ROPE (-0.181 – 0.181).

### 2.3. Summary and Interim Discussion

Socialness had an effect on response accuracy; trials containing social words were judged more accurately than trials containing non-social words. The simple main effects further revealed the effect of socialness was specific to trials containing words weakly related to motivation. There were no other effects, including on RTs. Thus, we only partially confirmed Hypothesis 1 that there would be facilitatory effects of socialness on semantic processing.

This finding nonetheless aligns with the semantic richness effects observed in single-word tasks (Diveica et al., 2023, 2024b), and provides evidence that socialness effects generalize, to some degree, to the SJT.

We did not observe any effects of motivation on behavioural performance, with the Bayesian analyses providing support for the null hypothesis. Therefore, our data do not provide support for Hypothesis 2 that motivation-related information influences semantic processing. This contrasts with previous research showing that motivation-related information relates to behavioural performance in single-word lexical and semantic tasks (Giurgea et al., 2025). Notably, our study was not sufficiently powered to detect motivation effects smaller than a ∼30 ms difference between high and low motivation trials.

Consequently, better-powered studies will be needed to more conclusively assess the possible existence of subtler effects.

The presence of the socialness effect only when motivation-related information was absent points towards a potential interaction between the two dimensions. This pattern aligns more with Hypothesis 4 than Hypothesis 3, and an overlap between the dimensions rather than independence, with motivation-related information potentially constraining the influence of other social information on semantic processing. The interaction was not statistically significant in the frequentist analyses, but the Bayesian analyses provided only limited support for the null hypothesis. Therefore, these findings do not offer strong evidence for either the presence or absence of an interaction, and they may be limited by statistical power to detect an interaction. A higher-powered study is therefore needed to more definitely assess potential interactive effects between socialness and motivation on semantic task performance.

The failure to observe more robust socialness effects (extending to RT) and any motivation effects on behaviour may reflect the specific demands of the task employed. The SJT requires associative judgements between word pairs, as well as comparison of association strength across pairs, leading to longer overall RTs than tasks like lexical decision or syntactic classification, where responses to single words typically occur within about one second (e.g., Diveica et al., 2024b). The SJT requires more controlled, reflective processing, which may attenuate bottom-up influences like the semantic richness effects observable in single-word tasks.

Relatedly, it has been shown that semantic richness effects, including socialness effects, are task-dependent (Diveica et al., 2024b; Goriachun et al., 2025, 2026). Task demands are thought to constrain the influence of semantic dimensions on behaviour, such that only those aspects of meaning relevant to the decision at hand affect behavioural performance (Muraki et al., 2023; Newcombe et al., 2012; Tousignant & Pexman, 2012). Because target words and distractors were matched on socialness/motivation ratings, socialness/motivation did not provide useful information for determining which word was more related to the probe in a given trial. For this reason, participants may have relied on other features besides socialness or motivation to make their decision, resulting in no widespread processing benefits on RT and accuracy like those observed in other task types.

Another possible explanation for the null results is that by tightly matching across conditions for other psycholinguistic variables, we weakened the socialness and motivation manipulations. For example, the mean socialness rating in the non-social and social conditions was around 3 and 5, respectively, of a 7-point scale, and therefore did not contain the least/most social words. Diveica et al. (2024b) showed socialness effects having used the full socialness continuum amongst their stimuli and slightly higher mean socialness values for the social conditions (e.g., 5.5). Future research could maximise these manipulations and reevaluate, although there might be the disadvantage of weaker matching on other potentially confounding variables, which was a strength of the current study.

Finally, the fact there were no differences in accuracy or RT between the SJT and NJT confirmed that the number task would be a suitable baseline condition in the fMRI experiment for controlling effects of working memory, attention and executive skills associated with general cognitive processing. Moreover, the fact that there were no large differences in accuracy/RT between SJT conditions means that effects of socialness or motivation at the level of the brain could not be attributed to general cognitive effort.

## 3. Experiment 2: fMRI study of social concepts and motivation-related concepts

We next performed an fMRI study on an independent sample (N=33) using the same task, stimuli and controls. We sought to replicate socialness effects observed at the neural level in several prior studies (Binney et al., 2016; Lin et al., 2015; Lin et al., 2018; Ross & Olson, 2010; Zahn et al., 2007), but using new socialness norms validated at scale, and having carefully controlled for many more lexical and semantic variables than these prior studies. We also sought to investigate, for the first time, whether motivation-related information influences the neural activation underpinning conceptual processing, and explore a potential role of motivation-related information in accounting for activation of lateral ATL regions by social concepts.

The experimental design included two other key features. First, our imaging protocol was optimized to be sensitive to Blood-Oxygen Level-Dependent (BOLD) signal across all parts of the ATL, given that the *graded hub* account predicts that in addition to selectivity to social concepts in lateral ATL regions, there will be general omni-category response in the vlATL. Special measures are needed because the vlATL is especially prone to magnetic susceptibility-induced signal loss and image distortion in conventional echo planar imaging (Devlin et al., 2000; Visser et al., 2010) and, when the field of view in the ATL is limited by such artefacts, it impacts the inferences that are made about the function of this region (e.g., missing the general semantic responses makes it appear as if the region is specialised for social information processing; Binney al., 2016; Balgova et al., 2022). We used the same acquisition parameters and distortion correction procedures as Balgova et al. (2022), which obtained a good signal to noise ratio (SNR) in the ATL (Muraki et al., 2025; see SM Section S4 & Tables S35 & S36 for tSNR of each ROI). We used a dual-echo gradient-echo echo-planar imaging (EPI) fMRI sequence, which acquires images at both a short echo-time (12ms) that is less prone to signal loss due to spin dephasing, and a long echo-time (35ms) that is more sensitive to BOLD contrast. This dual-echo sequence is more effective at detecting signal in inferior temporal regions compared to standard single gradient-echo sequences and spin-echo sequences (Halai et al., 2014, 2015). Moreover, we acquired the images with a left-to-right phase encoding direction, which reduces signal pileup in the inferior ATL (Balgova et al., 2022; Embleton et al., 2010). Finally, we applied a post-acquisition k-space spatial correction procedure to address geometric distortions in the data, which provides a more effective method of distortion correction compared to other commonly used procedures, including those based on B0 field maps (Embleton et al., 2010).

Second, this was the first fMRI study of social concepts to use both (magnitude-based) univariate analyses and (information-based) multivariate analyses. Univariate fMRI analyses assess each voxel independently, identifying regions that are more or less active under specific conditions. Multivariate approaches, by contrast, examine patterns of activation across voxels, detecting distributed representations even without overall amplitude changes. In essence, univariate analyses tell us where changes in activation occur, while multivariate analyses reveal whether information (like socialness) is encoded in local patterns of neural activation (Haynes & Rees, 2006; Norman et al., 2006).

Our preregistered predictions for this experiment were as follows. First, we expected vlATL to respond equivalently (in magnitude) for all types of concepts, consistent with a role as a supramodal and omni-category semantic hub (Binney et al., 2016; H1). Moreover, if the Graded Semantic Hub Hypothesis (Binney et al., 2016) is correct, then any activation differences between specific semantic task conditions and the baseline number task in lateral ATL regions will be smaller in magnitude than those in the vlATL (H2; Rice et al., 2018).

Second, if socialness plays a unique role in the representation of conceptual knowledge in the ATLs, univariate analyses will reveal a main effect of socialness in the lateral ATL such that it is more strongly activated by social as compared to non-social words. This will hold true both for words that are strongly and weakly associated with motivation (H3). We did not explicitly pre-register hypotheses for the MVPA analysis but, in line with univariate predictions, we expected them to yield above-chance classification for social versus non-social trials in the lateral ATL. Third, we predicted that, if word meaning relies on systems involved in reward-related processing, then there will be analogous effects to those predicted for socialness either in lateral ATL regions or the ventral striatum, subgenual anterior cingulate cortex or orbitofrontal cortex reflecting the known association of these regions with reward processing (Martins et al., 2021; Rijpma et al., 2023; H4). Finally, if socialness and motivation ratings make independent contributions to concept representation in the ATLs, this will be reflected by: (i) an additive effect in the same ATL area, such that it will be activated the most by social words that are strongly linked to motivation, followed by non-social words that are strongly linked to motivation, as well as social words that are weakly associated with motivation. It will be activated the least by non-social words that are weakly associated with motivation; and/or (ii) spatially non-overlapping main effects (i.e., they occur in distinguishable areas of temporal cortex; H5). MVPA analyses should reveal that patterns of local activation enable above-chance classification of socialness that generalizes across levels of association with motivation, and vice versa.

### 3.1. Methods

#### 3.1.1. Participants

Thirty-three participants took part in this experiment (female = 22). All participants were aged between 18-40 years (M = 22.88; SD = 4.65), first-language English speakers, with normal or corrected-to-normal vision, no history of neurological or psychiatric conditions or language impairments and were assessed to be right-handed using the Edinburgh Handedness Inventory (Oldfield, 1971). On average, participants had completed 15.82 years (SD = 4.72) of formal education. We removed data from three participants who had below chance performance on the NJT. The final sample consisted of 30 participants (19 female, M_age_ = 22.9; SD_age_ = 4.65, age range = 19-40; education = 15.73 years (SD = 4.88)). The study was conducted at Bangor University and was approved by the local research ethics review committee. Participants received monetary compensation in exchange for study participation (£5 for every 30 minutes of participation).

#### 3.1.2. Experimental and Baseline Tasks

The two tasks and stimuli were the same as those used in Experiment 1.

#### 3.1.3. Experimental Procedure

A PC running PsychoPy (Version 2023.2.3; Peirce et al., 2019) was used for the presentation of stimuli and recording of responses. Participants completed 4 runs, each lasting approximately 7 minutes. Within a run, the SJT and NJT alternated in a block design with a 14 second rest after each number block. There were four SJT blocks each containing 15 trials (block duration range = 66750ms - 73550ms). There were four NJT blocks each containing 8 trials (block duration = 22000ms). Each trial began with a 500ms fixation cross (blue cross on number trials, black cross on word trials) presented in the middle of the screen and then a number or word was presented for 2250ms in black, lower-case font on a white background. After the 2250ms response window, word trials had a variable interstimulus interval (0- 13100ms; mean ISI = 2120ms). Number trials were presented without an inter-trial interval, appearing in immediate succession. Participants were instructed to respond as quickly and accurately as possible after the word or number appeared on the screen using two buttons indicating either “left” or “right” on an MR-compatible response box, held in their right-hand.

Nested within the block design, we employed a rapid event-related design for the presentation of the four different semantic conditions. This was achieved using a pseudorandomized sequence of conditions (1 trial per sequence position) that spanned the four SJT blocks and was used in all four runs. We created 4 lists, each containing 15 trials from all four of the SJT conditions (60 word triads in total). Those lists were matched across the same set of psycholinguistic and lexical variables that the complete sets of stimuli were matched for. Each list was used once across the four runs. The list/run order was counterbalanced across participants using a balanced Latin Square design. The condition order within each run followed a fixed sequence (e.g., begin with a block of 6 number trials followed by an SJT trial from the social high motivation condition, then a trial from the non-social high motivation condition, and so on), but the exact stimuli were randomly selected from the appropriate list. Similarly, four unique lists of 32 numbers were created and the order in which they were presented was counterbalanced across participants. All study materials are available for download on the OSF project page (https://osf.io/r5dj6/).

In the same scanning session, the participants also completed further runs involving the same tasks but different stimuli as part of a separate study that is not reported here. The order of completion of the two studies was counterbalanced to avoid fatigue effects.

#### 3.1.4. Imaging Acquisition

All imaging was performed on a 3.0T Philips Elition MRI scanner with a 32-channel head coil and a SENSE factor of 2.5. We used a dual-echo gradient-echo EPI fMRI sequence to acquire 31 axial slices covering the whole brain in ascending sequential order with the following parameters: short echo time (TE) = 12ms and long TE = 35ms, repetition time (TR) = 2000ms, and flip angle = 85°. The functional EPIs were acquired with an in-plane reconstructed resolution of 2.5 x 2.5 mm and a slice thickness of 4mm voxels (reconstruction matrix = 96 × 96; FOV (mm) = 240 × 240 × 124). We acquired five dummy scans prior to each run (to develop a steady state magnetisation), followed by 215 volumes per run. We acquired three main functional runs with a single direction k-space traversal in the left-right phase-encoding direction. In addition, we acquired a short EPI ‘pre-scan’ with the participants at rest. The parameters of the pre-scan matched the main functional scans with the exception that they included interleaved dual direction k-space traversals. This provided us with 10 pairs of images with opposing direction distortions (10 left–right and 10 right–left) which were used in a k-space distortion correction procedure for addressing geometric distortion and mislocalisation due to magnetic susceptibility artifacts. We acquired a high-resolution T2-weighted scan to assess the accuracy of our distortion correction procedure, which contained 36 slices covering the whole brain with the following parameters: TE = 78ms, TR = 3410ms, flip angle = 90°. The T2-weighted scan was acquired with a reconstructed voxel size (mm) of 0.6 x 0.6 x 4 and a slice thickness of 4 mm voxels with a 1 mm slice gap (reconstruction matrix = 400 x 370; FOV (mm) = 240 x 240 x 179). In addition, we used a T1-weighted imaging sequence to acquire an anatomical image used to estimate the transformation of our functional images into Montreal Neurological Institute (MNI) space. The T1 images contained 175 slices covering the whole brain and were acquired with the following parameters: TE = 18ms, TR = 3.5ms, flip angle = 8°, reconstructed voxel size (mm) = 0.94 x 0.94 x 1 and reconstruction matrix = 240 x 240. All images were acquired with a 30° axial tilt.

#### 3.1.5. Data Analysis

##### Within-scanner behavioural data

All analyses were performed in RStudio (version 4.3.3; 2025). We collected a total of 7920 observations for the SJT and 4224 for the NJT. Following exclusions, the datasets were reduced to 7200 and 3840 observations, respectively. Task response time was assessed using a linear mixed effects model, with a random effect of participant and a fixed effect of trial. To assess task accuracy, we used a logistic mixed effects model, with a random effect of participant and a fixed effect of trial. We only included correct trials in our response time models. Inferential statistics were calculated to compare performance in the SJT and NJT, and compare between conditions in the SJT, which allowed us to detect any differences in task/condition difficulty and check overall engagement with the semantic task and non-semantic active baseline task.

##### fMRI distortion correction and preprocessing

fMRI preprocessing and analyses were conducted using the statistical software package MATLAB 2023b (The MathWorks Inc., 2023) and Statistical Parametric Mapping software (SPM; version 12; Welcome Trust Centre for Neuroimaging, London, UK). A spatial remapping correction was computed separately for images acquired at the long and the short echo time using the method reported in Embleton et al. (2010), implemented via an in-house MATLAB script (available on the OSF project page https://osf.io/r5dj6/), as well as SPM12’s 6-parameter rigid body registration algorithm. In the first step, each functional volume was registered to the mean of the 10 pre-scan volumes acquired at the same echo time. This step is both a required part of the distortion correction procedure and corrects the timeseries for differences in participant positioning in between functional runs and for minor motion artefacts within a run. Next, one spatial transformation matrix per echo time was calculated from the opposingly distorted pre-scan images. These transformations consisted of the remapping needed to correct geometric distortion and were then applied to each of the main functional volumes. This resulted in two motion and distortion-corrected time-series per run (215 volumes per echo), which were subsequently combined at each timepoint using a simple linear average of image pairs.

Slice-timing correction was performed using the middle slice for reference. The T1- weighted anatomical image was coregistered to a mean of the distortion- and motion-corrected images using a six-parameter rigid-body transform and the normalized mutual information objective function. SPM12’s unified segmentation and normalization function was used to estimate a spatial transform to register the anatomical image to MNI standard stereotaxic space. This transform was subsequently applied to the co-registered functional volumes which were then resampled to a 3 x 3 x 3 mm voxel size. As a final step, the normalized functional images were smoothed using an 8mm full-width half-maximum Gaussian filter.

##### Univariate analyses

Functional data were analysed using the general linear model approach (GLM). We conducted within-subjects fixed effects analyses with all functional runs incorporated into a single GLM per participant. For each semantic condition, SJT trials and NJT trials were modelled as events with a duration of 2.25 seconds. These regressors were convolved with the canonical hemodynamic response function and treated with a high-pass filter (cutoff of 128s). We also entered the extracted motion parameters into the GLM as regressors of no interest. The analyses were restricted to grey matter using an explicit mask generated from a group-level probabilistic tissue map computed using the SPM12 unified segmentation and normalization procedure, and binarized with a 0.5 threshold.

To address the five hypotheses, we conducted both whole brain analyses and region of interest (ROI) analyses using ROIs selected a priori from studies exploring semantic processing and, specifically, social concept processing. Whole-brain multi-subject random effects analyses were conducted on each of the following contrasts: a) to assess the omni-category role of the vlATL: all word trials > numbers, social high motivation > numbers, social low motivation > numbers, non-social high motivation > numbers, non-social low motivation > numbers (H1); b) to assess the simple effects of socialness: social high motivation > non-social high motivation, social low motivation > non-social low motivation (H3); c) to assess the simple effects of motivation: social high motivation > social low motivation, non-social high motivation > non-social low motivation (H5). We performed one-sample t-tests on all these sets of contrast images, restricting the statistical maps to cerebral tissue using the same explicit group-level mask as used in the single subject analyses. We also performed a factorial ANOVA including the contrasts of each semantic conditions minus the number baseline condition to estimate the main effects of socialness (H3) and motivation (H4), and their interaction (H5). Cluster-wise statistical significance was assessed using a cluster-defining voxel-height threshold of *p* < .001 uncorrected, and a family-wise error (FWE) corrected cluster extent threshold at *p* < .05. We conducted additional robustness analyses to ensure the observed effects are not driven by differences in task difficulty between conditions. This involved repeating the GLM analyses with additional modelling of decision time data, and on accurate trials only (discarding inaccurate trials).

We also used a priori ROIs to extract and quantify the magnitude of activation within different ATL subregions, and to increase sensitivity to the hypothesized effects. Coordinates were used to define the centre of mass of spherical ROIs with radii of 8mm. This was implemented using the SPM MarsBar toolbox (Brett et al., 2002). The ROIs were chosen from previous studies that found activation for semantic vs non-semantic stimuli (Muraki et al., 2025), as well as those that found activation for social concepts relative to non-social concepts when controlling for various psycholinguistic variables (e.g., Binney et al., 2016; Wang et al., 2019; see Figure 2).

**Figure 2.**
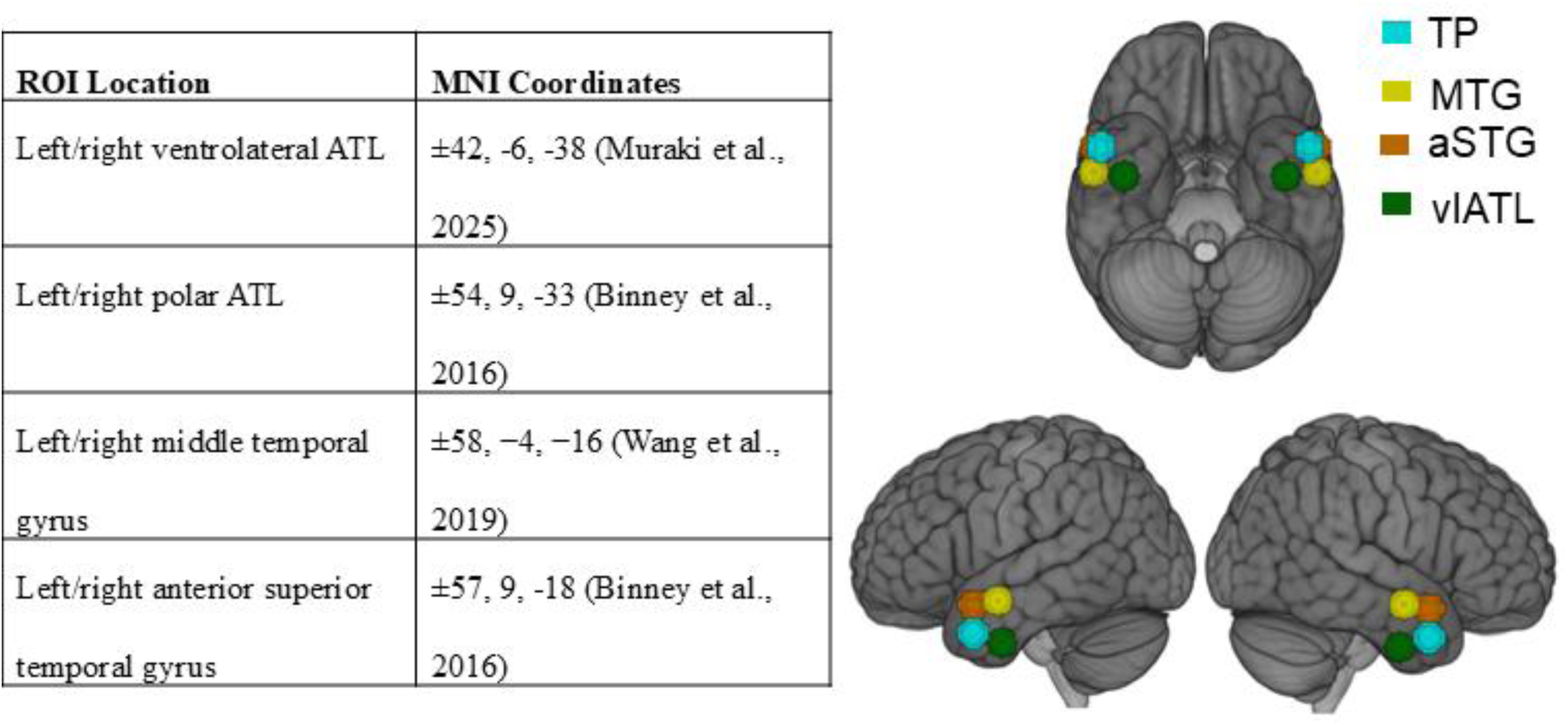
Locations of Spherical Regions of Interest. TP = temporal pole; MTG = middle temporal gyrus; aSTG = anterior superior temporal gyrus; vlATL = ventrolateral ATL; Blue = TP; Yellow = MTG; Orange = aSTG; Green = vlATL

##### Multivariate analyses

We performed a multivoxel pattern classifier analysis (MVPA) to test whether local voxel- activation patterns in the ATL can classify (a) social versus non-social trials, and (b) high motivation versus low motivation trials, suggestive of encoding of social/motivation semantic content (H3 & H4). In addition, we conducted cross-decoding analyses by training the model to classify social versus non-social trials with strong motivation association and then testing its ability to classify social versus non-social trials with weak motivation association (and vice versa). This cross-decoding analysis evaluated whether neural patterns associated with socialness are consistent across levels of association with motivation (and vice versa), indicative of independent encoding of socialness and motivation features (H5). Since reporting of training/testing direction is inconsistent in the literature, we report results for both directions separately in the SM (section S5) and collapsing across directions, with the latter being less susceptible to the influence of potential SNR differences between training/testing trial sets (Van Den Hurk & Op De Beeck, 2019).

We first conducted these analyses within the same pre-defined ROIs used in the univariate analyses. We also conducted a searchlight analysis restricted to the ATL. The ATL search volume was defined by a mask previously described by Hung et al. (2020), and as previously adapted by our group in Muraki et al. (2025). The searchlight analysis involved repeatedly conducting decoding within spherical regions with an 8mm-radius centred on each voxel within the ATL mask. This approach allowed us to assess whether patterns of activation outside the pre-defined ROIs encode the words’ social or motivation semantic content. The size of the sphere was a slight deviation from the pre-registration to ensure consistency with the size of the spheres used in the ROI analyses.

We used a 4-fold take-one-run-out cross-validation and a linear discriminant analysis (LDA) classifier as implemented in the nilearn (Abraham et al., 2014) and scikit-learn (Pedregosa et al., 2011) packages in Python (version 3.13.2; virtual environment and Jupyter Notebook scripts available on the OSF page). For each participant, activation patterns were extracted for each trial via a GLM model (same HRF model, same nuisance regressors, same auto-correlation model and high-pass filtering as the univariate GLM) applied to unsmoothed data. The GLM included one regressor for the trial of interest and a separate regressor for all remaining trials (Mumford et al., 2012). To account for differences in RTs between trials, we included an additional RT duration regressor of no interest that was convolved with the HRF (ConsDurRTDur in Mumford et al., 2024). We performed classification on the resulting z-scored beta-estimates.

We assessed classifier accuracy by calculating the percentage of trials correctly classified. Statistical significance was assessed at the group level with one-sample t-tests against chance accuracy (50%). Given we used a one-tailed t-test with an alpha level of .05, we report 90% confidence intervals (Kennedy et al., 2026; Lakens et al., 2018). For the searchlight analyses, in line with previous studies (Kuhnke et al., 2023; Murphy et al., 2017), we smoothed the first-level classification accuracy maps (minus chance accuracy) with an 8mm FWHM Gaussian kernel using AFNI’s 3dBlurinMask command (version 25.2.18). The group-level accuracy maps were corrected for multiple comparisons using probabilistic threshold-free cluster enhancement as implemented in the R package pTFCE (Spisák et al., 2019) and a threshold of p<.05.

To verify that the main decoding effects were not influenced by differences in accuracy, and, hence, in engagement with semantic processing, we performed a robustness analysis restricted to accurate trials. Excluding inaccurate trials led to unequal trial counts across conditions. As classification accuracy can be influenced by such imbalance, we balanced the trial numbers by determining the minimum number of accurate trials across conditions for each participant and randomly subsampling each participant’s data accordingly.

### 3.2. Results

#### 3.2.1. Behavioural Data

There were no significant differences between the SJT conditions, or between the SJT and the NJT trials, on RT or accuracy (see Section S3 of SM for all results). This largely replicates the results from the large-scale online behavioural study described in Experiment 1, with the exception that in Experiment 1 there was an effect of socialness on accuracy.

#### 3.2.2. Univariate fMRI Analyses Whole Brain Contrasts

##### Semantic versus Number Similarity Judgements

We conducted whole brain univariate analyses to contrast the overall SJT with the active baseline NJT, and each different SJT condition with the NJT. When contrasted against the NJT, the overall SJT was associated with robust activation in the semantic network (see Figure 3A). This included a cluster in the left ATL which comprised the left vlATL (see Section S4 of SM for peak coordinates), and lateral ATL regions including the anterior MTG, STG and STS, extending into the TP. This cluster also extended right along the STS and MTG. In the right hemisphere, ATL activation was focused around the STG/STS and did not extend as ventrally as in the left hemisphere. There was a cluster in the left inferior frontal gyrus (IFG), including pars triangularis and pars orbitalis which have been implicated in semantic control processes (Lambon Ralph et al., 2017). There was also activation in the left superior frontal cortex, the bilateral occipital lobe which could be attributed to greater visual complexity of the word stimuli compared to the number stimuli, and in the right cerebellum (see Table 6).

**Figure 3.**
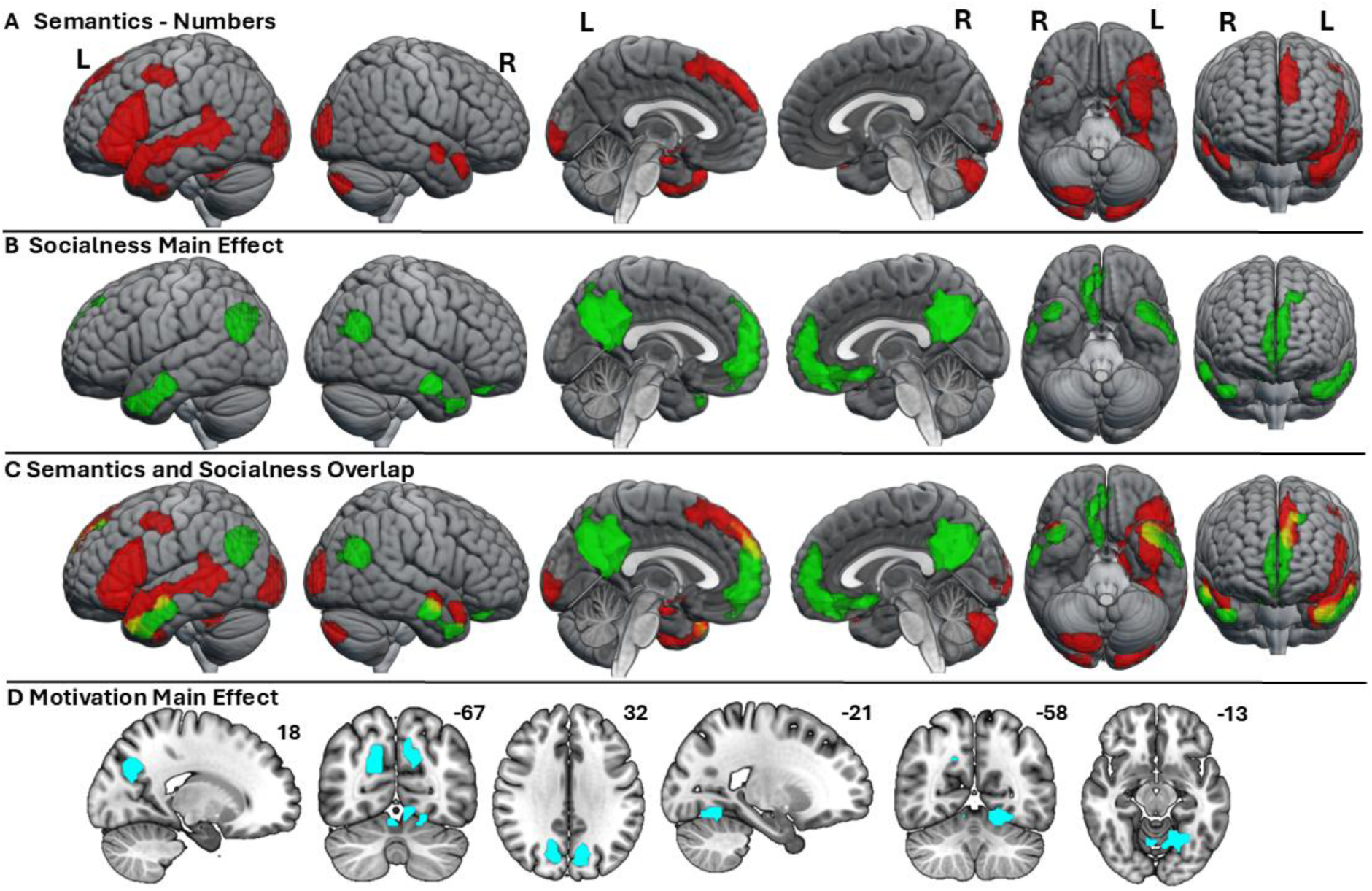
Cortical regions activated by: **A)** all semantic judgement conditions relative to the number judgement condition (one-sample t-test); **B)** social conditions relative to the non-social conditions (main effect from the ANOVA); **C)** overlay of the SJT minus NJT contrast and the socialness main effect; **D)** high motivation conditions relative to low motivation conditions. The statistical maps are thresholded with a voxel-height threshold of p <.001 uncorrected, and an FWE-corrected cluster extent threshold of p<0.05

**Table 6.** Significant activation clusters in the SJT (Words) minus NJT (Numbers) whole- brain analysis

| Cluster name and location of maximum | Cluster extent (voxels) | Peak (Z) | Maxima MNI coordinates (mm) |  |  |
| --- | --- | --- | --- | --- | --- |
|  |  |  | x | y | z |
| **R Cerebellum** | **455** |  |  |  |  |
| Crus II |  | 7.55 | 12 | -79 | -37 |
| **L Fronto-Temporal Cortex** | **3151** |  |  |  |  |
| Inferior frontal gyrus (pars triangularis) |  | 7.15 | -51 | 20 | 26 |
| Middle temporal gyrus |  | 7.1 | -51 | -34 | 2 |
| Inferior frontal gyrus (pars triangularis) |  | 6.91 | -42 | 26 | -1 |
| Hippocampus |  | 6.89 | -24 | -13 | -16 |
| Fusiform gyrus |  | 6.84 | -42 | -43 | -22 |
| Inferior frontal gyrus (pars opercularis) |  | 6.44 | -54 | 17 | 14 |
| Middle temporal gyrus |  | 6.42 | -60 | -43 | 2 |
| Inferior frontal gyrus (pars triangularis) |  | 6.41 | -51 | 26 | 14 |
| **L Occipito-Temporal Cortex** | **390** |  |  |  |  |
| Middle occipital gyrus |  | 6.34 | -15 | -91 | -7 |
| Inferior occipital gyrus |  | 6.14 | -21 | -97 | -10 |
| Middle occipital gyrus |  | 6.08 | -24 | -97 | 2 |
| Inferior occipital gyrus |  | 4.33 | -27 | -82 | -10 |
| **R Occipital Lobe** | **224** |  |  |  |  |
| Calcarine |  | 6.3 | 18 | -94 | -1 |
| Calcarine |  | 5.97 | 24 | -100 | 2 |
| Middle occipital gyrus |  | 5.05 | 27 | -97 | 14 |
| Middle occipital gyrus |  | 4.77 | 36 | -94 | 2 |
| Middle occipital gyrus |  | 3.61 | 36 | -91 | 11 |
| Superior occipital gyrus |  | 3.37 | 30 | -91 | 23 |
| **R Temporal Lobe** | **90** |  |  |  |  |
| Superior temporal pole |  | 5.73 | 48 | 14 | -19 |
| **L Parietal Lobe** | **171** |  |  |  |  |
| Rolandic Operculum |  | 5.59 | -42 | -28 | 23 |
| Rolandic Operculum |  | 4.33 | -48 | -13 | 17 |
| Putamen |  | 3.93 | -30 | -16 | 8 |
| Insula |  | 3.41 | -36 | -1 | 17 |
| **L Frontal Lobe** | **312** |  |  |  |  |
| Superior medial frontal gyrus |  | 5.27 | -9 | 50 | 41 |
| Superior frontal gyrus |  | 5.03 | -12 | 59 | 32 |
| Superior medial frontal gyrus |  | 4.83 | -9 | 35 | 53 |
| Superior frontal gyrus |  | 4.75 | -12 | 41 | 47 |
| Supplementary motor area |  | 4.5 | -3 | 14 | 53 |
| **R Temporal Lobe** | **91** |  |  |  |  |
| Middle temporal gyrus |  | 4.9 | 57 | -4 | -13 |
| **R Parietal Lobe** | **105** |  |  |  |  |
| Heschl's gyrus |  | 4.57 | 39 | -22 | 17 |
| Rolandic Operculum |  | 4.11 | 39 | -7 | 17 |

*Note.* Significant activation clusters in the semantic relatedness judgement task minus number judgement task contrast
identified at $p < .05$ , FWE-corrected, resulting in an extent threshold of $k = 90$ voxels, after a cluster-defining threshold of $p < .001$ uncorrected. Table shows up to 8 local maxima more than 8mm apart in each cluster

When contrasting each semantic condition with the NJT using one-sample t-tests, we found similar patterns of activation of the semantic network (see Figure S2). All conditions were associated with activity in the left vlATL as well as the anterior STG/STS/MTG. All conditions activated semantic control regions in the left IFG and the posterior lateral temporal cortex (PLTC). Notable differences were as follows. In the right ATL, activation was less extensive than in the left ATL and it was limited to lateral regions and only the social conditions. Moreover, the social low motivation condition revealed only right anterior MTG activation whereas in the social high motivation condition this cluster extended to the aSTG and TP (See Figure S2A and B).

##### Socialness and Motivation Effects

Contrasts of each word condition with the NJT were also entered into a full factorial ANOVA. We found a main effect of socialness where social concepts activated the bilateral aMTG extending into the TP (although this was overall stronger in the left hemisphere), the bilateral angular gyrus, medial frontal cortex and the precuneus (see Figure 3B; see Figure 3C for overlay of the main effect of socialness with the effect of task (all semantic conditions minus numbers)). The main effect of motivation was associated with activation in the posterior parahippocampal gyrus/cerebellum and the precuneus/occipital parietal sulcus. However, there was no significant main effect of motivation in the ATL and no significant socialness and motivation interaction effect (see Figure 3D and Table 7).

**Table 7.**
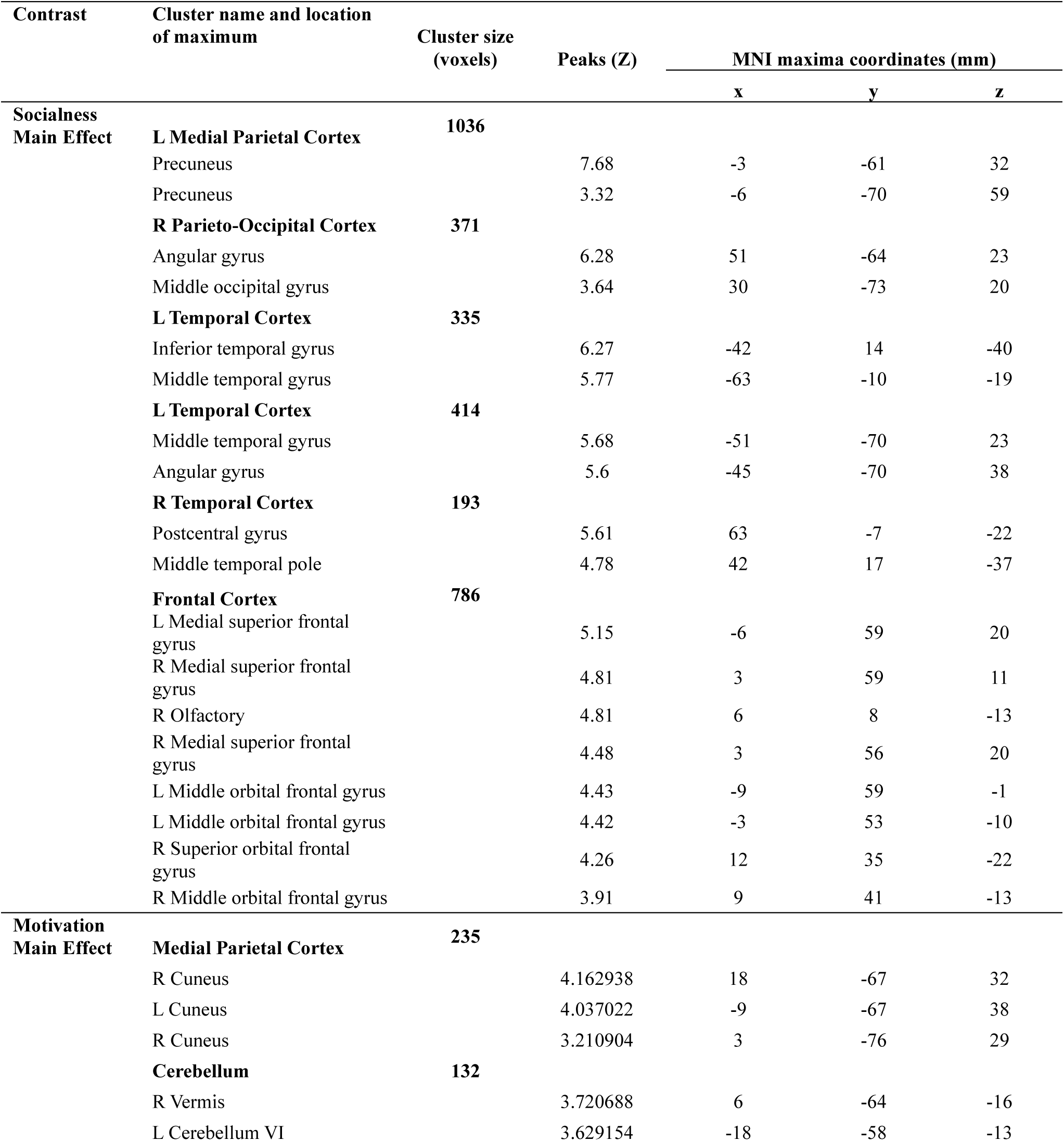
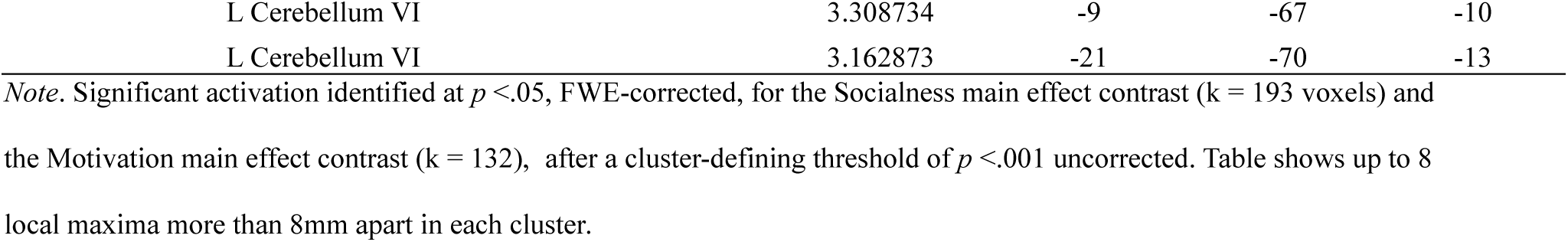
Significant activation cluster in whole-brain ANOVA

| Contrast | Cluster name and location of maximum | Cluster size (voxels) | Peaks (Z) | MNI maxima coordinates (mm) |  |  |
| --- | --- | --- | --- | --- | --- | --- |
|  |  |  |  | x | y | z |
| **Socialness Main Effect** | **L Medial Parietal Cortex** | **1036** |  |  |  |  |
|  | Precuneus |  | 7.68 | -3 | -61 | 32 |
|  | Precuneus |  | 3.32 | -6 | -70 | 59 |
|  | **R Parieto-Occipital Cortex** | **371** |  |  |  |  |
|  | Angular gyrus |  | 6.28 | 51 | -64 | 23 |
|  | Middle occipital gyrus |  | 3.64 | 30 | -73 | 20 |
|  | **L Temporal Cortex** | **335** |  |  |  |  |
|  | Inferior temporal gyrus |  | 6.27 | -42 | 14 | -40 |
|  | Middle temporal gyrus |  | 5.77 | -63 | -10 | -19 |
|  | **L Temporal Cortex** | **414** |  |  |  |  |
|  | Middle temporal gyrus |  | 5.68 | -51 | -70 | 23 |
|  | Angular gyrus |  | 5.6 | -45 | -70 | 38 |
|  | **R Temporal Cortex** | **193** |  |  |  |  |
|  | Postcentral gyrus |  | 5.61 | 63 | -7 | -22 |
|  | Middle temporal pole |  | 4.78 | 42 | 17 | -37 |
|  | **Frontal Cortex** | **786** |  |  |  |  |
|  | L Medial superior frontal gyrus |  | 5.15 | -6 | 59 | 20 |
|  | R Medial superior frontal gyrus |  | 4.81 | 3 | 59 | 11 |
|  | R Olfactory |  | 4.81 | 6 | 8 | -13 |
|  | R Medial superior frontal gyrus |  | 4.48 | 3 | 56 | 20 |
|  | L Middle orbital frontal gyrus |  | 4.43 | -9 | 59 | -1 |
|  | L Middle orbital frontal gyrus |  | 4.42 | -3 | 53 | -10 |
|  | R Superior orbital frontal gyrus |  | 4.26 | 12 | 35 | -22 |
|  | R Middle orbital frontal gyrus |  | 3.91 | 9 | 41 | -13 |
| **Motivation Main Effect** | **Medial Parietal Cortex** | **235** |  |  |  |  |
|  | R Cuneus |  | 4.162938 | 18 | -67 | 32 |
|  | L Cuneus |  | 4.037022 | -9 | -67 | 38 |
|  | R Cuneus |  | 3.210904 | 3 | -76 | 29 |
|  | **Cerebellum** | **132** |  |  |  |  |
|  | R Vermis |  | 3.720688 | 6 | -64 | -16 |
|  | L Cerebellum VI |  | 3.629154 | -18 | -58 | -13 |

|  |  |  |  |  |
| --- | --- | --- | --- | --- |
| L Cerebellum VI | 3.308734 | -9 | -67 | -10 |
| L Cerebellum VI | 3.162873 | -21 | -70 | -13 |

*Note.* Significant activation identified at $p < .05$ , FWE-corrected, for the Socialness main effect contrast ( $k = 193$ voxels) and the Motivation main effect contrast ( $k = 132$ ), after a cluster-defining threshold of $p < .001$ uncorrected. Table shows up to 8 local maxima more than 8mm apart in each cluster.

As pre-registered, we also used one-sample t-tests to examine the simple effects of socialness and motivation. Consistent with the main effect of socialness described above, we found significant simple effects of socialness within both levels of motivation. Within high motivation trials, social trials elicited greater activation in the left MTG (anterior and posterior), superior and middle TP (extending into the inferolateral ATL, including inferior temporal gyrus (ITG)). They also activated left angular gyrus, and bilateral precuneus and medial frontal cortex. Within low motivation trials, social trials elicited greater activation in the bilateral angular gyrus, precuneus, and medial prefrontal cortex, but not the ATL. Finally, there was a significant simple effect of motivation within the social condition, such that high compared to low motivation trials elicited greater activation in the left PLTC, posterior STG, supramarginal gyrus, posterior IFG, and ventral pre- and post-central gyri. There was no significant simple effect of motivation within the non-social condition (see Section S4 of SM for peak coordinates and figures).

##### Robustness Analyses

We repeated the whole-brain analyses to control for potential confounds: 1) processing speed, by including RT as a parametric regressor in the GLM, and 2) misunderstanding of word meaning or lapses in attention, by restricting the analyses to accurate trials. After correcting for multiple comparisons, there was no significant socialness main effect in the right hemisphere and no significant main effect of motivation.

##### ATL ROI Analyses

We extracted regional response magnitudes relative to the NJT from eight a-priori ROIs and compared them across semantic conditions. We focused on the vlATL and 3 lateral ATL ROIs centered on peak coordinates previously shown to exhibit preferential responses to social relative to non-social concepts (see Figure 2). Mean and standard error of activation magnitude for each ROI and each condition are provided in Table S19, while the distributions across participants are displayed in Figure 4.

**Figure 4.**
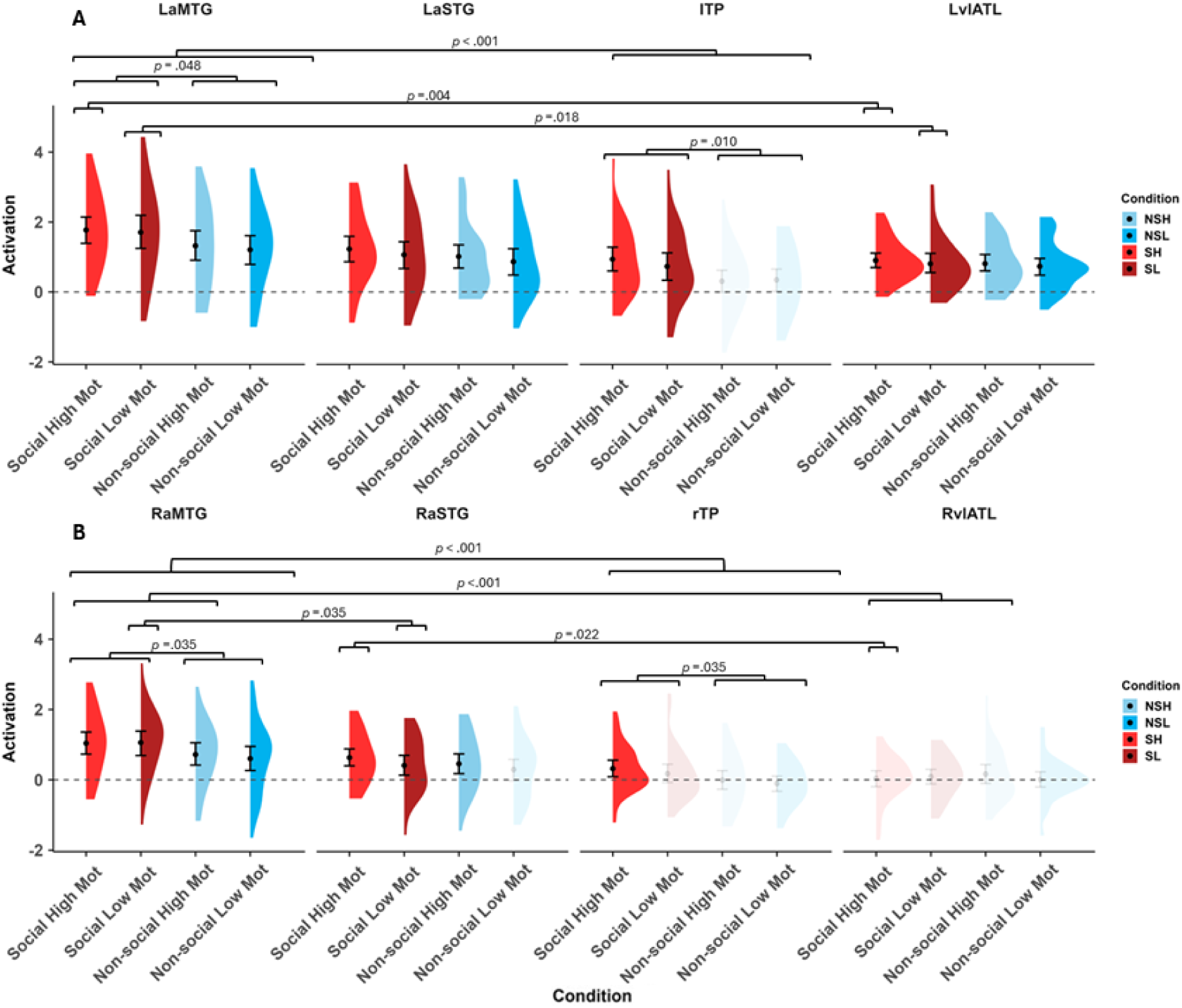
Summary of ROI analysis comparing mean semantic activation (relative to number judgement) in the left (**A**) and right hemisphere (**B**) by condition. The dotted line represents baseline. Error bars represent 95% confidence intervals bootstrapped around the mean. Transparency indicates non-significance in one-sample t-test compared to baseline. Dark red = Social High Motivation (SH). Light Red = Social Low Motivation (SL). Dark Blue = Non-social High Motivation (NSH). Light Blue = Non-social Low Motivation (NSL).

First, we used two-tailed one-sample t-tests to assess whether the four semantic conditions significantly activated the ROIs relative to the NJT baseline, thus testing H1. We observed significant activation for all semantic conditions in the left vlATL (green ROI), the left aSTG (orange ROI), and bilateral aMTG (yellow ROI). The right TP activated significantly above baseline only for social high motivation trials. The left TP (blue) was activated only by the social conditions, while the right aSTG was activated by all semantic conditions except for non-social low motivation (see Table S20 & S27). There was no significant activation for any of the semantic conditions in the right vlATL.

We then conducted separate three-way ANOVAs for each hemisphere, with ROI, socialness, and motivation as factors. We found a significant bilateral main effect of ROI and of socialness (Table S21 & S28). We further inspected the results using Wilcoxon pairwise t-tests. To test H2, we compared activation magnitudes between ROIs separately for each semantic condition. In the left hemisphere, we found significant differences between aMTG and vlATL in all social conditions, and between aMTG and TP in all conditions, such that aMTG displayed greater activation (Table S22). In the right hemisphere, there were significant differences in all conditions between aMTG and TP, and between aMTG and vlATL (except non-social low motivation), as well as between aMTG and aSTG in the social low motivation condition, such that aMTG was associated with greater activation. We also found significant differences between aSTG and vlATL in the social high motivation condition specifically, such that aSTG was associated with greater activation (Table S29). To test H3, we compared activation magnitudes between socialness levels and between motivation levels separately for each ROI. In both hemispheres, activation was greater for social relative to non-social trials in the TP and aMTG (Table S23 & S25; see Figure 4 & Table S30 & S32). There were no significant differences in activation magnitude between motivation levels in any of the ROIs.

Finally, we conducted exploratory (not pre-registered) pairwise Wilcoxon tests to examine differences in activation between hemispheres for each ROI and each semantic condition. We found that there was significantly greater activation in the left vlATL compared to the right vlATL for all semantic conditions. No other comparisons were significant (see Table S34).

#### 3.2.3. Multivariate Analyses ROI-level

##### Pattern Classification

We conducted a pattern classification analysis to assess if the local activation patterns within the pre-defined ROIs encode socialness. This analysis revealed above chance accuracy for classifying socialness in the left vlATL, left TP, and bilateral aMTG across all trials (see Figure 5A; see Section S5 SM for condition-specific analyses). The robustness analysis that included only accurate trials showed the same pattern. These results indicate that socialness is encoded in the local activity patterns of most ATL ROIs.

**Figure 5.**
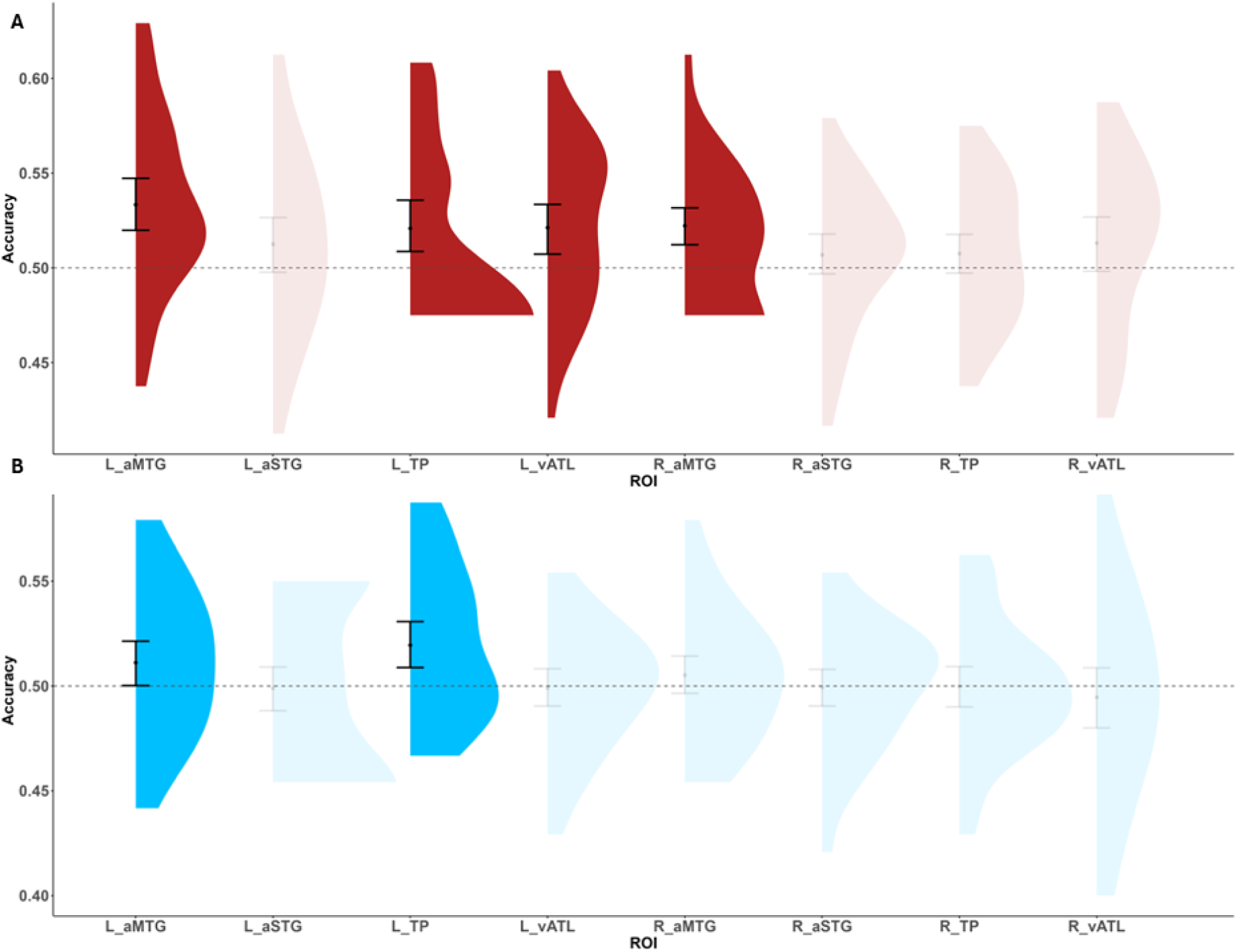
The accuracy with which patterns of local activation within ATL ROIs were able to classify **A**) Socialness within all trials, and **B**) Motivation within all trials. Dotted line represents chance-level (50%) accuracy. Error bars represent 90% confidence intervals bootstrapped around the mean. Transparency indicates non-significance in one-sample t-test.

We conducted a pattern classification analysis to assess if the local activation patterns within the pre-defined ROIs encode motivation. This analysis revealed above chance accuracy for classifying motivation in the left TP and left aMTG across all trials (see Figure 5B; see Section S5 SM for condition-specific analyses). In the robustness analysis in which only accurate trials were considered, only the local activity patterns in the left TP were able to classify motivation significantly above chance. These results indicate that motivation-related information is encoded within specific ATL sub-regions.

##### Cross-decoding

We conducted a cross-decoding analysis to test whether the local activity patterns encoding socialness generalize across levels of motivation, and vice versa. We focus on reporting the results for the ROIs that showed robust pattern classification across all trials (see the ***Pattern Classification*** section above). When decoding socialness across levels of association with motivation, we found significantly above chance classification accuracy only in the left aMTG and bilateral vlATL (see Figure 6A). The robustness analyses which considered only accurate trials confirmed these results, and additionally revealed significantly above chance classification accuracy in the left TP. Motivation was not classified above chance across socialness levels in either of the two ROIs showing above chance classification across all trials (see Section S5 of SM for specific training direction results and additional motivation decoding results).

**Figure 6.**
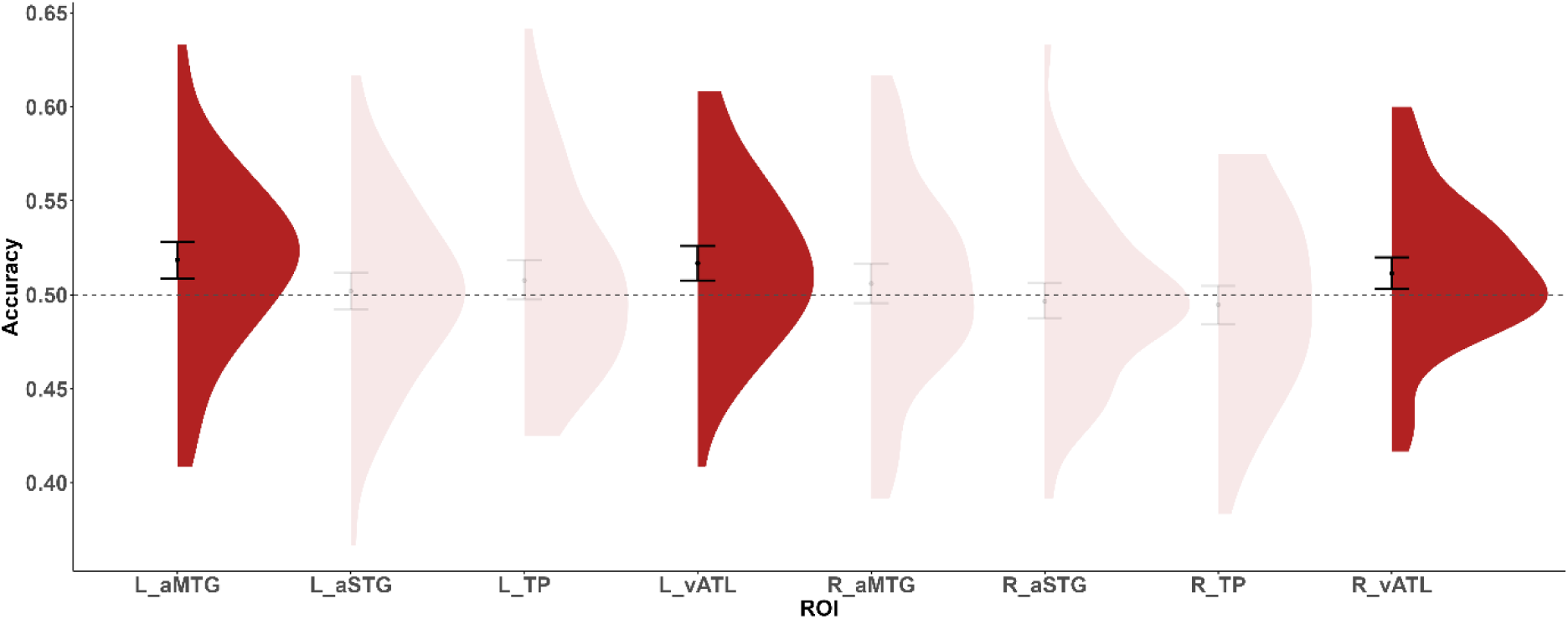
Accuracy levels for cross-decoding Socialness when collapsing across training directions. Dotted line represents chance-level (50%) accuracy. Error bars represent 90% confidence intervals bootstrapped around the mean. Transparency indicates non-significance in one-sample t-test.

##### ATL-restricted Searchlight

###### Pattern Classification

We conducted a searchlight analysis restricted to the ATL to examine whether any local patterns of activation other than in the a priori ROIs might encode socialness and/or motivation. Socialness was classified with above chance accuracy in groups of voxels in the inferolateral (ITG, fusiform gyrus) portion of the left ATL and the right lateral ATL (testing/training on all trials; see Figure 7A; see Section S5 of SM for condition specific results). In the robustness analysis, these groups of voxels were restricted to the left ITG. Motivation was decoded with above chance accuracy in a smaller group of voxels in the left ITG (testing/training on all trials; see Figure 7B; see Section S5 of SM for within-condition results). In the robustness analysis, no voxels showed above chance accuracy when classifying motivation.

**Figure 7.**
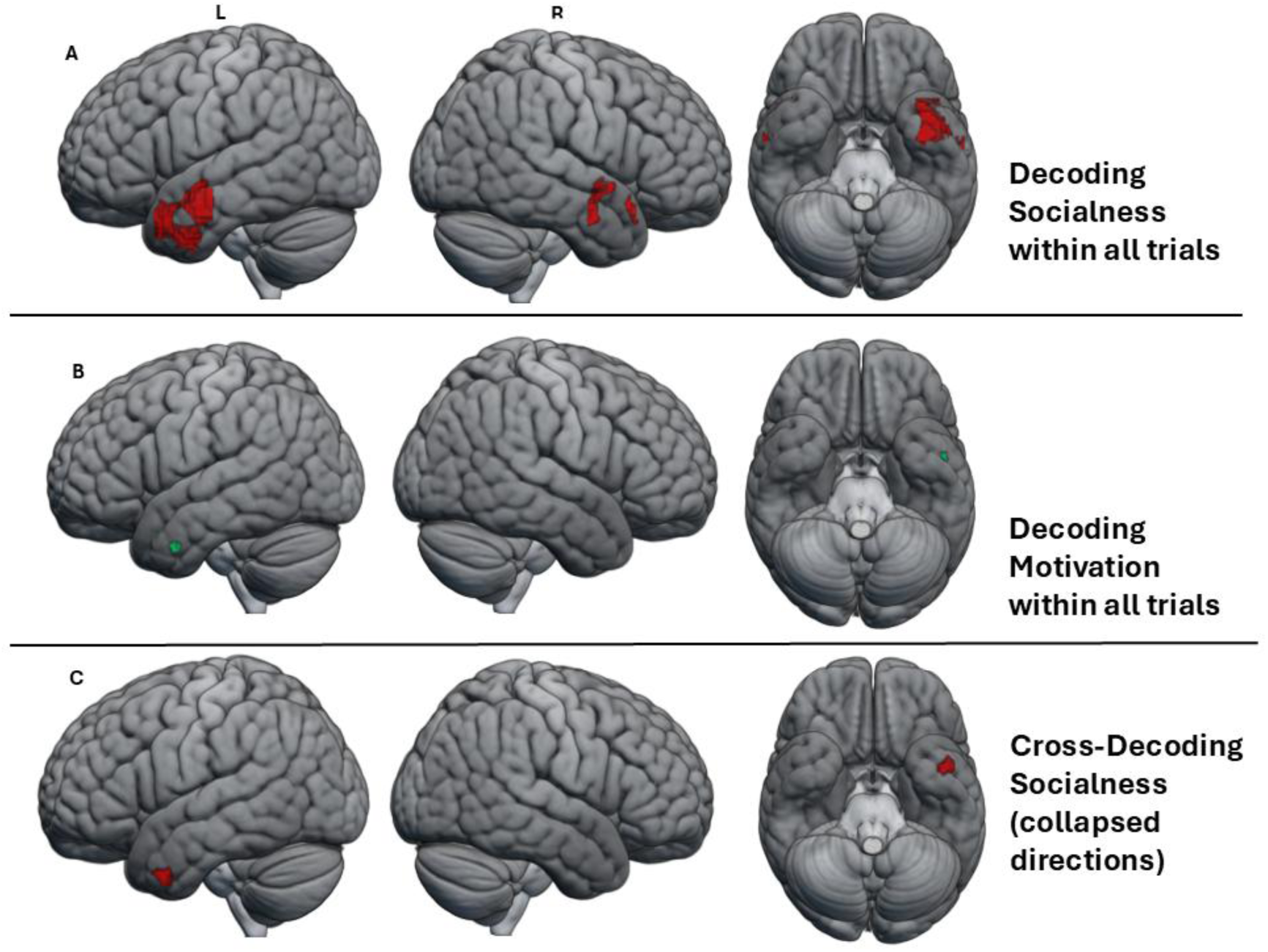
Cortical regions associated with above chance classification accuracy in the searchlight MVPA for **A**) decoding socialness within all trials, **B**) decoding motivation within all trials, **C**) cross-decoding socialness (collapsed across training directions) in the left and right hemisphere, and ventrally, respectively.

###### Cross-decoding

We examined whether local activity patterns encoding socialness in regions beyond the ATL ROIs generalize across levels of association with motivation, and vice versa. In addition to the left MTG, a group of voxels in the left ITG also showed above chance classification accuracy of socialness when collapsing across training/testing directions (see Figure 7C; see Section S5 of SM for specific training direction results). In the robustness analysis, there was no significantly above chance decoding. No ATL regions showed above chance accuracy when classifying motivation across levels of association with socialness, which remained the same in the robustness analysis (see Section S5 of SM for all cluster tables).

### 3.3. Summary and Interim Discussion

We hypothesised that the vlATL would show greater activation for each and every semantic condition relative to the baseline number task, consistent with its proposed role as an omni-category supramodal semantic hub (H1). This was confirmed in the left vlATL but the right vlATL did not respond to the semantic conditions. The contrast between semantic conditions and the baseline task also revealed activation extending throughout the canonical semantic network, from the left vlATL to the bilateral lateral ATL, left inferior frontal and left posterior temporal regions (Binder et al., 2009; Lambon Ralph et al., 2017). We tested a prediction of the *graded hub* hypothesis that the vlATL would show greater activation for semantic relative to number trials when compared to other ATL sub-regions (H2). Although the left vlATL was equally activated by all semantic conditions, this response was not greater than that observed in other ATL regions (e.g., aMTG). Moreover, regions such as the aMTG and aSTG exhibited similar response profiles, with activation across all semantic conditions. These findings suggest that the vlATL does not exhibit quantitatively stronger contributions to semantic processing than other ATL regions.

In line with the proposal that socialness makes a unique contribution to semantic representation, we predicted preferential activation for social relative to non-social concepts (H3). Indeed, social concept processing was associated with both greater activation magnitude and distinct activity patterns within the lateral ATL. Specifically, whole-brain and ROI analyses revealed greater activation for social than non-social concepts in the bilateral TP and aMTG. Moreover, TP activation appears specific to social concepts, whereas MTG was activated by all concept types. We also found some evidence of right TP selectivity for social concepts. In addition, encoding of social information in local activity patterns was observed throughout the left lateral ATL, including theleft aMTG plus the vlATL.

The sensitivity of the lateral ATL regions to the socialness of word stimuli is consistent with prior findings (Binney et al., 2016; Lin et al., 2015; Wang et al., 2019), but we extend this line of evidence in important ways. First, we demonstrate greater activation for social versus non-social concepts using a validated measure of socialness and while tightly controlling for many potentially confounding variables, such as concreteness, valence, and semantic diversity. Second, we show that the local activity patterns encoding socialness are independent of motivation-related information. Together, these findings rule out the possibility that the involvement of these regions in social concept processing is driven by motivation-related content or other semantic variables. Third, our findings appear to specifically pinpoint the left TP as the region that has a specialised role for processing social concepts. Beyond the ATL, the angular gyrus, precuneus, and medial frontal regions were more active for social concepts relative to nonsocial concepts, demonstrating engagement of brain regions implicated in social cognition more broadly (Adolphs, 2001, 2003, 2009; Carrington & Bailey, 2008; Diveica et al., 2021; Molenberghs et al., 2016; Schurz et al., 2014).

In line with the proposal that reward-related information makes a unique contribution to semantic representation, we predicted preferential activation for concepts associated with high versus low motivation (H4). We observed a main effect of motivation in posterior parahippocampal gyrus/cerebellum and the precuneus/occipital parietal sulcus. These regions have been linked to reward anticipation (e.g., Cao et al., 2019) and motivational states (Murty et al. 2016). The evidence for ATL involvement is less compelling. While high motivation trials activated vlATL and lateral ATL regions in the ROI analyses relative to the baseline NJT, they did not do so to a greater extent than trials involving low-motivation concepts, but trials with both social and motivation-related features appear to activate the bilateral ATL more than those with just social information (see Figure S2). Motivation level was encoded in the local activity patterns of the left TP and left aMTG, but these patterns were not independent of socialness. Taken together, these findings provide limited evidence for an independent contribution of motivation-related information to semantic representation in the brain. They also suggest that the effect of motivation-related semantic information on ATL activity could reflect some form of social information processing.

Lastly, we predicted that, if socialness makes contributions to word meaning that are independent to motivation-related information, this would be reflected either in additive effects (e.g., in the ATL), or in spatially non-overlapping main effects. In line with this, the whole-brain analyses revealed that the main effect of socialness implicated regions distinct from those implicated in the simple effect of motivation, suggesting a degree of independence. This independence is further supported by the finding that local activity patterns in certain ATL regions encode socialness independently of motivation level. At the same time, and as alluded to above, some results may suggest a potential interaction between the two semantic dimensions. In the whole brain analysis, simple effects of socialness in the left ATL were observed only in the high motivation condition and not the low-motivation conditions, although the ROI analysis did not reveal any significant interaction between the two factors in these regions. A simple effect of motivation on activation magnitude was observed only in social, but not non-social, trials. Moreover, the right TP ROI showed significant activation above the NJT baseline only for trials combining high social and high motivation content, but not for either dimension independently, perhaps suggesting that the right TP may respond to the variance shared between socialness and motivation dimensions.

Overall, our results suggest that socialness makes a unique contribution to semantic representation, observable in both differences in brain activation magnitude and local activity patterns. While certain brain regions are sensitive to the motivation content of concepts, this effect does not appear to be reliably independent of socialness.

## 4. General Discussion

The aim of the present study was to address the role of socialness and motivation-related information in semantic processing. Prior studies of socialness have been limited by various factors but most crucially by the lack of validated socialness norms and the degree to which they controlled for other lexical and semantic variables. Having addressed these limitations, we observed robust socialness effects in the left lateral ATL largely consistent with the results of these previous studies (Binney et al., 2016; Lin et al., 2015; Wang et al., 2019), but specifically pinpointing the left temporal pole (TP) as having a specialized role in social concept processing. Moreover, we provide the first evidence that semantic information related to motivation is represented across temporal and parietal cortex, including the left lateral ATL. We discuss the implications of these results, as well as some unexpected observations, in the sections below.

### 4.1. Implications for Multiple Representation Theories of Semantics

Multiple representation theories extend embodied accounts of semantics by proposing that conceptual knowledge is represented in many ways and is derived, not only from sensorimotor experience, but also from linguistic, emotional and other introspective experiences (Barsalou, 2020; Borghi et al., 2019). We provide new evidence to support claims that both socialness and motivation-related information should be incorporated into neurocognitive theories about the organisation of the semantic representational space (Binney et al., 2016; Diveica et al., 2024a; Giurgea et al., 2025; Lin et al., 2015; Wang et al., 2019).

Socialness emerged as an important aspect of meaning in both the behavioural and neuroimaging studies. We replicated a recently reported facilitatory effect of socialness on behaviour during semantic processing (Diveica et al., 2023, 2024b) and for the first time in the context of semantic decisions made on word triads (as opposed to single words). This was a partial replication because we only observed an effect on accuracy, and not reaction times. Facilitatory effects are consistent with a semantic richness effect whereby richer semantic representations enable faster semantic settling and/or stronger semantic activation, in turn facilitating behavioural performance (Balota et al., 1991; Pexman, 2012, 2020; Pexman et al., 2013; Tousignant & Pexman, 2012). The fact that the socialness effect occurred only in semantic judgments involving low-motivation words is consistent with there being an interaction between these semantic dimensions, rather than independence. In this case, the presence of motivation-related information could limit the influence of social information, at least in the context of the stimuli and task used here. At the brain level, we replicated prior observations of a preferential response of the left lateral ATL to social concepts (Binney et al., 2016; Lin et al., 2015; Wang et al., 2019). Beyond the ATL, social trials also elicited greater activation throughout a set of brain regions implicated in social cognition (Carrington & Bailey, 2009; Molenberghs et al., 2016). Together, our findings add to a growing body of evidence in support of the claim that social experience has an important role in shaping conceptual representation (Barsalou, 2020; Borghi et al., 2019).

It has been argued that hedonic evaluation has an important role in the representation and/or appraisal of conceptual knowledge (Giurgea et al., 2025; Rankin, 2020; Rijpma et al., 2023). Our findings provide limited evidence for a unique influence of motivation-related information during semantic decisions made on word triads. Motivation-related information did not affect accuracy or reaction times, failing to replicate the facilitatory effect previously observed in single-word tasks (Giurgea et al., 2025). This, in turn, suggests behavioural effects of motivation-related information are task-dependent (which has also been shown to be true of socialness effects; Diveica et al., 2024b). Moreover, the ventral striatum, caudate/basal ganglia, orbitofrontal and cingulate cortex are linked to the motivational aspect of reward processing (Klein-Flügge et al., 2022; Martins et al., 2021; Rogers et al., 2004; Rudebeck & Murray, 2014), but we were unable to find any evidence from among the univariate analyses that these regions are engaged by motivation-related information during semantic processing. Instead, among the main effects, we found activation in the posterior parahippocampal gyrus/cerebellum and the precuneus/occipital parietal sulcus. There is some recent evidence that has linked these regions with reward anticipation, motivational states and related functions (Cao et al., 2019; Dadario & Sughrue, 2023; Filio et al., 2026; Wang et al., 2025), but the strength of evidence in the present study was limited as the main effect was not replicated among the simple effects analyses. Therefore, overall, we provide limited support for embodied perspectives that would predict that motivation-related concepts will directly engage brain systems involved in reward processing (Barsalou et al., 2018; Borghi et al., 2019).

On the other hand, the multivariate analyses revealed encoding of motivation-related information in the left lateral ATL which suggests it is represented in supramodal semantic regions (if not directly within reward systems) during retrieval of concepts. However, the local activity patterns distinguishing high from low motivation in the ATL did not generalize across levels of socialness, and therefore are not robust. This, like the univariate analyses, also raises the question of how independent the contribution of motivation is from socialness in this region.

### 4.2. Functional Specialisation in and across the Bilateral Semantic Hub

The *graded hub* hypothesis proposes (i) that the vlATL serves as a supramodal, category- general centrepoint of a semantic hub and (ii) that regions beyond the vlATL (i.e. the TP, the MTG, STS and STG) are relatively specialised for specific types of semantic information.

The vlATL responded equivalently across all semantic conditions which is consistent with this hypothesis. Contrary to expectations, this profile was also true of the aMTG and aSTG, suggesting that, at least in terms of activation magnitude, the semantic contribution is similar across these neighbouring ATL subregions. Moreover, the aMTG exhibited the greatest magnitude of semantic activation of all regions in the left ATL, followed by aSTG, and then vlATL. This was contrary to our prediction that vlATL would show the strongest semantic activation, as observed by Binney et al. (2016). These discrepancies could be partially explained by inherently lower BOLD signal in the vlATL, compared to other ATL subregions, due to its heightened susceptibility to signal drop-out. We used a dual-echo EPI acquisition and distortion correction technique which improves overall signal quality in this region (Embleton et al., 2010; Balgova et al., 2022), but the contrast-to-noise ratio remains nonetheless impoverished in the vlATL relative to the aMTG (Halai et al., 2025; see Table S35). Moreover, the unique characteristics of vlATL function are perhaps unlikely to be reflected in response magnitude, and may be more apparent in the temporal dynamics of activation and connectivity (c.f., Bajada et al., 2019; Rogers et al., 2021).

We also found evidence for functional distinctions across the lateral ATL, as hypothesised. We replicated prior observations of a preferential response of the left lateral ATL to social concepts (Binney et al., 2016; Lin et al., 2015; Wang et al., 2019). In these previous studies, greater activation to social relative to non-social concepts has been observed in different lateral ATL sub-regions, including the TP, aMTG, and aSTG. Having used a recently validated measure of socialness and tightly controlling for a range of lexical and semantic variables, our study revealed greater and seemingly selective responses to social trials specifically in the left TP. Left aMTG responses were also greater in magnitude for social relative to non-social trials but this region activated for all semantic stimuli which suggests it plays a more category-general semantic function than the TP. To the best of our knowledge, this was the first neuroimaging study of social concepts to use not only univariate approaches but also MVPA, and these latter analyses provide further support for an important role for the left TP and aMTG in representing social semantic information. However, only local activity patterns in the aMTG were able to distinguish between social and non-social trials in a way that generalized across levels of motivation. Together, these findings provide the most compelling evidence to date for claims that that there are partially distinct neural bases for social relative to non-social semantic knowledge (Binney et al., 2016), but they also reveal novel insights into distinct functional profiles among the adjacent left TP and aMTG subregions. We argue that the response profile of the left TP is consistent with a specialised role in processing social semantic information (Binney et al., 2016; Lin et al., 2018), and we differentiate this role from that of the aMTG which is complementary and sensitive to socialness, but not domain specific (see Section 4.3 for more detailed discussion).

We also provide evidence that the right TP is selectively engaged by socialness. Indeed, we observed two key distinctions between left and right ATL activity during the semantic task. First, the left ATL exhibited overall stronger responses than the right ATL in both social and non-social semantic conditions, perhaps due to the verbal nature of the stimuli and preferential connectivity of the left ATL to the language production system (Lambon Ralph et al., 2001; Rice et al., 2016). Second, the weaker, right-lateralised ATL activation was limited to dorsolateral regions. This pattern is consistent with many of the previous neuroimaging studies of social concept processing (Binney et al., 2016; Ross & Olson, 2010; Zahn et al., 2007), and we have previously interpreted it in support of there being a dominant role of the left ATL in representing social semantics (Binney et al., 2016; Balgova et al., 2022, 2024; also see complementary evidence from Rice et al, 2015; Rouse, Halai et al, 2024). This perspective contrasts with proposals stemming from patient studies of semantic dementia, where it is argued that socio-emotional semantic processing relies mostly on the right ATL (Irish et al., 2013; Younes et al., 2022; also see Gainotti, 2015). The functional contributions of left ATL subregions to general semantics are overall better understood than those in the right, and what it is that right ATL subregions could be contributing to social semantic processing is far from clear, and beyond the scope of the present study.

Interestingly, though, in the present study the right TP was only significantly activated above baseline during trials containing both social and motivation-related semantic content, raising the possibility that this region is sensitive to the shared variance between the two dimensions. It has been proposed that social concepts are more likely than non-social concepts to have hedonic evaluations contribute to their meaning (Arioli et al., 2021; Rankin, 2020), and this particular finding may reflect a role of the right TP in integrating motivation computed by frontal regions into semantic representations (c.f. Rankin, 2020). Further research is needed to test this hypothesis directly.

### 4.3. Socialness and the Default Mode Network

Social concepts elicited activation not just within the ATL, but also across a wider distributed set of regions associated with social cognition, including the angular gyrus, medial prefrontal cortex, and precuneus (Carrington & Bailey, 2009; Molenberghs et al., 2016). This is consistent with observations of Lin et al. (2015) who found that social action verbs (*salute)* activated the Theory of Mind network when contrasted with private action verbs (*walk*) and non-human verbs (*drip*). It is increasingly recognised that semantic and social cognitive processing engage an overlapping set of brain regions (Balgova et al., 2024; Diveica et al., 2021). From the perspective of multiple representation theories, the greater involvement of these regions in response to socially-relevant concepts could suggest reliance on social cognitive processing in support of social conceptual representation (Borghi et al., 2019; 2025; Lin et al., 2018). However, the functional contribution of these regions remains debated. The main locus of this convergence corresponds to the default mode network (DMN), a large-scale functional network that has been more broadly implicated in internally driven cognitive states, including, but not limited to, social cognition (Andrews-Hanna et al., 2014; Menon, 2023; Smallwood et al., 2021; Spreng & Andrews-Hanna, 2015). The DMN’s position at the transmodal end (as opposed to sensorimotor end) of the ‘principal gradient’ of functional organisation of the cortex (Margulies et al., 2016; Smallwood et al., 2021) makes it ideally suited for the generation and updating of mental models of the world via the integration of prior (semantic) knowledge with external perceptual inputs (Barnett & Bellana, 2025; Fernandino & Binder, 2024).

It is possible that the role of aMTG in social concept processing, which was one of only two ATL regions encoding socialness independently from motivation, reflects its affiliation with the DMN. In line with this, Lin et al. (2020) demonstrated that the aMTG region that responded preferentially to social as opposed to non-social concepts was functionally connected with other DMN regions. Relatedly, using a similar definition of socialness as the current investigation (Diveica et al., 2023), Thye et al. (2024) showed that high relative to low social content in narratives drives greater engagement of the aMTG. Moreover, they observed similar engagement of aMTG and other DMN regions in support of other aspects of semantic narrative content, which suggests a more general role of the aMTG in situating incoming semantic information in context to facilitate comprehension, rather than a role in engaging social cognitive processes per se. This appears consistent with the omni-category response of the aMTG in the present study. Thye et al. (2025) also found that the functional connectivity between aMTG and other DMN regions covaries with general semantic richness rather than social semantic content specifically during narrative moments in a movie-watching paradigm. These previous findings suggest that, by virtue of its affiliation with the DMN, the aMTG might play a key role in contextualizing meaningful inputs of all kinds through integration in situation models (Thye et al., 2024, 2025). While the current findings do not directly test this proposal, they highlight the aMTG (particularly in the left hemisphere) as a key contributor to semantic processing, exhibiting a distinct functional profile among ATL ROIs. Specifically, we found that this region encodes both socialness and motivation-related semantic content, while also showing stronger activation than other ATL regions across all semantic conditions relative to the NJT baseline.

### 4.4. Conclusion

The aim of the present study was to re-examine socialness effects in semantic processing on behaviour and on neural responses in the lateral ATL. Moreover, we sought to untangle the unique contributions of socialness and motivation-related information to conceptual representations. Alongside our previous studies, the present findings demonstrate that socialness (Diveica et al., 2024b) and motivation (Giurgea et al., 2025) effects are task-dependent at the behavioural level. However, in the case of socialness at least, the effects are robust at the neural level. Overall, this supports the claim that socialness is an important semantic dimension (Pexman et al., 2023). The present study also provides novel insights into the functional organisation of the lateral ATL; having used a new set of socialness norms (Diveica et al., 2023) to carefully manipulate the social content of stimuli while maintaining unprecedented control of other semantic variables, we found new evidence of two regions with distinct functional profiles. First, the left TP appears to be selective for social concepts relative to non-social concepts. Second, the left aMTG exhibits a more general semantic function which is not limited to social concepts but does appear to be particularly important for understanding those kinds of concepts. Finally, we demonstrated that motivation-related semantic information is represented across temporal and parietal cortex but it is not independent of social semantic content. Overall, these present findings align with multiple representation theories and further constrain neurocognitive theories of conceptual knowledge (Lambon Ralph et al., 2017; Patterson et al., 2007) and social semantics (Binney & Ramsey, 2020; Pexman et al., 2023). Further work is needed to map out the graded organization of the ATL semantic hub and clarify the ways in which information derived from social experiences and motivated cognition interact to shape semantic representations.

## Contributions

Doina-Irina Giurgea: Conceptualization, Methodology, Script Writing, Formal Analysis, Investigation, Writing – Original Draft, Writing – Review and Editing, Visualisation, Project Administration

Veronica Diveica: Conceptualization, Methodology, Script Writing, Writing – Review and Editing, Visualisation

Penny M. Pexman: Conceptualization, Methodology, Writing – Review and Editing Richard J. Binney: Conceptualization, Methodology, Resources, Script Writing, Writing – Original Draft, Writing – Review and Editing, Supervision

## Acknowledgements

The authors thank Rigerta Disha, Rija Fatima, Cinthya Luna Cuevas, and Rashmil Tufail for their help with participant recruitment, and Andrew Fischer, Ferrida Ponce, and Deyan Mitev for assisting with data acquisition.

## Data availability statement

All data and materials are available on the OSF project page (https://osf.io/r5dj6/).

## Notes

https://osf.io/r5dj6/

### Competing Interest Statement

The authors have declared no competing interest.

