## Supplemental Information for "Social Experience and Motivated Cognition Shape the Representation of Concepts: A Behavioural and Functional Neuroimaging Study"

for

### Section S1. Additional Word Stimuli Information

**Table S1.** *Frequencies of parts of speech by word condition for probes*

| Condition | Adjective | Noun | Verb | Total |
|---|---|---|---|---|
| Non-social Low Motivation | 16 | 38 | 6 | 60 |
| Non-social High Motivation | 16 | 38 | 6 | 60 |
| Social Low Motivation | 17 | 37 | 6 | 60 |
| Social High Motivation | 16 | 38 | 6 | 60 |
| Total | 65 | 151 | 24 | 240 |

**Table S2.** *Wilcoxon test results for motivation ratings of probes*

| IV | Group 1 | Group 2 | Statistic | P-value | Adjusted p-value |
|---|---|---|---|---|---|
| Motivation | A1_B1 | A2_B1 | 1,456.50 | 0.07 | 0.14 |
| | A1_B2 | A2_B2 | 1,722.50 | 0.69 | 0.69 |
| | A1_B1 | A1_B2 | 0.00 | 0.00 | 0.00 |
| | A2_B1 | A2_B2 | 0.00 | 0.00 | 0.00 |

*Note.* A1B1 = low socialness low motivation; A1B2 = low socialness high motivation; A2B1 = high socialness low motivation; A2B2 = high socialness high motivation

**Table S3.** *Wilcoxon test results for socialness ratings of probes*

| IV | Group 1 | Group 2 | Statistic | P-value | Adjusted p-value |
|---|---|---|---|---|---|
| Socialness | A1_B1 | A2_B1 | 0.00 | 0.00 | 0.00 |
| | A1_B2 | A2_B2 | 0.00 | 0.00 | 0.00 |
| | A1_B1 | A1_B2 | 1,539.50 | 0.17 | 0.34 |
| | A2_B1 | A2_B2 | 1,752.50 | 0.80 | 0.80 |

*Note.* A1B1 = low socialness low motivation; A1B2 = low socialness high motivation; A2B1 = high socialness low motivation; A2B2 = high socialness high motivation

**Table S4.** *Wilcoxon test results for all other variables for probes*

| IV | Group 1 | Group 2 | Statistic | P-value | Adjusted p-value |
|---|---|---|---|---|---|
| Length | A1_B1 | A2_B1 | 1,696.00 | 0.58 | 1.00 |
| | A1_B2 | A2_B2 | 1,679.50 | 0.52 | 1.00 |
| | A1_B1 | A1_B2 | 1,630.50 | 0.37 | 1.00 |
| | A2_B1 | A2_B2 | 1,612.50 | 0.32 | 1.00 |
| Frequency | A1_B1 | A2_B1 | 1,818.00 | 0.93 | 1.00 |
| | A1_B2 | A2_B2 | 1,728.50 | 0.71 | 1.00 |
| | A1_B1 | A1_B2 | 1,762.00 | 0.84 | 1.00 |
| | A2_B1 | A2_B2 | 1,625.50 | 0.36 | 1.00 |
| AoA.Rating | A1_B1 | A2_B1 | 1,694.00 | 0.58 | 1.00 |

| IV | Group 1 | Group 2 | Statistic | P-value | Adjusted p-value |
|---|---|---|---|---|---|
| | A1_B2 | A2_B2 | 1,592.50 | 0.28 | 1.00 |
| | A1_B1 | A1_B2 | 1,773.00 | 0.89 | 1.00 |
| | A2_B1 | A2_B2 | 1,683.00 | 0.54 | 1.00 |
| Concreteness | A1_B1 | A2_B1 | 1,780.00 | 0.92 | 1.00 |
| | A1_B2 | A2_B2 | 1,978.50 | 0.35 | 1.00 |
| | A1_B1 | A1_B2 | 1,879.00 | 0.68 | 1.00 |
| | A2_B1 | A2_B2 | 2,138.00 | 0.08 | 0.31 |
| Valence | A1_B1 | A2_B1 | 1,989.50 | 0.32 | 1.00 |
| | A1_B2 | A2_B2 | 2,104.50 | 0.11 | 0.44 |
| | A1_B1 | A1_B2 | 1,470.50 | 0.08 | 0.34 |
| | A2_B1 | A2_B2 | 1,601.50 | 0.30 | 1.00 |
| Semantic Diversity | A1_B1 | A2_B1 | 1,832.50 | 0.62 | 1.00 |
| | A1_B2 | A2_B2 | 2,160.00 | 0.06 | 0.24 |
| | A1_B1 | A1_B2 | 1,484.50 | 0.13 | 0.52 |
| | A2_B1 | A2_B2 | 1,689.50 | 0.67 | 1.00 |

*Note.* A1B1 = low socialness low motivation; A1B2 = low socialness high motivation; A2B1 = high socialness low motivation; A2B2 = high socialness high motivation

**Table S5.** *Wilcoxon test results for motivation ratings of targets*

| IV | Group 1 | Group 2 | Statistic | P-value | Adjusted p-value |
|---|---|---|---|---|---|
| Motivation | A1_B1_pair | A2_B1_pair | 1,518.00 | 0.14 | 0.56 |
| | A1_B2_pair | A2_B2_pair | 1,587.50 | 0.27 | 1.00 |
| | A1_B1_pair | A1_B2_pair | 4.00 | 0.00 | 0.00 |
| | A2_B1_pair | A2_B2_pair | 0.00 | 0.00 | 0.00 |

*Note.* A1B1 = low socialness low motivation; A1B2 = low socialness high motivation; A2B1 = high socialness low motivation; A2B2 = high socialness high motivation; IV = independent variable

**Table S6.** *Wilcoxon test results for socialness ratings of targets*

| IV | Group 1 | Group 2 | Statistic | P-value | Adjusted p-value |
|---|---|---|---|---|---|
| Socialness | A1_B1_pair | A2_B1_pair | 0.00 | 0.00 | 0.00 |
| | A1_B2_pair | A2_B2_pair | 0.00 | 0.00 | 0.00 |
| | A1_B1_pair | A1_B2_pair | 1,442.50 | 0.06 | 0.24 |
| | A2_B1_pair | A2_B2_pair | 1,441.00 | 0.06 | 0.24 |

*Note.* A1B1 = low socialness low motivation; A1B2 = low socialness high motivation; A2B1 = high socialness low motivation; A2B2 = high socialness high motivation; IV = independent

**Table S7.** *Wilcoxon test results for all other variables for targets*

| IV | Group 1 | Group 2 | Statistic | P-value | Adjusted p-value |
|---|---|---|---|---|---|
| Length | A1_B1_pair | A2_B1_pair | 1,535.50 | 0.16 | 0.64 |
| | A1_B2_pair | A2_B2_pair | 1,446.00 | 0.06 | 0.25 |
| | A1_B1_pair | A1_B2_pair | 1,683.00 | 0.54 | 1.00 |
| | A2_B1_pair | A2_B2_pair | 1,565.50 | 0.22 | 0.86 |
| Frequency | A1_B1_pair | A2_B1_pair | 1,921.50 | 0.52 | 1.00 |
| | A1_B2_pair | A2_B2_pair | 1,843.00 | 0.70 | 1.00 |
| | A1_B1_pair | A1_B2_pair | 1,683.50 | 0.54 | 1.00 |
| | A2_B1_pair | A2_B2_pair | 1,489.00 | 0.14 | 0.54 |
| AoA.Rating | A1_B1_pair | A2_B1_pair | 1,655.50 | 0.45 | 1.00 |
| | A1_B2_pair | A2_B2_pair | 1,273.00 | 0.01 | 0.02 |
| | A1_B1_pair | A1_B2_pair | 1,982.00 | 0.34 | 1.00 |
| | A2_B1_pair | A2_B2_pair | 1,561.50 | 0.21 | 0.85 |
| Concreteness | A1_B1_pair | A2_B1_pair | 1,807.50 | 0.97 | 1.00 |
| | A1_B2_pair | A2_B2_pair | 1,917.00 | 0.54 | 1.00 |
| | A1_B1_pair | A1_B2_pair | 1,955.00 | 0.42 | 1.00 |
| | A2_B1_pair | A2_B2_pair | 2,168.00 | 0.05 | 0.22 |
| Valence | A1_B1_pair | A2_B1_pair | 1,747.00 | 0.78 | 1.00 |
| | A1_B2_pair | A2_B2_pair | 1,821.50 | 0.91 | 1.00 |
| | A1_B1_pair | A1_B2_pair | 1,490.50 | 0.10 | 0.42 |
| | A2_B1_pair | A2_B2_pair | 1,611.00 | 0.32 | 1.00 |
| Semantic Diversity | A1_B1_pair | A2_B1_pair | 2,063.50 | 0.01 | 0.05 |
| | A1_B2_pair | A2_B2_pair | 2,167.50 | 0.00 | 0.01 |
| | A1_B1_pair | A1_B2_pair | 1,436.00 | 0.08 | 0.30 |
| | A2_B1_pair | A2_B2_pair | 1,163.00 | 0.05 | 0.20 |

*Note.* A1B1 = low socialness low motivation; A1B2 = low socialness high motivation; A2B1 = high socialness low motivation; A2B2 = high socialness high motivation; IV = independent variable

**Table S8.** *Wilcoxon test results for motivation ratings of distractors*

| IV | Group 1 | Group 2 | Statistic | P-value | Adjusted p-value |
|---|---|---|---|---|---|
| Motivation | A1_B1_distractor | A2_B1_distractor | 1,445.50 | 0.06 | 0.25 |
| | A1_B2_distractor | A2_B2_distractor | 1,513.50 | 0.13 | 0.53 |
| | A1_B1_distractor | A1_B2_distractor | 0.00 | 0.00 | 0.00 |
| | A2_B1_distractor | A2_B2_distractor | 0.00 | 0.00 | 0.00 |

*Note.* A1B1 = low socialness low motivation; A1B2 = low socialness high motivation; A2B1 = high socialness low motivation; A2B2 = high socialness high motivation; IV = independent variable

**Table S9.** *Wilcoxon test results for socialness ratings of distractors*

| IV | Group 1 | Group 2 | Statistic | P-value | Adjusted p-value |
|---|---|---|---|---|---|
| Socialness | A1_B1_distractor | A2_B1_distractor | 0.00 | 0.00 | 0.00 |
| | A1_B2_distractor | A2_B2_distractor | 0.00 | 0.00 | 0.00 |
| | A1_B1_distractor | A1_B2_distractor | 1,528.50 | 0.16 | 0.62 |
| | A2_B1_distractor | A2_B2_distractor | 1,656.00 | 0.45 | 1.00 |

*Note.* A1B1 = low socialness low motivation; A1B2 = low socialness high motivation; A2B1 = high socialness low motivation; A2B2 = high socialness high motivation; IV = independent variable

**Table S10.** *Wilcoxon test results for all other variables for distractors*

| IV | Group 1 | Group 2 | Statistic | P-value | Adjusted p-value |
|---|---|---|---|---|---|
| Length | A1_B1_distractor | A2_B1_distractor | 1,575.50 | 0.23 | 0.93 |
| | A1_B2_distractor | A2_B2_distractor | 1,374.00 | 0.02 | 0.09 |
| | A1_B1_distractor | A1_B2_distractor | 1,617.50 | 0.33 | 1.00 |
| | A2_B1_distractor | A2_B2_distractor | 1,460.50 | 0.07 | 0.29 |
| Frequency | A1_B1_distractor | A2_B1_distractor | 1,846.00 | 0.69 | 1.00 |
| | A1_B2_distractor | A2_B2_distractor | 1,713.00 | 0.65 | 1.00 |
| | A1_B1_distractor | A1_B2_distractor | 1,577.00 | 0.24 | 0.97 |
| | A2_B1_distractor | A2_B2_distractor | 1,421.00 | 0.06 | 0.26 |
| AoA.Rating | A1_B1_distractor | A2_B1_distractor | 1,716.00 | 0.78 | 1.00 |
| | A1_B2_distractor | A2_B2_distractor | 1,900.50 | 0.60 | 1.00 |
| | A1_B1_distractor | A1_B2_distractor | 1,842.00 | 0.83 | 1.00 |
| | A2_B1_distractor | A2_B2_distractor | 1,934.50 | 0.38 | 1.00 |
| Concreteness | A1_B1_distractor | A2_B1_distractor | 1,608.50 | 0.32 | 1.00 |
| | A1_B2_distractor | A2_B2_distractor | 1,760.00 | 0.84 | 1.00 |
| | A1_B1_distractor | A1_B2_distractor | 1,934.00 | 0.48 | 1.00 |
| | A2_B1_distractor | A2_B2_distractor | 2,109.50 | 0.10 | 0.42 |
| Valence | A1_B1_distractor | A2_B1_distractor | 1,782.50 | 0.93 | 1.00 |
| | A1_B2_distractor | A2_B2_distractor | 1,862.50 | 0.74 | 1.00 |
| | A1_B1_distractor | A1_B2_distractor | 1,460.50 | 0.08 | 0.30 |
| | A2_B1_distractor | A2_B2_distractor | 1,651.50 | 0.44 | 1.00 |
| Semantic Diversity | A1_B1_distractor | A2_B1_distractor | 1,939.50 | 0.11 | 0.44 |
| | A1_B2_distractor | A2_B2_distractor | 1,838.50 | 0.72 | 1.00 |
| | A1_B1_distractor | A1_B2_distractor | 1,905.00 | 0.38 | 1.00 |
| | A2_B1_distractor | A2_B2_distractor | 1,579.00 | 0.57 | 1.00 |

*Note.* A1B1 = low socialness low motivation; A1B2 = low socialness high motivation; A2B1 = high socialness low motivation; A2B2 = high socialness high motivation; IV = independent variable

**Table S11.** *Wilcoxon test results for mean reverse cosine distance between probes and targets and probes and distractors*

| IV | Group 1 | Group 2 | Statistic | P-value | Adjusted p-value |
|---|---|---|---|---|---|
| Reverse Cosine Distance | A1B1 | A1B1_pd | 775.00 | 0.00 | 0.00 |
| | A1B2 | A1B2_pd | 734.00 | 0.00 | 0.00 |
| | A2B1 | A2B1_pd | 612.00 | 0.00 | 0.00 |
| | A2B2 | A2B2_pd | 724.00 | 0.00 | 0.00 |

*Note.* A1B1 = low socialness low motivation; A1B2 = low socialness high motivation; A2B1 = high socialness low motivation; A2B2 = high socialness high motivation. "_pd" refers to the values for the mean reverse cosine distance between probes and distractors. IV = independent variable

**Table S12.** *Wilcoxon test results for mean reverse cosine distance between probes and targets*

| IV | Group 1 | Group 2 | Statistic | P-value | Adjusted p-value |
|---|---|---|---|---|---|
| Reverse Cosine Distance | A1B1 | A2B1 | 1,966.00 | 0.38 | 1.00 |
| | A1B2 | A2B2 | 2,071.00 | 0.16 | 0.62 |
| | A1B1 | A1B2 | 1,777.00 | 0.91 | 1.00 |
| | A2B1 | A2B2 | 1,868.00 | 0.72 | 1.00 |

*Note.* A1B1 = low socialness low motivation; A1B2 = low socialness high motivation; A2B1 = high socialness low motivation; A2B2 = high socialness high motivation; IV = independent variable

**Table S13.** *Wilcoxon test results for mean reverse cosine distance between probes and distractors*

| IV | Group 1 | Group 2 | Statistic | P-value | Adjusted p-value |
|---|---|---|---|---|---|
| Reverse Cosine Distance | A1B1_pd | A2B1_pd | 1,984.00 | 0.34 | 1.00 |
| | A1B2_pd | A2B2_pd | 2,096.00 | 0.12 | 0.48 |
| | A1B1_pd | A1B2_pd | 1,775.00 | 0.90 | 1.00 |
| | A2B1_pd | A2B2_pd | 1,902.00 | 0.59 | 1.00 |

*Note.* A1B1 = low socialness low motivation; A1B2 = low socialness high motivation; A2B1 = high socialness low motivation; A2B2 = high socialness high motivation; IV = independent variable

**Table S14.** *Mean reverse cosine distances for all conditions*

| Condition | probes - targets | probes - distractors |
|---|---|---|
| A1B1 | 0.04 | 0.08 |
| A1B2 | 0.04 | 0.08 |
| A2B1 | 0.03 | 0.07 |
| A2B2 | 0.03 | 0.07 |

*Note.* A1B1 = low socialness low motivation; A1B2 = low socialness high motivation; A2B1 = high socialness low motivation; A2B2 = high socialness high motivation; IV = independent variable

### Section S2. Additional Results for Behavioural Online Experiment

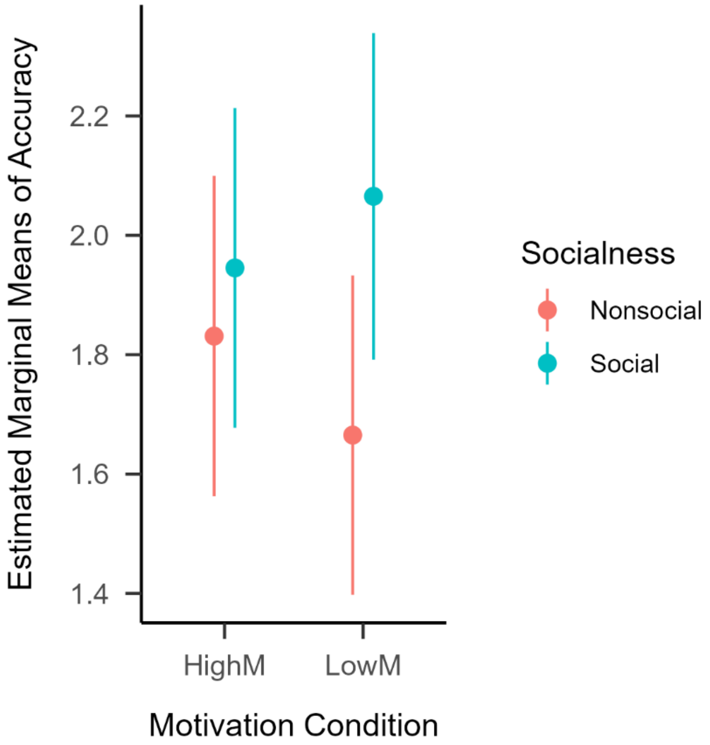

***Figure S1.*** Estimated marginal means for the socialness simple effects between levels of motivation.

**Section S3. fMRI Experiment Behavioural Data**

The descriptive statistics for both tasks are presented in Table S16. We performed pairwise Wilcoxon tests and found no significant differences between SJT condition on RT or accuracy. There were also no significant differences between the SJT conditions and the NJT. The raw RT data did not meet the assumption of normality, and, thus, we conducted all analyses on log transformed RTs (see Table 4 for mean RT and accuracy scores for each condition; see SM for individual participant accuracy scores). The model with the optimal random effects structure for our data included one random intercept per participant and one random intercept per word. This was implemented using the following formula: logRT ~ Socialness*Motivation + (1|Participant) + (1|Word). The main effects of socialness and motivation and their interaction were nonsignificant (see Table S17).

The model with the optimal random effects structure for the accuracy data also included one random intercept per participant and one random intercept per word: Accuracy ~ Socialness*Motivation + (1| Participant) + (1|Word). The main effects of socialness and motivation and their interaction were nonsignificant (see Table S17).

**Table S15.** *Descriptive statistics for the SJT by condition*

| Condition | Total number of trials | Mean RT (SD) in milliseconds | Number of accurate trials | Mean RT (SD) in milliseconds for accurate trials | Mean Accuracy % (SD) |
|---|---|---|---|---|---|
| Non-social Low Motivation | 1800 | 1388 (330) | 1496 | 1350 (317) | 83.11 (7.8) |
| Non-social High Motivation | 1800 | 1359 (320) | 1525 | 1329 (308) | 84.72 (6.7) |
| Social Low Motivation | 1800 | 1350 (323) | 1559 | 1322 (312) | 86.61 (8.8) |
| Social High Motivation | 1800 | 1367 (306) | 1534 | 1347 (301) | 85.22 (7.9) |
| Number | 3840 | 1372 (293) | 3304 | 1363 (285) | 86.04 (7.55) |

**Table S16.** *Mixed effects models predicting SJT response time and accuracy*

| | log RT | | | Accuracy | | |
|---|---|---|---|---|---|---|
| *Predictors* | *Estimates* | *CI* | *p* | *Odds Ratios* | *CI* | *p* |
| (Intercept) | 0.12 | 0.10 – 0.14 | **<0.001** | 8.89 | 6.84 – 11.55 | **<0.001** |
| Socialness | 0.00 | -0.01 – 0.02 | 0.623 | 0.79 | 0.57 – 1.08 | 0.137 |
| Motivation | 0.01 | -0.01 – 0.03 | 0.231 | 1.06 | 0.77 – 1.45 | 0.726 |
| Socialness x Motivation | -0.02 | -0.04 – 0.01 | 0.139 | 0.69 | 0.37 – 1.29 | 0.240 |
| **Random Effects** | | | | | | |
| $\sigma^2$ | 0.01 | | | 3.29 | | |
| $\tau_{00\ Word}$ | 0.00 | | | 1.17 | | |
| $\tau_{00\ participant}$ | 0.00 | | | 0.32 | | |
| ICC | 0.40 | | | 0.31 | | |
| $N_{participant}$ | 30 | | | 30 | | |
| $N_{Word}$ | 240 | | | 240 | | |
| Observations | 6943 | | | 7200 | | |
| Marginal $R^2$ / Conditional $R^2$ | 0.002 / 0.404 | | | 0.005 / 0.315 | | |

Note. CI = confidence interval; Socialness is a factor variable with social words set as the reference group. Motivation is a factor variable with high motivation set as the reference group. Accuracy is a binary dependent variable with inaccurate responses as the reference group (0) and accurate responses as the focus group (1). The marginal $R^2$ includes only the variance from the fixed effects and the conditional $R^2$ includes variance from both the fixed and random effects.

**Section S4. Additional Univariate Analyses Results**

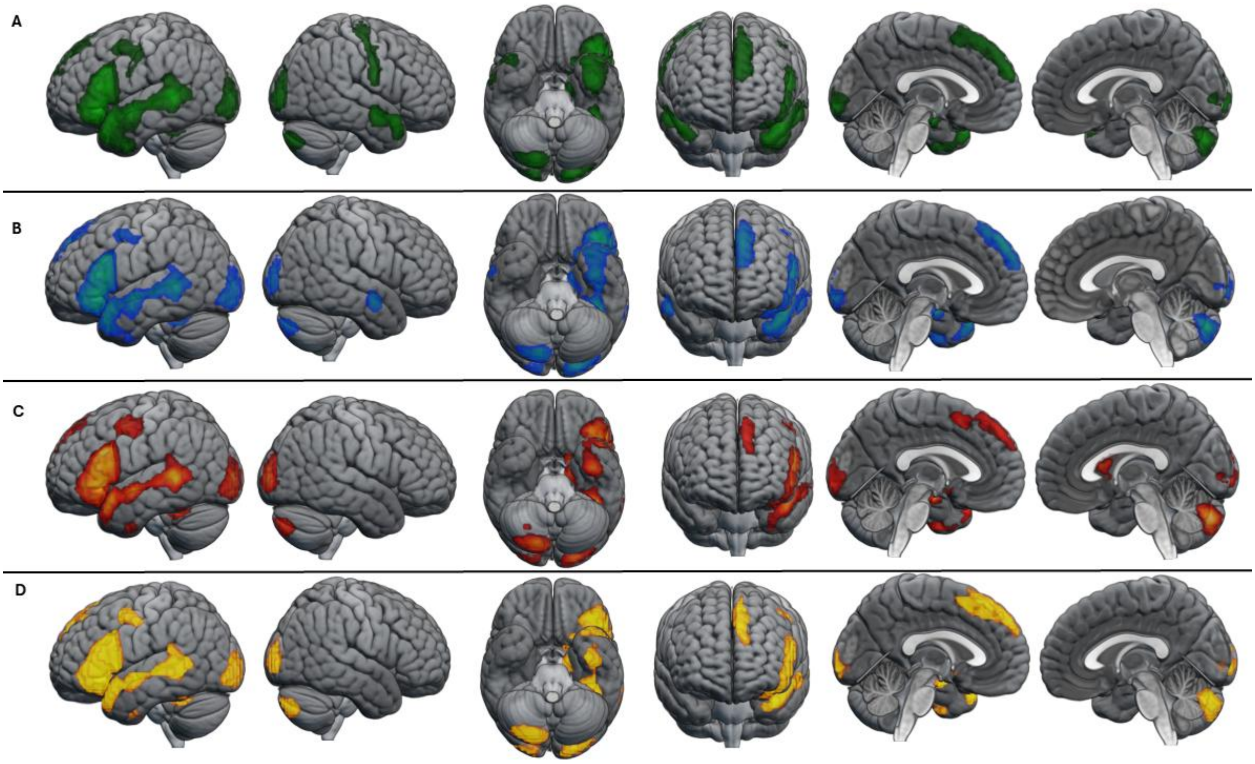

***Figure S2.*** Cortical regions activated by the A) social high motivation condition relative to the number judgement condition, B) social low motivation condition relative to the number judgement condition, C) non-social high motivation condition relative to the number judgement condition, D) non-social low motivation condition relative to the number judgement condition. The statistical map is thresholded with a voxel-height threshold of $p$ <.001, uncorrected and FWE corrected at $p < 0.05$.

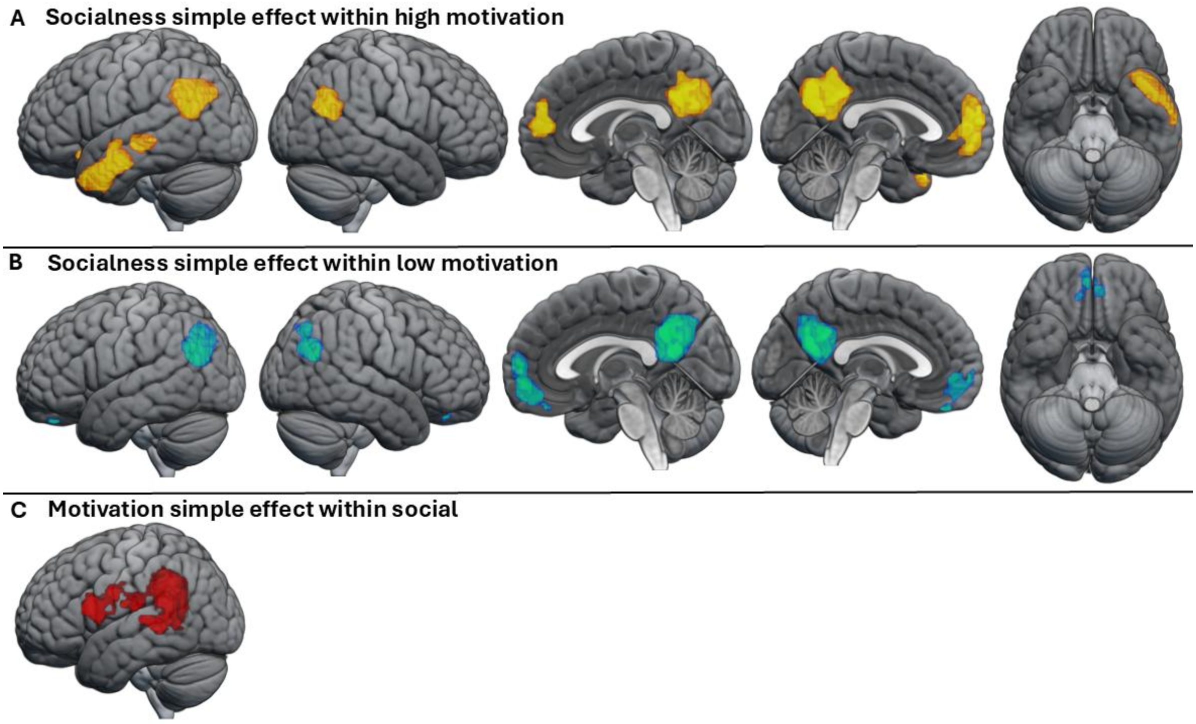

***Figure S3.*** Cortical regions activated by the A) socialness simple effect within high motivation, B) socialness simple effect within low motivation, C) motivation simple effects within social. The statistical map is thresholded with a voxel-height threshold of $p$ <.001, uncorrected and FWE corrected.

**Table S17.** *Significant activation clusters in whole-brain analysis*

| Contrast | Cluster name and location of maximum | Cluster size (voxels) | Cluster extent threshold | Peak (Z) | MNI maxima coordinates (mm) | | |
|---|---|---|---|---|---|---|---|
| | | | | | x | y | z |
| **Social High Motivation > NJT** | **L Fronto-Temporal Cortex** | **3264** | 143 | | | | |
| | Middle temporal gyrus | | | 7.11 | -51 | -34 | 2 |
| | Middle temporal gyrus | | | 6.87 | -60 | -43 | 11 |
| | Inferior frontal gyrus (pars triangularis) | | | 6.75 | -48 | 20 | 26 |

| | | | | | |
|---|---|---|---|---|---|
| Inferior frontal gyrus (pars orbitalis) | | 6.72 | -42 | 29 | -7 |
| Middle temporal gyrus | | 6.55 | -54 | -7 | -13 |
| Middle temporal gyrus | | 6.52 | -60 | -46 | 2 |
| Inferior frontal gyrus (pars triangularis) | | 6.48 | -51 | 26 | 14 |
| Inferior frontal gyrus (pars opercularis) | | 6.42 | -54 | 17 | 14 |
| **R Cerebellum** | **406** | | | | |
| Crus II | | 7.02 | 18 | -79 | -37 |
| **L Occipital Lobe** | **639** | | | | |
| Middle occipital gyrus | | 6.96 | -15 | -91 | -7 |
| Fusiform gyrus | | 6.65 | -42 | -43 | -22 |
| Middle occipital gyrus | | 6.45 | -24 | -97 | 2 |
| Middle occipital gyrus | | 5.22 | -27 | -97 | 14 |
| Inferior occipital gyrus | | 3.52 | -42 | -73 | -13 |
| **R Temporal Lobe** | **262** | | | | |
| Calcarine | | 6.65 | 18 | -94 | -4 |
| Superior occipital gyrus | | 4.86 | 27 | -94 | 20 |
| Middle occipital gyrus | | 3.78 | 36 | -91 | 11 |
| **R Parietal Lobe** | **143** | | | | |
| Postcentral gyrus | | 5.38 | 66 | -4 | 26 |
| Postcentral gyrus | | 5.35 | 63 | -7 | 38 |
| Precentral gyrus | | 4.75 | 48 | -13 | 59 |

| | | | | | | | |
|---|---|---|---|---|---|---|---|
| | Precentral gyrus | | | 4.45 | 42 | -13 | 65 |
| | Precentral gyrus | | | 4.42 | 54 | -10 | 50 |
| | Precentral gyrus | | | 3.49 | 27 | -19 | 74 |
| | **R Superior Temporal Cortex** | **175** | | | | | |
| | Heschl's gyrus | | | 5.37 | 42 | -25 | 14 |
| | Heschl's gyrus | | | 5.18 | 42 | -19 | 8 |
| | **R Superior Temporal Pole** | **306** | | | | | |
| | Superior temporal pole | | | 5.34 | 48 | 14 | -19 |
| | Middle temporal pole | | | 5.23 | 57 | -4 | -13 |
| | **L Frontal Lobe** | **356** | | | | | |
| | Superior frontal gyrus | | | 5.22 | -12 | 56 | 29 |
| | Medial superior frontal gyrus | | | 5.13 | -9 | 50 | 41 |
| | Superior frontal gyrus | | | 5.04 | -12 | 38 | 47 |
| | Medial superior frontal gyrus | | | 4.92 | -6 | 29 | 53 |
| | Supplementary Motor Area | | | 4.12 | -9 | 20 | 62 |
| **Social Low Motivation > NJT** | **R Cerebellum** | **365** | 109 | | | | |
| | Crus II | | | 6.98 | 18 | -79 | -37 |
| | **L Temporal Occipital Cortex** | **3057** | | | | | |
| | Middle temporal gyrus | | | 6.82 | -51 | -34 | 2 |
| | Fusiform gyrus | | | 6.71 | -39 | -40 | -22 |
| | Inferior frontal gyrus (pars triangularis) | | | 6.65 | -42 | 26 | 2 |

| | | | | | | | |
|---|---|---|---|---|---|---|---|
| | Inferior frontal gyrus (pars triangularis) | | | 6.50 | -54 | 20 | 26 |
| | Inferior frontal gyrus (pars orbitalis) | | | 6.45 | -42 | 32 | -13 |
| | Inferior frontal gyrus (pars triangularis) | | | 6.34 | -57 | 26 | 14 |
| | Hippocampus | | | 6.22 | -21 | -10 | -13 |
| | Inferior occipital gyrus | | | 6.12 | -24 | -94 | -10 |
| | **R Occipital Lobe** | **176** | | | | | |
| | Lingual gyrus | | | 5.47 | 21 | -88 | -4 |
| | Middle occipital gyrus | | | 4.67 | 27 | -97 | 14 |
| | Middle occipital gyrus | | | 3.33 | 33 | -91 | 17 |
| | **L Frontal Lobe** | **250** | | | | | |
| | Medial superior frontal gyrus | | | 5.46 | -9 | 50 | 44 |
| | Superior frontal gyrus | | | 5.02 | -12 | 59 | 32 |
| | Medial superior frontal gyrus | | | 4.93 | -9 | 35 | 53 |
| | Superior frontal gyrus | | | 4.89 | -12 | 41 | 44 |
| | **R Temporal Lobe** | **109** | | | | | |
| | Middle temporal gyrus | | | 5.11 | 57 | -4 | -16 |
| **Non-social High Motivation > NJT** | **R Cerebellum** | **459** | 112 | | | | |
| | Crus II | | | 7.11 | 12 | -79 | -37 |
| | Crus VIII | | | 3.78 | 27 | -64 | -52 |

| | | | | | |
|---|---|---|---|---|---|
| **L Temporal Lobe** | **3079** | | | | |
| Medial temporal gyrus | | 7.07 | -48 | -40 | 2 |
| Inferior frontal gyrus (pars triangularis) | | 6.87 | -51 | 20 | 26 |
| Hippocampus | | 6.78 | -24 | -13 | -16 |
| Fusiform gyrus | | 6.67 | -42 | -43 | -22 |
| Inferior frontal gyrus (pars triangularis) | | 6.55 | -42 | 26 | -1 |
| Middle temporal gyrus | | 6.5 | -63 | -43 | 5 |
| Inferior frontal gyrus (pars triangularis) | | 6.39 | -42 | 11 | 26 |
| Inferior frontal gyrus (pars opercularis) | | 6.15 | -51 | 17 | 14 |
| **R Occipital Lobe** | **222** | | | | |
| Calcarine | | 6.41 | 18 | -94 | -1 |
| Middle occipital gyrus | | 5.48 | 27 | -97 | 14 |
| Middle occipital gyrus | | 3.81 | 33 | -91 | 17 |
| Middle occipital gyrus | | 3.48 | 42 | -88 | 2 |
| **L Occipital Lobe** | **384** | | | | |
| Middle occipital gyrus | | 6.01 | -15 | -91 | -7 |
| Inferior occipital gyrus | | 5.88 | -21 | -97 | -10 |
| Middle occipital gyrus | | 5.73 | -27 | -97 | 2 |
| Inferior occipital gyrus | | 5.02 | -33 | -91 | -10 |
| Superior occipital gyrus | | 4.52 | -15 | -100 | 17 |

| | | | | | | | |
|---|---|---|---|---|---|---|---|
| | Inferior occipital gyrus | | | 4.03 | -27 | -82 | -10 |
| | **R Posterior Temporal Cortex** | **112** | | | | | |
| | Caudate nucleus | | | 5.71 | 12 | 17 | 11 |
| | Putamen | | | 4.21 | 15 | 11 | -4 |
| | **L Frontal Lobe** | **192** | | | | | |
| | Superior medial frontal gyrus | | | 4.61 | -9 | 50 | 41 |
| | Supplementary motor area | | | 4.46 | -3 | 14 | 53 |
| | Superior frontal gyrus | | | 4.25 | -12 | 59 | 32 |
| | Superior medial frontal gyrus | | | 3.78 | -3 | 35 | 47 |
| **Non-social Low Motivation > NJT** | **L Frontal Lobe** | **2468** | 160 | | | | |
| | Inferior frontal gyrus (pars triangularis) | | | 7.34 | -51 | 23 | 23 |
| | Inferior frontal gyrus (pars orbitalis) | | | 7.18 | -42 | 26 | -4 |
| | Middle temporal gyrus | | | 7.06 | -48 | -40 | 2 |
| | Inferior frontal gyrus (pars opercularis) | | | 6.57 | -54 | 14 | 14 |
| | Inferior frontal gyrus (pars triangularis) | | | 6.55 | -51 | 26 | 14 |
| | Hippocampus | | | 6.39 | -27 | -16 | -16 |
| | Fusiform gyrus | | | 6.34 | -39 | -43 | -25 |
| | Inferior frontal gyrus (pars orbitalis) | | | 6.24 | -51 | 32 | -4 |
| | **R Cerebellum** | **437** | | | | | |
| | Crus II | | | 7.26 | 12 | -79 | -34 |
| | Crus II | | | 7.23 | 18 | -79 | -40 |

| | | | | | | | |
|---|---|---|---|---|---|---|---|
| | **R Occipital Lobe** | **160** | | | | | |
| | Calcarine | | | 6.14 | 18 | -94 | -1 |
| | Middle occipital gyrus | | | 4.82 | 36 | -94 | 2 |
| | **L Occipital Lobe** | **280** | | | | | |
| | Middle occipital gyrus | | | 5.72 | -24 | -97 | 2 |
| | Inferior occipital gyrus | | | 5.65 | -21 | -97 | -10 |
| | **L Frontal Lobe** | **250** | | | | | |
| | Superior medial frontal gyrus | | | 5.04 | -3 | 35 | 47 |
| | Superior medial frontal gyrus | | | 4.81 | -9 | 53 | 41 |
| | Superior medial frontal gyrus | | | 4.36 | -9 | 32 | 56 |
| | Supplementary motor area | | | 4.33 | -3 | 14 | 53 |
| | Superior frontal gyrus | | | 4.33 | -12 | 41 | 47 |
| **Social High Motivation > Non-social High motivation** | **L Medial Parietal Cortex** | **641** | 124 | | | | |
| | Precuneus | | | 5.82 | -6 | -55 | 32 |
| | **L Medial Temporal Cortex** | **490** | | | | | |
| | Middle temporal gyrus | | | 5.73 | -48 | -61 | 23 |
| | Angular gyrus | | | 5.65 | -45 | -73 | 32 |
| | **L Temporal Lobe** | **538** | | | | | |
| | Inferior temporal gyrus | | | 5.63 | -45 | 14 | -37 |

| | | | | | | | |
|---|---|---|---|---|---|---|---|
| | Inferior temporal gyrus | | | 5.36 | -48 | 2 | -31 |
| | Middle temporal gyrus | | | 5.18 | -57 | -10 | -16 |
| | Middle temporal pole | | | 4.69 | -39 | 14 | -31 |
| | Inferior frontal gyrus, orbital part | | | 4.62 | -36 | 20 | -13 |
| | Middle temporal gyrus | | | 4.17 | -60 | -31 | -4 |
| | Insular cortex | | | 3.95 | -27 | 14 | -19 |
| | **L Frontal Lobe** | **247** | | | | | |
| | Superior medial frontal gyrus | | | 5.03 | -6 | 59 | 11 |
| | Medial orbitofrontal cortex | | | 4.36 | -9 | 59 | -4 |
| | Superior frontal gyrus | | | 3.22 | -12 | 53 | 29 |
| | **R Frontal Lobe** | **124** | | | | | |
| | Superior medial frontal gyrus | | | 4.50 | 6 | 59 | 8 |
| | Superior medial frontal gyrus | | | 3.87 | 3 | 56 | 23 |
| | **R Temporal Occipital Cortex** | **241** | | | | | |
| | Middle temporal gyrus | | | 4.36 | 51 | -64 | 23 |
| | Middle occipital gyrus | | | 3.39 | 36 | -70 | 32 |
| **Social Low Motivation > Non-social Low Motivation** | **L Parietal Lobe** | **250** | 206 | | | | |
| | Angular gyrus | | | 5.57 | -42 | -73 | 41 |
| | Angular gyrus | | | 5.07 | -54 | -67 | 29 |
| | **L Medial Parietal Cortex** | **493** | | | | | |

| | | | | | | | |
|---|---|---|---|---|---|---|---|
| | Precuneus | | | 4.80 | -3 | -52 | 17 |
| | **R Parieto-Occipital Cortex** | **217** | | | | | |
| | Middle occipital gyrus | | | 4.72 | 45 | -67 | 29 |
| | Angular gyrus | | | 4.24 | 39 | -70 | 44 |
| | **Medial Frontal Cortex** | **206** | | | | | |
| | R Superior medial frontal gyrus | | | 4.14 | 9 | 62 | 8 |
| | R Rectus Gyrus | | | 3.97 | 3 | 53 | -19 |
| | L Rectus Gyrus | | | 3.82 | -6 | 44 | -28 |
| | L Medial orbitofrontal cortex | | | 3.52 | -3 | 50 | -13 |
| | R Rectus Gyrus | | | 3.36 | 9 | 41 | -25 |
| | L Medial orbitofrontal cortex | | | 3.30 | -3 | 65 | -7 |
| **Social High Motivation > Social Low Motivation** | **L Temporal Frontal Cortex** | **1185** | 1185 | | | | |
| | Middle temporal gyrus | | | 5.8 | -48 | -52 | 23 |
| | Middle temporal gyrus | | | 5.45 | -48 | -58 | 11 |
| | Superior temporal gyrus | | | 5.26 | -54 | -43 | 23 |
| | Inferior frontal gyrus (pars opercularis) | | | 4.88 | -54 | 11 | 20 |
| | Precentral gyrus | | | 4.62 | -51 | -4 | 29 |
| | Supramarginal gyrus | | | 4.52 | -54 | -49 | 35 |
| | Postcentral gyrus | | | 4.46 | -57 | -19 | 20 |
| | Inferior parietal lobule | | | 4.36 | -54 | -34 | 38 |

*Note*. Significant activation clusters in each condition minus number judgement contrast identified at *p* <.05, FWE-corrected, resulting in the specified extent threshold, after a cluster-defining threshold of *p* <.001 uncorrected. Table shows up to 8 local maxima more than 8mm apart in each cluster.

**Table S18.** *Significant activation clusters in whole-brain analysis within the anterior temporal lobe*

| Contrast | Cluster name and location of maximum | Cluster size (voxels) | Cluster extent threshold | Peak (Z) | MNI maxima coordinates (mm) | | |
|---|---|---|---|---|---|---|---|
| | | | | | x | y | z |
| SJT > NJT | L Middle temporal gyrus | **572** | 90 | 6.05 | -54 | -7 | -13 |
| | L Superior temporal pole | | | 5.98 | -51 | 17 | -16 |
| | L Superior temporal pole | | | 5.80 | -45 | 17 | -25 |
| | L Fusiform gyrus | | | 5.48 | -42 | -7 | -43 |
| | L Middle temporal pole | | | 4.90 | -33 | 14 | -40 |
| | L Inferior temporal gyrus | | | 4.62 | -36 | -7 | -34 |
| | R Superior temporal pole | **90** | | 5.73 | 48 | 14 | -19 |

*Note*. Significant activation clusters for words minus number judgement contrast identified at *p* <.05, FWE-corrected, resulting in the specified extent threshold, after a cluster-defining threshold of *p* <.001 uncorrected. Table shows up to 8 local maxima more than 8mm apart in each cluster. SJT = synonym judgement task; NJT = number judgement task

### ROI Analyses

**Table S19.** *Mean activation in all ROIs in the left and right hemisphere for each condition*

| ROIs | Word Condition | Mean Activation | SE |
|---|---|---|---|
| lMTG | NSH | 1.32 | 0.22 |
| lMTG | NSL | 1.21 | 0.21 |
| lMTG | SH | 1.77 | 0.19 |
| lMTG | SL | 1.71 | 0.24 |
| lTP | NSH | 0.31 | 0.17 |

| | | | |
|---|---|---|---|
| lTP | NSL | 0.35 | 0.17 |
| lTP | SH | 0.94 | 0.18 |
| lTP | SL | 0.73 | 0.20 |
| laSTG | NSH | 1.02 | 0.17 |
| laSTG | NSL | 0.87 | 0.19 |
| laSTG | SH | 1.23 | 0.19 |
| laSTG | SL | 1.07 | 0.21 |
| lvATL | NSH | 0.81 | 0.12 |
| lvATL | NSL | 0.73 | 0.12 |
| lvATL | SH | 0.90 | 0.11 |
| lvATL | SL | 0.81 | 0.14 |
| rMTG | NSH | 0.72 | 0.16 |
| rMTG | NSL | 0.60 | 0.19 |
| rMTG | SH | 1.04 | 0.17 |
| rMTG | SL | 1.06 | 0.17 |
| rTP | NSH | -0.01 | 0.14 |
| rTP | NSL | -0.10 | 0.11 |
| rTP | SH | 0.32 | 0.12 |
| rTP | SL | 0.18 | 0.14 |
| raSTG | NSH | 0.46 | 0.15 |
| raSTG | NSL | 0.29 | 0.16 |
| raSTG | SH | 0.63 | 0.13 |
| raSTG | SL | 0.41 | 0.15 |
| rvATL | NSH | 0.17 | 0.14 |
| rvATL | NSL | 0.01 | 0.11 |

| | | | |
|---|---|---|---|
| rvATL | SH | 0.04 | 0.12 |
| rvATL | SL | 0.09 | 0.11 |

*Note.* laSTG = left anterior superior temporal gyrus, lMTG = left middle temporal gyrus, lTP = left temporal pole, lvATL = left ventrolateral anterior temporal lobe, raSTG = right anterior superior temporal gyrus, rMTG = right middle temporal gyrus, rTP = right temporal pole, rvATL = right ventrolateral anterior temporal lobe. SH = Social High Motivation; SL = Social Low Motivation; NSH = Non-social High Motivation; NSL = Non-social Low Motivation.

**Table S20.** *Results of one-sample t-test in the ROI analyses in the left hemisphere*

| ROI | Condition | t-Statistic | P-value |
|---|---|---|---|
| LaMTG | NSH | 435 | *** |
| LaMTG | NSL | 428 | *** |
| LaMTG | SH | 464 | *** |
| LaMTG | SL | 450 | *** |
| LaSTG | NSH | 452 | *** |
| LaSTG | NSL | 417 | *** |
| LaSTG | SH | 441 | *** |
| LaSTG | SL | 425 | *** |
| LvlATL | NSH | 447 | *** |
| LvlATL | NSL | 441 | *** |
| LvlATL | SH | 462 | *** |
| LvlATL | SL | 448 | *** |
| lTP | NSH | 313 | 0.100 |
| lTP | NSL | 319 | 0.077 |
| lTP | SH | 433 | *** |
| lTP | SL | 392 | *** |

*Note.* laSTG = left anterior superior temporal gyrus, lMTG = left middle temporal gyrus, lTP = left temporal pole, lvATL = left ventrolateral anterior temporal lobe. SH = Social High Motivation; SL = Social Low Motivation; NSH = Non-social High Motivation; NSL = Non-social Low Motivation; *$p$<.05; **$p$<.01; ***$p$<.001

**Table S21.** *Results of three-way ANOVA in the ROI analyses in the left hemisphere*

| | Df | Sum Squared | Mean Squared | F value | P-value |
|---|---|---|---|---|---|
| ROI | 3 | 55.6 | 18.54 | 18.88 | *** |
| soc | 1 | 12.1 | 12.13 | 12.35 | *** |
| mot | 1 | 1.3 | 1.30 | 1.32 | 0.251 |

| | | | | | |
|---|---|---|---|---|---|
| ROI:soc | 3 | 3.8 | 1.28 | 1.31 | 0.272 |
| ROI:mot | 3 | 0.1 | 0.04 | 0.04 | 0.990 |
| soc:mot | 1 | 0.1 | 0.10 | 0.10 | 0.748 |
| ROI:soc:mot | 3 | 0.4 | 0.12 | 0.13 | 0.945 |
| Residuals | 464 | 455.6 | 0.98 | | |

*Note.* Df = degrees of freedom; soc = Socialness; mot = Motivation; *$p$<.05; **$p$<.01; ***$p$<.001

**Table S22.** *Results of pairwise Wilcoxon test between ROIs in the left hemisphere*

| Condition | | Group 1 | Group 2 | Statistic | P-value | Adjusted P-value |
|---|---|---|---|---|---|---|
| Socialness | Motivation | | | | | |
| social | highmot | laSTG | lMTG | 326 | 0.0 | 0.406 |
| social | highmot | laSTG | lTP | 532 | 0.230 | 1.000 |
| social | highmot | laSTG | lvATL | 545 | 0.164 | 0.984 |
| social | highmot | lMTG | lTP | 657 | 0.002 | * |
| social | highmot | lMTG | lvATL | 675 | 0.001 | ** |
| social | highmot | lTP | lvATL | 430 | 0.775 | 1.000 |
| social | lowmot | laSTG | lMTG | 315 | 0.046 | 0.277 |
| social | lowmot | laSTG | lTP | 532 | 0.230 | 1.000 |
| social | lowmot | laSTG | lvATL | 514 | 0.350 | 1.000 |
| social | lowmot | lMTG | lTP | 640 | 0.005 | * |
| social | lowmot | lMTG | lvATL | 648 | 0.003 | * |
| social | lowmot | lTP | lvATL | 422 | 0.686 | 1.000 |
| non-social | highmot | laSTG | lMTG | 385 | 0.343 | 1.000 |
| non-social | highmot | laSTG | lTP | 623 | 0.010 | 0.060 |
| non-social | highmot | laSTG | lvATL | 477 | 0.697 | 1.000 |
| non-social | highmot | lMTG | lTP | 659 | 0.002 | * |
| non-social | highmot | lMTG | lvATL | 561 | 0.103 | 0.618 |
| non-social | highmot | lTP | lvATL | 298 | 0.024 | 0.146 |
| non-social | lowmot | laSTG | lMTG | 363 | 0.203 | 1.000 |
| non-social | lowmot | laSTG | lTP | 568 | 0.082 | 0.494 |
| non-social | lowmot | laSTG | lvATL | 459 | 0.901 | 1.000 |
| non-social | lowmot | lMTG | lTP | 645 | 0.004 | * |
| non-social | lowmot | lMTG | lvATL | 585 | 0.046 | 0.277 |
| non-social | lowmot | lTP | lvATL | 343 | 0.116 | 0.696 |

*Note.* laSTG = left anterior superior temporal gyrus ; lMTG = left middle temporal gyrus; lTP = left temporal pole; lvATL = left ventrolateral anterior temporal lobe; lowmot = low motivation; highmot = high motivation; *$p$<.05; **$p$<.01; ***$p$<.001

**Table S23.** *Results of pairwise Wilcoxon test assessing the main effects of socialness across all ROIs in the left hemisphere*

| ROI | Group 1 | Group 2 | Statistic | P-value | Adjusted P-value |
|---|---|---|---|---|---|
| laSTG | social | non-social | 2178 | * | * |
| lTP | social | non-social | 2293 | ** | ** |
| lvATL | social | non-social | 2031 | 0.226 | 0.226 |
| lMTG | social | non-social | 1926 | 0.510 | 0.510 |

*Note.* laSTG = left anterior superior temporal gyrus ; lMTG = left middle temporal gyrus; lTP = left temporal pole; lvATL = left ventrolateral anterior temporal lobe; *$p$<.05; **$p$<.01; ***$p$<.001

**Table S24.** *Results of pairwise Wilcoxon test assessing the main effects of motivation across all ROIs in the left hemisphere*

| ROI | Group 1 | Group 2 | Statistic | P-value | Adjusted P-value |
|---|---|---|---|---|---|
| laSTG | highmot | lowmot | 1872 | 0.707 | 0.707 |
| lTP | highmot | lowmot | 1875 | 0.696 | 0.696 |
| lvATL | highmot | lowmot | 1973 | 0.365 | 0.365 |
| lMTG | highmot | lowmot | 1970 | 0.374 | 0.374 |

*Note.* laSTG = left anterior superior temporal gyrus ; lMTG = left middle temporal gyrus; lTP = left temporal pole; lvATL = left ventrolateral anterior temporal lobe; highmot = high motivation; lowmot = low motivation

**Table S25.** *Results of simple effects analyses for socialness across levels of motivation in all ROIs in the left hemisphere*

| Contrast | ROI | Motivation | Estimate | SE | Df | T Ratio | P-value |
|---|---|---|---|---|---|---|---|
| social - non-social | laSTG | highmot | 0.21 | 0.26 | 464 | 0.84 | 0.401 |
| social - non-social | lMTG | highmot | 0.45 | 0.26 | 464 | 1.76 | 0.078 |
| social - non-social | lTP | highmot | 0.63 | 0.26 | 464 | 2.46 | * |
| social - non-social | lvATL | highmot | 0.09 | 0.26 | 464 | 0.36 | 0.721 |
| social - non-social | laSTG | lowmot | 0.20 | 0.26 | 464 | 0.80 | 0.436 |
| social - non-social | lMTG | lowmot | 0.50 | 0.26 | 464 | 1.96 | 0.052 |
| social - non-social | lTP | lowmot | 0.39 | 0.26 | 464 | 1.50 | 0.133 |
| social - non-social | lvATL | lowmot | 0.07 | 0.26 | 464 | 0.29 | 0.775 |

*Note.* laSTG = left anterior superior temporal gyrus ; lMTG = left middle temporal gyrus; lTP = left temporal pole; lvATL = left ventrolateral anterior temporal lobe; lowmot = low motivation; highmot = high motivation; SE = standard error; Df = degrees of freedom; *p<.05; **p<.01; ***p<.001

**Table S26.** *Results of simple effects analyses for motivation across levels of socialness in all ROIs in the left hemisphere*

| Contrast | ROI | Socialness | Estimate | SE | Df | T Ratio | P-value |
|---|---|---|---|---|---|---|---|
| highmot - lowmot | laSTG | social | 0.16 | 0.26 | 464 | 0.64 | 0.520 |
| highmot - lowmot | lMTG | social | 0.07 | 0.26 | 464 | 0.26 | 0.795 |
| highmot - lowmot | lTP | social | 0.21 | 0.26 | 464 | 0.81 | 0.421 |
| highmot - lowmot | lvATL | social | 0.10 | 0.26 | 464 | 0.37 | 0.710 |
| highmot - lowmot | laSTG | non-social | 0.15 | 0.26 | 464 | 0.58 | 0.559 |
| highmot - lowmot | lMTG | non-social | 0.11 | 0.26 | 464 | 0.44 | 0.660 |
| highmot - lowmot | lTP | non-social | -0.04 | 0.26 | 464 | -0.15 | 0.879 |
| highmot - lowmot | lvATL | non-social | 0.08 | 0.26 | 464 | 0.30 | 0.764 |

*Note.* laSTG = left anterior superior temporal gyrus ; lMTG = left middle temporal gyrus; lTP = left temporal pole; lvATL = left ventrolateral anterior temporal lobe; lowmot = low motivation; highmot = high motivation; SE = standard error; Df = degrees of freedom

**Table S27.** *Results of one-sample t-tests in the ROI analyses in the right hemisphere*

| ROI | Condition | t-Statistic | P-value |
|---|---|---|---|
| RaMTG | NSH | 402 | *** |
| RaMTG | NSL | 374 | ** |
| RaMTG | SH | 440 | *** |
| RaMTG | SL | 437 | *** |
| RaSTG | NSH | 363 | ** |
| RaSTG | NSL | 316 | 0.088 |
| RaSTG | SH | 421 | *** |
| RaSTG | SL | 346 | * |
| RvlATL | NSH | 271 | 0.440 |

| | | | |
|---|---|---|---|
| RvlATL | NSL | 233 | 1.000 |
| RvlATL | SH | 266 | 0.503 |
| RvlATL | SL | 274 | 0.404 |
| rTP | NSH | 223 | 0.855 |
| rTP | NSL | 192 | 0.416 |
| rTP | SH | 347 | * |
| rTP | SL | 272 | 0.428 |

*Note.* raSTG = right anterior superior temporal gyrus, rMTG = right middle temporal gyrus, rTP = right temporal pole, rvATL = right ventrolateral anterior temporal lobe. SH = Social High Motivation; SL = Social Low Motivation; NSH = Non-social High Motivation; NSL = Non-social Low Motivation; *$p$<.05; **$p$<.01; ***$p$<.001

**Table S28.** *Results of three-way ANOVA in the ROI analyses in the right hemisphere*

| | Df | Sum Squared | Mean Squared | F value | P-value |
|---|---|---|---|---|---|
| ROI | 3 | 48.34 | 16.113 | 26.029 | *** |
| soc | 1 | 5.03 | 5.027 | 8.121 | ** |
| mot | 1 | 1.28 | 1.281 | 2.07 | 0.151 |
| ROI:soc | 3 | 2.9 | 0.968 | 1.563 | 0.198 |
| ROI:mot | 3 | 0.41 | 0.138 | 0.223 | 0.881 |
| soc:mot | 1 | 0.1 | 0.104 | 0.167 | 0.683 |
| ROI:soc:mot | 3 | 0.41 | 0.136 | 0.22 | 0.882 |
| Residuals | 464 | 287.23 | 0.619 | | |

*Note.* Df = degrees of freedom; soc = Socialness; aud = Auditory Embodiment; *$p$<.05; **$p$<.01; ***$p$<.001

**Table S29.** *Results of pairwise Wilcoxon test between ROIs in the right hemisphere*

| Condition | | Group 1 | Group 2 | Statistic | P-value | Adjusted P-value |
|---|---|---|---|---|---|---|
| soc | mot | | | | | |
| social | highmot | raSTG | rMTG | 336 | 0.094 | 0.561 |
| social | highmot | raSTG | rTP | 552 | 0.134 | 0.804 |
| social | highmot | raSTG | rvATL | 644 | 0.004 | * |
| social | highmot | rMTG | rTP | 662 | 0.001 | ** |
| social | highmot | rMTG | rvATL | 721 | 0.000 | *** |
| social | highmot | rTP | rvATL | 535 | 0.213 | 1.000 |
| social | lowmot | raSTG | rMTG | 265 | 0.006 | * |
| social | lowmot | raSTG | rTP | 545 | 0.164 | 0.984 |

| Condition soc | Condition mot | Group 1 | Group 2 | Statistic | P-value | Adjusted P-value |
|---|---|---|---|---|---|---|
| social | lowmot | raSTG | rvATL | 553 | 0.130 | 0.780 |
| social | lowmot | rMTG | rTP | 706 | 0.000 | *** |
| social | lowmot | rMTG | rvATL | 732 | 0.000 | *** |
| social | lowmot | rTP | rvATL | 442 | 0.912 | 1.000 |
| non-social | highmot | raSTG | rMTG | 370 | 0.242 | 1.000 |
| non-social | highmot | raSTG | rTP | 593 | 0.034 | 0.207 |
| non-social | highmot | raSTG | rvATL | 555 | 0.123 | 0.738 |
| non-social | highmot | rMTG | rTP | 670 | 0.001 | ** |
| non-social | highmot | rMTG | rvATL | 631 | 0.007 | * |
| non-social | highmot | rTP | rvATL | 397 | 0.440 | 1.000 |
| non-social | lowmot | raSTG | rMTG | 363 | 0.203 | 1.000 |
| non-social | lowmot | raSTG | rTP | 578 | 0.059 | 0.355 |
| non-social | lowmot | raSTG | rvATL | 549 | 0.146 | 0.876 |
| non-social | lowmot | rMTG | rTP | 645 | 0.004 | * |
| non-social | lowmot | rMTG | rvATL | 627 | 0.008 | 0.050 |
| non-social | lowmot | rTP | rvATL | 405 | 0.513 | 1.000 |

*Note.* raSTG = right anterior superior temporal gyrus ; rMTG = right middle temporal gyrus; rTP = right temporal pole; rvATL = right ventrolateral anterior temporal lobe; lowmot = low motivation; highmot = high motivation; **p*<.05; ***p*<.01; ****p*<.001

**Table S30.** *Results of pairwise Wilcoxon test assessing the main effects of socialness across all ROIs in the right hemisphere*

| ROI | Group 1 | Group 2 | Statistic | P-value | Adjusted P-value |
|---|---|---|---|---|---|
| raSTG | social | non-social | 2203 | * | * |
| rTP | social | non-social | 2202 | * | * |
| rvATL | social | non-social | 1995 | 0.307 | 0.307 |

| | | | | | |
|---|---|---|---|---|---|
| rMTG | social | non-social | 1884 | 0.661 | 0.661 |

*Note.* raSTG = right anterior superior temporal gyrus ; rMTG = right middle temporal gyrus; rTP = right temporal pole; rvATL = right ventrolateral anterior temporal lobe; *$p$<.05; **$p$<.01; ***$p$<.001

**Table S31.** *Results of pairwise Wilcoxon test assessing the main effects of motivation across all ROIs in the right hemisphere*

| ROI | Group 1 | Group 2 | Statistic | P-value | Adjusted P-value |
|---|---|---|---|---|---|
| raSTG | highmot | lowmot | 1818 | 0.927 | 0.927 |
| rTP | highmot | lowmot | 2023 | 0.243 | 0.243 |
| rvATL | highmot | lowmot | 2035 | 0.218 | 0.218 |
| rMTG | highmot | lowmot | 1836 | 0.852 | 0.852 |

*Note.* raSTG = right anterior superior temporal gyrus ; rMTG = right middle temporal gyrus; rTP = right temporal pole; rvATL = right ventrolateral anterior temporal lobe; highmot = high motivation; lowmot = low motivation

**Table S32.** *Results of simple effects analyses for socialness across levels of motivation in all ROIs in the right hemisphere*

| Contrast | ROI | Motivation | Estimate | SE | Df | T Ratio | P-value |
|---|---|---|---|---|---|---|---|
| social - non-social | raSTG | highmot | 0.18 | 0.20 | 464 | 0.87 | 0.386 |
| social - non-social | rMTG | highmot | 0.32 | 0.20 | 464 | 1.58 | 0.115 |
| social - non-social | rTP | highmot | 0.33 | 0.20 | 464 | 1.62 | 0.107 |
| social - non-social | rvATL | highmot | -0.12 | 0.20 | 464 | -0.61 | 0.542 |
| social - non-social | raSTG | lowmot | 0.12 | 0.20 | 464 | 0.57 | 0.566 |
| social - non-social | rMTG | lowmot | 0.46 | 0.20 | 464 | 2.24 | * |
| social - non-social | rTP | lowmot | 0.28 | 0.20 | 464 | 1.37 | 0.171 |
| social - non-social | rvATL | lowmot | 0.09 | 0.20 | 464 | 0.42 | 0.673 |

*Note.* raSTG = right anterior superior temporal gyrus ; rMTG = right middle temporal gyrus; rTP = right temporal pole; rvATL = right ventrolateral anterior temporal lobe; lowmot = low motivation; highmot = high motivation; SE = standard error; Df = degrees of freedom; *$p$<.05; **$p$<.01; ***$p$<.001

**Table S33.** *Results of simple effects analyses for motivation across levels of socialness in all ROIs in the right hemisphere*

| Contrast | ROI | Socialness | Estimate | SE | Df | T Ratio | P-value |
|---|---|---|---|---|---|---|---|
| highmot - lowmot | raSTG | social | 0.22 | 0.20 | 464 | 1.10 | 0.273 |
| highmot - lowmot | rMTG | social | -0.02 | 0.20 | 464 | -0.09 | 0.925 |
| highmot - lowmot | rTP | social | 0.14 | 0.20 | 464 | 0.71 | 0.480 |
| highmot - lowmot | rvATL | social | -0.05 | 0.20 | 464 | -0.25 | 0.801 |
| highmot - lowmot | raSTG | non-social | 0.16 | 0.20 | 464 | 0.80 | 0.422 |
| highmot - lowmot | rMTG | non-social | 0.12 | 0.20 | 464 | 0.57 | 0.569 |
| highmot - lowmot | rTP | non-social | 0.09 | 0.20 | 464 | 0.46 | 0.646 |
| highmot - lowmot | rvATL | non-social | 0.16 | 0.20 | 464 | 0.78 | 0.436 |

**Table S34.** *Wilcoxon pairwise t-test comparing mean activation within the same ROI and condition across hemispheres*

| Group 1 | Group 2 | Statistic | P-value | Adjusted P-value |
|---|---|---|---|---|
| lTP_SH | rTP_SH | 624 | 0.010 | 0.154 |
| lTP _NSH | rTP _NSH | 540 | 0.187 | 1.000 |
| lTP _SL | rTP _SL | 612 | 0.016 | 0.259 |
| lTP _NSL | rTP _NSL | 584 | 0.048 | 0.766 |
| lvATL_SH | rvATL_SH | 761 | 0.000 | *** |
| lvATL_NSH | rvATL_NSH | 673 | 0.001 | * |
| lvATL_SL | rvATL_SL | 690 | 0.000 | ** |
| lvATL_NSL | rvATL_NSL | 724 | 0.000 | *** |
| lMTG_SH | rMTG_SH | 628 | 0.008 | 0.128 |
| lMTG_NSH | rMTG_NSH | 576 | 0.063 | 1.000 |
| lMTG_SL | rMTG_SL | 579 | 0.057 | 0.914 |
| lMTG_NSL | rMTG_NSL | 588 | 0.042 | 0.664 |
| laSTG_SH | raSTG_SH | 624 | 0.010 | 0.154 |
| laSTG_NSH | raSTG_NSH | 579 | 0.057 | 0.914 |
| laSTG_SL | raSTG_SL | 602 | 0.024 | 0.389 |
| laSTG_NSL | raSTG_NSL | 577 | 0.061 | 0.978 |

*Note.* laSTG = left anterior superior temporal gyrus, lMTG = left middle temporal gyrus, lTP = left temporal pole, lvATL = left ventrolateral anterior temporal lobe, raSTG = right anterior superior temporal gyrus, rMTG = right middle temporal gyrus, rTP = right temporal pole, rvATL = right ventrolateral anterior temporal lobe. SH = Social High Motivation; SL = Social Low Motivation; NSH = Non-social High Motivation; NSL = Non-social Low Motivation. *** <.001; ** <.01; * <.05

We extracted the tSNR values from each ROI by dividing the mean intensity in each voxel by its standard deviation across the four runs for each participant. We obtained a mean tSNR value per ROI for each participant. There were statistically significant differences in tSNR between right aSTG and aMTG (*p* = 0.003), right TP and MTG (*p* = 0.001), right TP and left MTG (*p* <.001), right vATL and MTG (*p* = 0.031), left TP and left MTG (*p* = 0.013), where the aMTG had a higher tSNR value (see Table S35 and S36).

**Table S35.** *tSNR values in each ROI*

| ROI | Mean tSNR |
|---|---|
| raSTG | 229.9035 |
| rTP | 208.7219 |
| rMTG | 276.5087 |
| rvATL | 238.1388 |
| laSTG | 241.6746 |
| lTP | 235.5663 |
| lMTG | 288.1422 |
| lvATL | 247.3211 |

*Note.* laSTG = left anterior superior temporal gyrus, lMTG = left middle temporal gyrus, lTP = left temporal pole, lvATL = left ventrolateral anterior temporal lobe, raSTG = right anterior superior temporal gyrus, rMTG = right middle temporal gyrus, rTP = right temporal pole, rvATL = right ventrolateral anterior temporal lobe.

**Table S36.** *Wilcoxon pairwise test comparing tSNR values across ROIs*

| Group 1 | Group 2 | Statistic | P-value | Adjusted P-value |
|---|---|---|---|---|
| raSTG | lMTG | 194 | *** | ** |
| rTP | rMTG | 183 | *** | ** |
| rTP | lMTG | 144 | *** | *** |
| rvATL | lMTG | 233 | ** | * |
| lTP | lMTG | 218 | *** | * |

*Note.* Table include only significant differences. lMTG = left middle temporal gyrus, lTP = left temporal pole, raSTG = right anterior superior temporal gyrus, rMTG = right middle temporal gyrus, rTP = right temporal pole, rvATL = right ventrolateral anterior temporal lobe. *** <.001; ** <.01; * <.05

### Section S5: Additional Multivariate Analyses Results

#### *ROI-level*

***Pattern Classification.*** We conducted a pattern classification analysis to assess if the local activation patterns within the pre-defined ROIs encode socialness. When training and testing within high motivation, we found above chance accuracy in the left TP. When training and testing within low motivation, we found above chance accuracy in the left MTG and vlATL. The robustness analyses that included only accurate trials showed the same pattern, with the exception that we found additional activation in the left TP for decoding socialness within low motivation (see Figure S4 A & B).

We conducted a pattern classification analysis to assess if the local activation patterns within the pre-defined ROIs encode motivation. When training and testing within social concepts we also found above chance accuracy in the left TP and right aSTG. When training and testing within non-social concepts we found no above chance accuracy (see Figure S4 C & D). In the robustness analyses, there was no significantly above chance decoding.

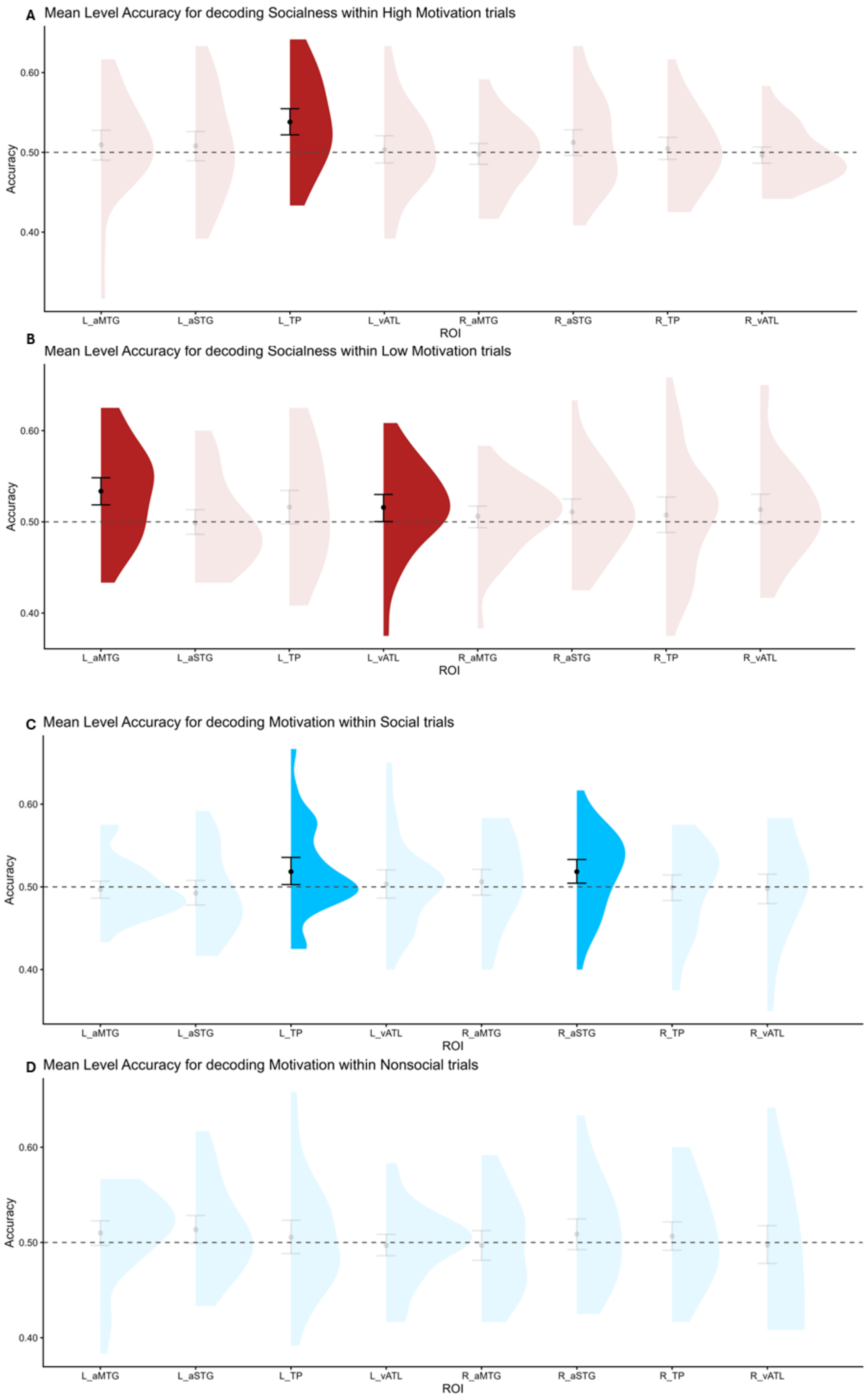
A Mean Level Accuracy for decoding Socialness within High Motivation trials
Accuracy
0.60
0.50
0.40
L_aMTG
L_aSTG
L_TP
L_vATL
R_aMTG
R_aSTG
R_TP
R_vATL
ROI
B
Mean Level Accuracy for decoding Socialness within Low Motivation trials
Accuracy
0.60
0.50
0.40
L_aMTG
L_aSTG
L_TP
L_vATL
R_aMTG
R_aSTG
R_TP
R_vATL
ROI
C Mean Level Accuracy for decoding Motivation within Social trials
Accuracy
0.60
0.50
0.40
L_aMTG
L_aSTG
L_TP
L_vATL
R_aMTG
R_aSTG
R_TP
R_vATL
ROI
D Mean Level Accuracy for decoding Motivation within Nonsocial trials
Accuracy
0.60
0.50
0.40
L_aMTG
L_aSTG
L_TP
L_vATL
R_aMTG
R_aSTG
R_TP
R_vATL
ROI

***Figure S4.*** *Accuracy levels for decoding A) Socialness within High Motivation trials, B) Socialness within Low Motivation trials, C) Motivation within Social trials, D) Motivation within Non-social trials. Dotted line represents chance-level (50%) accuracy. Error bars represent 90% confidence intervals bootstrapped around the mean. Transparency indicates non-significance in one-sample t-test.*

***Cross-decoding.*** When decoding social concepts across levels of association with motivation, we found significantly above chance accuracy in the bilateral vlATL and left aMTG when training on high motivation trials and testing on low motivation trials (see Figure 5A). Additionally when training on low motivation trials and testing on high motivation trials, we found significantly above chance decoding accuracy in the left vlATL and aMTG (see Figure 5B). In the robustness analysis, we found significantly above chance decoding accuracy in the left aMTG and TP when training on high motivation trials and testing on low motivation trials, and only in the left aMTG and vlATL when collapsing across training directions.

When decoding motivation across levels of association with socialness, accuracy was not significantly above chance. In the robustness analysis, we found additional effects in the left TP (collapsed) and in the bilateral TP (train non-social test social).

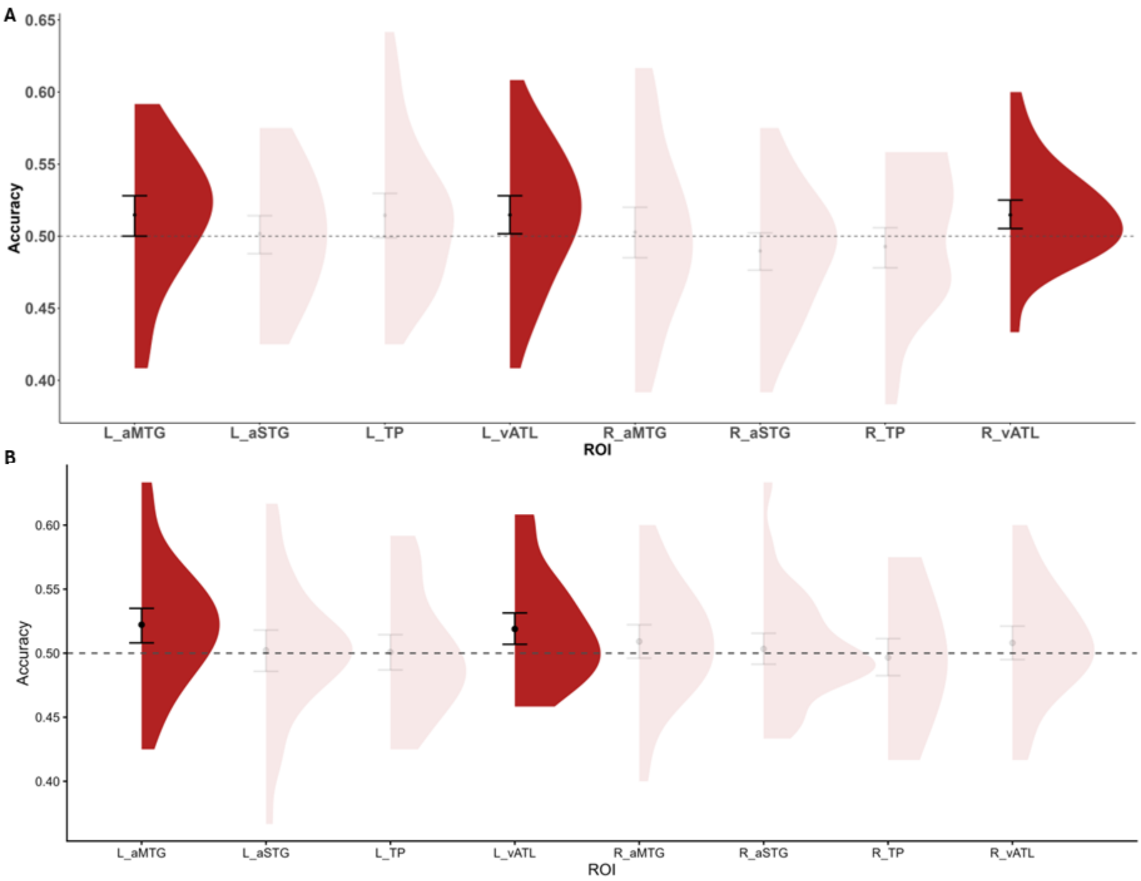

***Figure S5.*** *Accuracy levels for cross-decoding A) Socialness when training in high motivation trials and testing in low motivation trials, B) Socialness when training in low motivation trials and testing in high motivation trials. Dotted line represents chance-level (50%) accuracy. Error bars represent 90% confidence intervals bootstrapped around the mean. Transparency indicates non-significance in one-sample t-test.*

#### *ATL-restricted Searchlight*

***Pattern Classification.*** We conducted a searchlight analysis restricted to the ATL to examine local patterns of activation sensitive to socialness outside of the a priori ROIs. In addition to the above chance decoding accuracy in regions corresponding to the pre-defined ROIs, we found a cluster in the left superior TP when testing and training within low motivation trials only (see Figure S6 A). This was consistent in the robustness analysis. There was no significant above chance decoding of motivation across levels of association with socialness.

***Cross-decoding.*** We examined decoding of social concepts across levels of association with motivation outside of the a priori ROIs. A group of voxels in the left ITG showed above chance decoding accuracy when decoding in both directions (see Figure S6 B & C). In the robustness analyses, socialness was decoded in a smaller group of voxels in the left ITG with above chance accuracy only when training on low motivation trials and testing on high motivation trials.

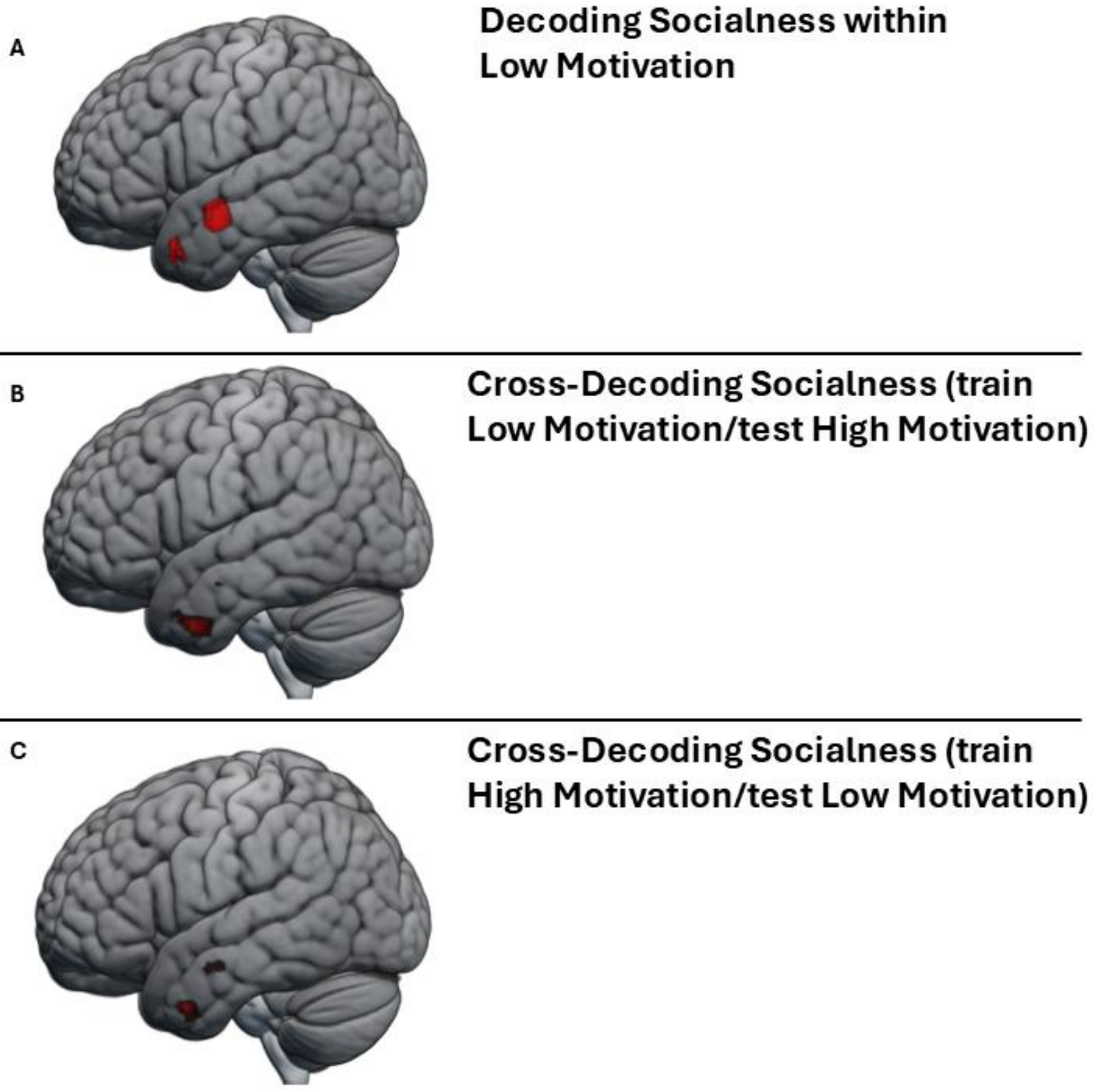

***Figure S6.*** ATL regions associated with above chance classification accuracy in the searchlight MVPA for A) decoding socialness within low motivation trials, B) cross-decoding socialness when training in high motivation trials and testing in low motivation trials, C) cross-decoding socialness when training in low motivation trials and testing in high motivation trials.

**Table S37.** *Significant activation clusters in the searchlight decoding analysis of socialness across all trials*

| Cluster ID | X | Y | Z | Peak Stat | Cluster Size (mm3) |
|---|---|---|---|---|---|
| **L Middle temporal gyrus** | -63 | -7 | -19 | 5.03 | 12285 |
| **L Inferior temporal gyrus** | -45 | 2 | -37 | 4.78 | |
| **L Middle temporal pole** | -45 | 14 | -34 | 4.17 | |
| **L Superior temporal pole** | -42 | 17 | -19 | 3.83 | |
| **R Superior temporal pole** | 48 | 20 | -19 | 3.29 | 432 |
| **R Middle temporal gyrus** | 60 | -7 | -25 | 3.25 | 702 |
| **R Superior temporal gyrus** | 60 | 2 | -7 | 3.17 | 351 |

**Table S38.** *Significant activation clusters in the searchlight decoding analysis of socialness across low motivation trials*

| Cluster ID | X | Y | Z | Peak Stat | Cluster Size (mm3) |
|---|---|---|---|---|---|
| **L Middle temporal gyrus** | -63 | -4 | -16 | 6.20 | 2376 |
| **L** | -39 | 14 | -34 | 3.95 | 1323 |
| | -48 | 14 | -34 | 3.86 | |
| **R Inferior temporal gyrus** | 54 | 17 | -16 | 3.40 | 27 |

**Table S39.** *Significant activation clusters in the searchlight decoding analysis of motivation across all trials*

| Cluster ID | X | Y | Z | Peak Stat | Cluster Size (mm3) |
|---|---|---|---|---|---|
| **L Inferior temporal gyrus** | -54 | 2 | -34 | 3.92 | 297 |

| Cluster ID | X | Y | Z | Peak Stat | Cluster Size (mm3) |
|---|---|---|---|---|---|
| | -48 | 11 | -28 | 3.46 | 135 |

**Table S40.** *Significant activation clusters in the searchlight cross-decoding analysis of socialness across motivation (train high test low)*

| Cluster ID | X | Y | Z | Peak Stat | Cluster Size (mm3) |
|---|---|---|---|---|---|
| **L Inferior temporal gyrus** | -45 | 2 | -37 | 4.29 | 891 |
| **L Middle temporal gyrus** | -63 | -1 | -19 | 3.96 | 351 |
| **L Inferior temporal gyrus** | -63 | -10 | -19 | 3.80 | |

**Table S41.** *Significant activation clusters in the searchlight cross-decoding analysis of socialness across motivation (train low test high)*

| Cluster ID | X | Y | Z | Peak Stat | Cluster Size (mm3) |
|---|---|---|---|---|---|
| **L Inferior temporal gyrus** | -48 | 2 | -37 | 4.62 | 2943 |
| **L Inferior temporal gyrus** | -48 | 14 | -34 | 3.81 | |
| | -39 | 17 | -34 | 3.49 | |
| | -36 | 8 | -37 | 3.46 | |
| **L Middle temporal gyrus** | -57 | -7 | -19 | 3.83 | 405 |
| **L Inferior temporal gyrus** | -48 | -10 | -13 | 3.48 | 54 |
